# C/EBPα confers dependence to fatty acid anabolic pathways and vulnerability to lipid oxidative stress in FLT3-mutant leukemia

**DOI:** 10.1101/2022.04.15.488522

**Authors:** Marie Sabatier, Rudy Birsen, Laura Lauture, Jonas Dehairs, Paolo Angelino, Sarah Mouche, Maël Heiblig, Emeline Boet, Ambrine Sahal, Estelle Saland, Thomas Farge, Guillaume Cognet, Federico Simonetta, Corentin Pignon, Antoine Graffeuil, Céline Mazzotti, Hervé Avet-Loiseau, Océane Delos, Justine Bertrand-Michel, Amélie Chedru, François Vergez, Véronique Mansat-De Mas, Sarah Bertoli, Suzanne Tavitian, Muriel Picard, Christian Récher, Olivier Kosmider, Pierre Sujobert, Benoit Colsch, Carine Joffre, Lucille Stuani, Johannes V. Swinnen, Hervé Guillou, Petros Tsantoulis, Clément Larrue, Didier Bouscary, Jérôme Tamburini, Jean-Emmanuel Sarry

## Abstract

While transcription factor C/AAT-enhancer binding protein α (C/EBPα) is critical for normal and leukemic differentiation, its role on cell and metabolic homeostasis is largely unknown in cancer. Here, multi-omics analyses uncovered a coordinated activation of C/EBPα and Fms-like tyrosine kinase 3 (FLT3) that increased lipid anabolism *in vivo* and in patients with *FLT3*-mutant acute myeloid leukemia (AML). Mechanistically, C/EBPα regulated FASN-SCD axis to promote fatty acid (FA) biosynthesis and desaturation. We further demonstrated that FLT3 or C/EBPα inactivation decreased mono-unsaturated FAs incorporation to membrane phospholipids through SCD downregulation. Consequently, SCD inhibition enhanced susceptibility to lipid redox stress. Moreover, this C/EBPα-dependent adaptation of FA homeostasis was exploited by combining FLT3 and glutathione peroxidase 4 (GPX4) inhibition to trigger lipid oxidative stress, enhancing ferroptotic death of *FLT3*-mutant AML cells. Altogether, our study reveals a C/EBPα function in lipid homeostasis and adaptation to redox stress, and a previously unreported vulnerability of FLT3-mutant AML with promising therapeutic application.

**SIGNIFICANCE:** The transcription factor C/EBPα is as a master regulator of normal and leukemic myeloid differentiation. Here, we discovered that C/EBPα regulates fatty acid biosynthesis and metabolic adaptation to redox imbalance in leukemic cells. This confers a vulnerability to lipid oxidative stress to *FLT3*-mutant cells and supports novel therapeutic opportunities for patients.

## INTRODUCTION

Acute myeloid leukemia (AML) are aggressive hematological malignancies characterized by an aberrant proliferation of myeloid progenitors leading to an impairment of normal hematopoiesis. *FLT3* mutations are detected in 30% of AML cases at diagnosis and are associated with adverse prognostic due to frequent relapse after intensive chemotherapy^1–4^. Most *FLT3* mutations are internal tandem duplication (ITD) in the juxta-membrane domain, while tyrosine kinase domain (TKD) point mutations are less frequently detected^1^. Both ITD and TKD mutations result in a constitutive activation of FLT3-dependent signaling pathways, supporting cell proliferation and survival through enhanced glucose, amino acid and redox metabolism^5–9^. Recently, FLT3 inhibitors (FLT3i) such as midostaurin or gilteritinib (GILT) were shown to improve therapeutic response in *FLT3*-mutated AML patients when combined to intensive chemotherapy frontline, or used as a monotherapy in the relapse setting^10,11^. However, resistance to FLT3i commonly appears, involving paracrine signaling and metabolic sensing from bone marrow microenvironment^12–17^, emergence of drug-resistant secondary *FLT3* mutations or clonal escape through activation of alternative pathways, as seen with RAS-activating mutations^17–20^.

C/AAT-enhancer binding protein α (C/EBPα, encoded by the *CEBPA* gene) is a key transcription factor of myeloid differentiation^21^. Different *CEBPA* mutations are detected in 5-9% of AML cases at diagnosis, with prognostic implications^1,22^. Moreover, *CEBPA* expression is downregulated in AML subtypes such as those harboring *RUNX1*–*CBF2T1* fusion gene, which participate to differentiation block, highlighting the importance of C/EBPα in leukemogenesis^23^. Beside these hematopoietic functions, C/EBPα is a critical regulator of transcriptional programs related to glucose and lipid metabolism in normal adipocytes and hepatocytes^24–26^. However, the role of C/EBPα in cancer and leukemic metabolism has not been studied so far.

To investigate the C/EBPα-induced metabolic dependencies of highly metabolically active AML cells^5–9^, we performed multi-omics and functional approaches *in vitro, in vivo* using *FLT3*-mutant AML patient-derived xenograft (PDX) models, and in primary specimens from patients treated with GILT. We uncovered a novel role of C/EBPα on lipid biosynthesis downstream of FLT3. Specifically, we showed that FLT3 and C/EBPα are coordinately activated in AML to induce the expression of Fatty Acid Synthase (*FASN*), Steroyl-CoA Desaturase (*SCD*) and Fatty Acid Desaturase 2 (*FADS2*), promoting *de novo* biosynthesis of cytoprotective mono-unsaturated fatty acids (MUFA) and generation of highly unsaturated FA from the essential poly-unsaturated FA (PUFA) linoleic acid precursor, respectively. Accordingly, FLT3 inhibition led to C/EBPα depletion, impacting the cellular lipid composition and FA distribution. This sensitized leukemic cells to lipid peroxidation and induced their vulnerability to ferroptotic cell death. Finally, we demonstrated that ferroptosis induction synergized with GILT *ex vivo* and *in vivo*, indicating new therapeutic approaches for patients with *FLT3*-mutant AML.

## RESULTS

### *FLT3*-mutant AML cells exhibit increased lipid biosynthesis

To investigate changes in metabolic pathways of FLT3-activated AML cells, we analyzed differentially expressed genes (DEG) in *FLT3*-ITD-mutated MOLM-14 and MV4-11 cell lines, after FLT3i treatment or FLT3 depletion by RNA interference, and we defined a FLT3-ITD_UP cell line signature composed of 299 downregulated genes common to these conditions compared to the control cell lines (**Figure S1A-D** and **Table S1**). This functionally-defined signature was enriched in genes related to cell cycle and DNA/RNA processing, amino acid and redox metabolism pathways as previously described^5,27–30^, and to lipid metabolism including cholesterol and fatty acid (FA) biosynthesis (**Figure S1E** and **Table S2**). The same signatures were enriched in transcriptomes of *FLT3*-ITD expressing cells from a MLL-AF9-driven murine AML model (GSE1639329^5^) (**Figure S1F** and **Table S2**).

To generate *FLT3*-mutant models more relevant to patients with AML, we amplified primary leukemic cells from three different AML patients through serial transplantations in NSG mice (**Figure 1A** and **Table S3**). Using this approach, we enriched samples in leukemic blasts having high variant allele frequency of *FLT3* mutation (**Figure 1A**). After *in vivo* expansion in third recipients, we performed a transcriptomic analysis on human leukemic cells incubated *ex vivo* with FLT3i (**Figure 1A**). Using Gene Set Enrichment Analysis (GSEA), we observed a depletion of gene signatures related to glucose, amino acids, mitochondria and lipid metabolism in FLT3i-treated compared to control cells (**Figure 1B**). Particularly, gene signatures related to lipid metabolism were strongly associated with FA and cholesterol biosynthesis, as observed in AML cell lines (**Figure 1B, Table S4** and **Figure S1E**). We treated six *FLT3*-mutant PDXs *in vivo* with GILT after disease establishment (ie. 8-50 weeks after xenotransplantation; **Figure 1C**). The genomic landscape of these samples revealed frequent concurrent *NPM1* and *DNMT3A* mutations, as expected^1^ (**Figure 1D** and **Table S3**). Moreover, we detected *FLT3* D835Y and *NRAS* G12D mutations in TUH84 and TUH86, and TUH73 patient samples, respectively (**Figure 1D**). After one week of treatment, GILT significantly reduced tumor burden in bone marrow (BM) and spleen (SP) of four PDXs (**Figure 1E** and **S2A-D**). In PDX TUH86 and TUH93, we observed significant tumor reduction in the SP but not in the BM after 7 days GILT, while both compartments had decreased leukemia burden after two weeks treatment in PDX TUH93, defining early and late responders to GILT *in vivo* (**Figure 1D-E** and **S2A-D**). Notably, the presence of *FLT3*-TKD or *NRAS* mutations was not associated with delayed GILT response in these assays (**Figure 1D-E**). We performed differential gene expression analysis in human leukemic cells sorted from vehicle- and GILT-treated mice, and showed a depletion of gene signatures related to glucose, amino acids, mitochondria and lipid metabolism in the GILT-treated conditions (**Figure 1F** and **Table S5**). Among lipid-related signatures, FA and cholesterol biosynthesis pathways were the most prominently depleted by GILT treatment (**Figure 1F** and **Table S5**).

**Figure 1:**
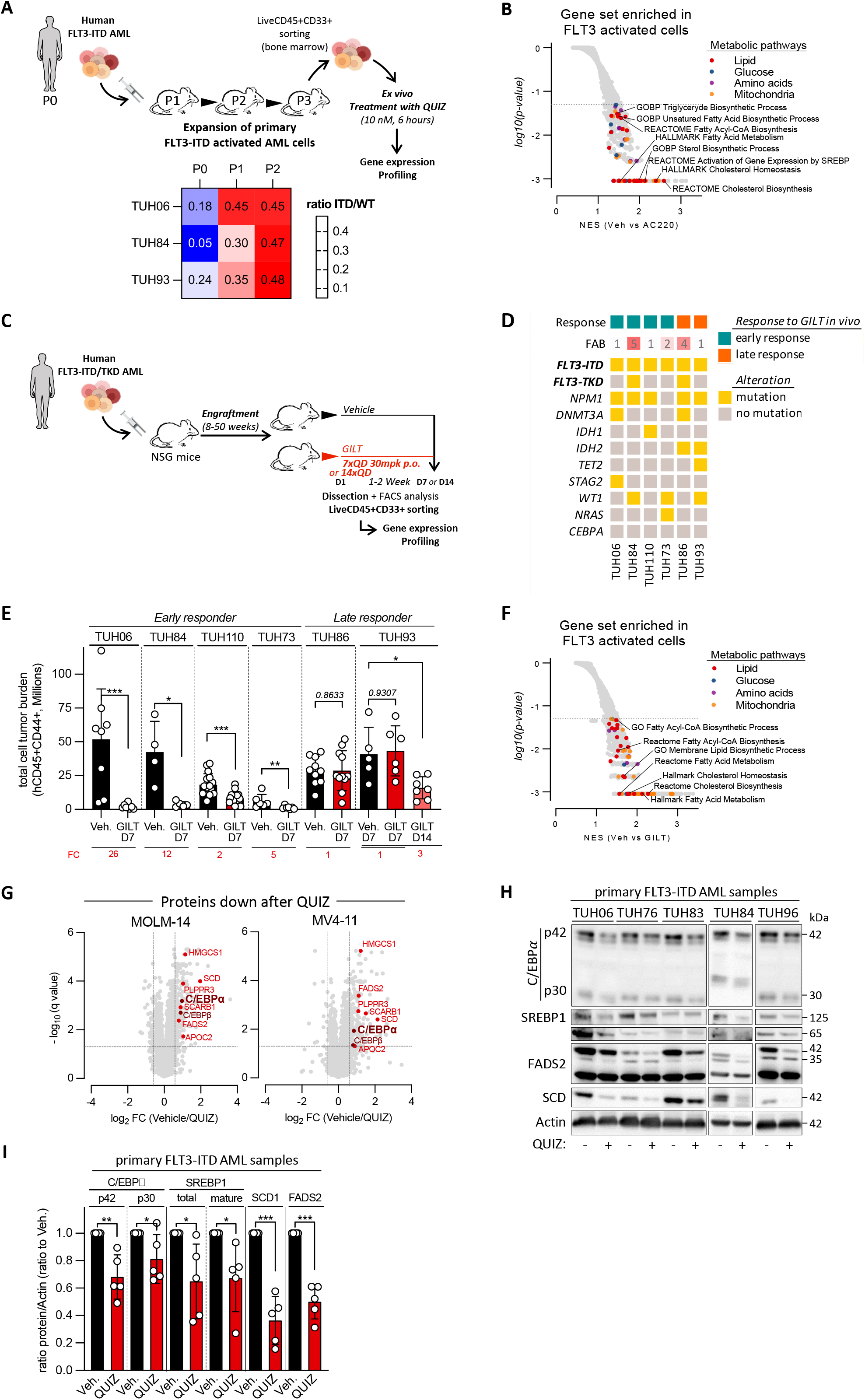
FLT3-mutant AML cells display increased lipid biosynthesis. **(A)** Experimental scheme of *in vivo* amplification of *FLT3*-ITD primary samples from patients with AML by serial transplantation in NSG mice (n=3). Human leukemic cells were sorted from mice bone marrow samples using hCD33/hCD45 staining, and incubated *ex vivo* with vehicle or 10nM quizartinib (QUIZ) during 6h before processing RNA samples for transcriptomics. *Bottom panel* represent the evolution of the ratio between FLT3-ITD mutation and its non-mutated counterpart during serial transplantation of PDX samples in NSG mice. **(B)** Differential gene expression (DGE) analysis using microarrays in patient-derived AML cells incubated with vehicle or 10nM QUIZ for 6h was analyzed using Gene Set Enrichment Analysis (GSEA) of Hallmark 2020, Biological Process (GO) and REACTOME gene sets (Normalized Enrichment Score, NES (vehicle vs QUIZ), p-value<0.05). **(C)** Schematic representation of AML PDX treatment by GILT given by oral gavage. Six different PDXs were treated with vehicle or 30mg/kg/day GILT (n=4-19 mice in each group) for 7 days. One PDX (TUH93) was also treated during 14 days. **(D)** Biological characteristics of samples from AML patients used to perform ex vivo and in vivo PDX assays. Mutations in *NPM1, IDH1, IDH2, DNMT3A, TET2, STAG2, WT1, CEBPA* and *FLT3* searched among a panel of 41 genes are reported. Presence of FLT3-ITD and FLT3-TKD are specified. French-American-British (FAB) subtypes are provided. **(E)** Number of viable CD33+/CD45+ human AML cells in bone marrow and spleen of six AML PDXs in vehicle and GILT groups. Fold change (FC) between the mean number of leukemic cells in vehicle and GILT groups are provided. **(F)** Differential gene expression (DGE) analysis using microarrays in vehicle- or GILT-treated PDXs analyzed using GSEA of Hallmark 2020, Biological Process (GO) and REACTOME gene sets (NES (vehicle vs GILT), p-value<0.05). **(G)** MOLM-14 and MV4-11 cell lines were incubated with vehicle or 3nM QUIZ for 14h. Quantitative proteomic analysis using LC-ESI-MS/MS was performed on these samples and results of fold-changes between vehicle and QUIZ on FDR q-value are provided (n=3). **(H-I)** Primary FLT3-ITD AML cells were incubated with vehicle or 10nM QUIZ for 14h. Western blots on total protein extracts were done using anti-C/EBPα, -SREBP1, FADS2, SCD and actin antibodies **(H)** and quantified **(I)**. Vertical bars indicate standard deviations. *: p<0.05; **: p<0.01; ***: p<0.001.

We next aimed to identify important effectors of lipid metabolism upon FLT3 inhibition in AML. To capture these variations, we performed a quantitative proteomic analysis in FLT3i-treated MOLM-14 and MV4-11 cells. First, we observed a downregulation of Stearoyl-CoA Desaturase (SCD) involved in mono-unsaturated FA (MUFA) biosynthesis, and of Fatty Acid Desaturase 2 (FADS2) involved in the biosynthesis of highly unsaturated FA from poly-unsaturated FA (PUFA) precursors^31^ (**Figure 1G** and **Table S6**). Importantly, we found that the protein expression of transcription factor C/EBPα and C/EBPβ was decreased (**Figure 1G** and **Table S6**), while their respective mRNA expression remained stable upon FLT3 inhibition (**Figure S3A-B**). We confirmed that FLT3i decreased the protein expression of these lipid metabolism regulators in FLT3-mutant AML PDX samples and AML cell lines (**Figure 1H-I** and **S3C-D**). In the same samples, we found that FLT3i decreased Sterol Regulatory Element-Binding Protein (SREBP1 and SREBP2) protein expression, which are transcription factors involved in FA (SREBP1) and cholesterol (SREBP2) biosynthesis^32^ (**Figure 1H-I** and **S3C-D**). However, we observed a decrease in the mature form of SREBP1 but not of SREBP2, suggesting that FLT3 inhibition predominantly redirected lipid metabolism towards FA rather than cholesterol biosynthesis inhibition (**Figure 1H-I** and **Figure S3C-D**).

Collectively, these results indicate that mutant FLT3 signaling sustains lipid metabolism and regulate key FA biosynthesis effectors including C/EBPα and SREBP1.

### Concomitant activation of FLT3 and C/EBPα is associated with lipid biosynthetic pathways

As FLT3-mutant receptors regulate C/EBPα function towards myeloid differentiation in AML^33,34^, we hypothesized that C/EBPα could represents a novel key regulator of lipid metabolism in *FLT3*-mutant leukemic cells. First, we observed an increased frequency of *FLT3*-ITD mutations in samples expressing high compared to low *CEBPA* levels in three independent cohorts of AML patients (BeatAML^35^; TCGA^36^; GSE14468^37^; **Figure 2A**). Moreover, FLT3-related and various lipid-related gene signatures were enriched in primary AML cells from these cohorts expressing high compared to low *CEBPA* (**Figure 2B**).

**Figure 2:**
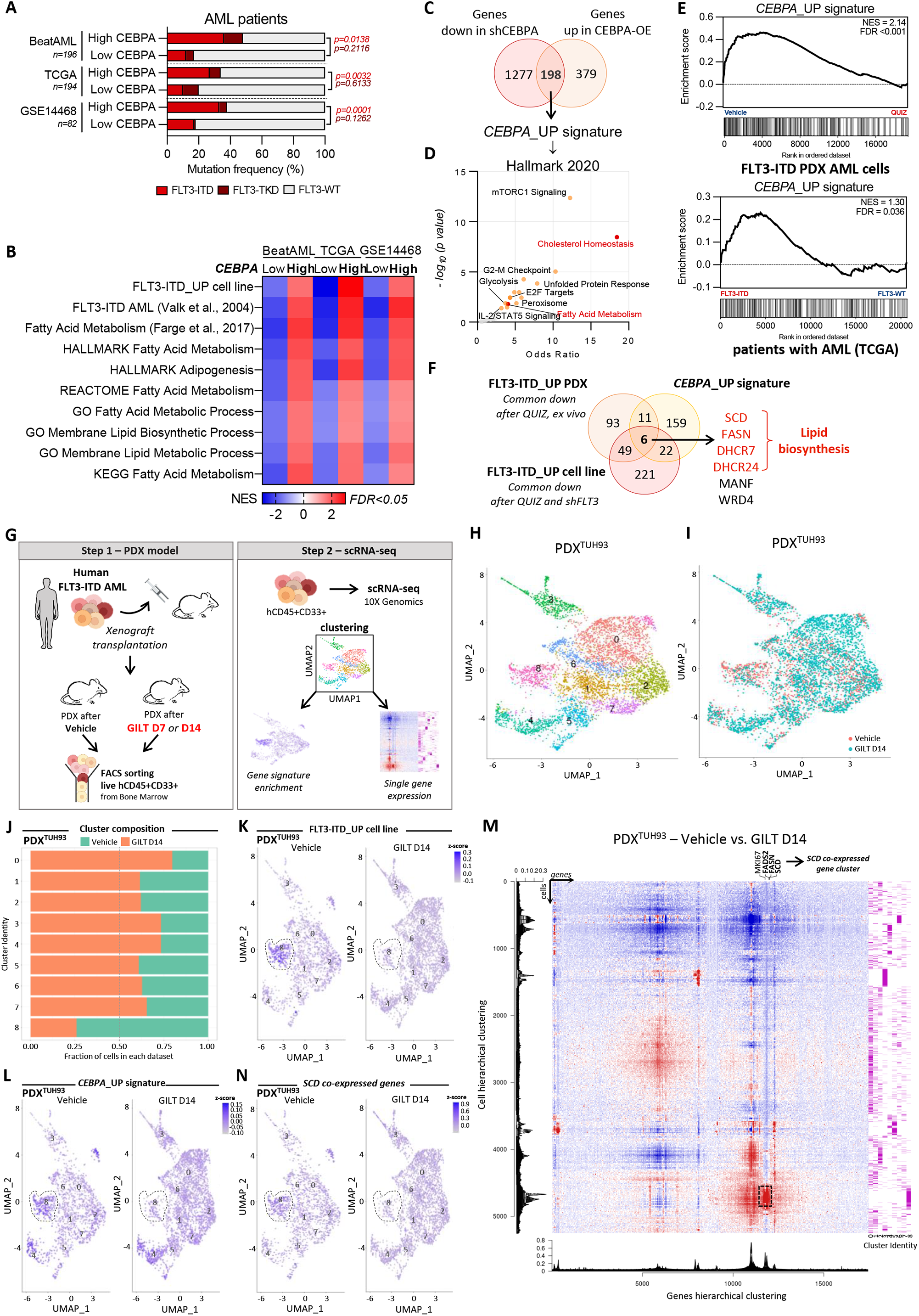
Concomitant activation of C/EBPα and FLT3 is associated with lipid biosynthetic pathways. **(A-B)** Publicly available datasets from three independent cohorts of AML patients (BeatAML, TCGA and GSE14468) were analyzed dependent on *CEBPA* low versus high expression. **(A)** *FLT3* mutation frequencies (ITD in red; TKD in dark red) in cells from patients with AML **(B)** GSEA of gene signatures related to FLT3-ITD and to lipid metabolism. **(C)** Generation of a *CEBPA*_UP signature by the intersection of genes downregulated after *CEBPA* depletion (FC<0.77, FDR, q-value<0.05) and upregulated after *CEBPA* overexpression (FC>1.3, FDR, q-value<0.05). **(D)** Enrichr online software was used to analyze the Hallmark 2020 gene signatures enriched in *CEBPA*_UP signature (p-value<0.05). **(E)** Enrichment analysis of the *CEBPA*_UP signature in DGE analysis between vehicle- and QUIZ-treated patient-derived AML samples (*upper panel*) and between FLT3-ITD versus FLT3 non-mutated (WT for wild type) samples from the TCGA cohort (*lower panel*). **(F)** Intersection between *CEBPA*_UP and FLT3-ITD-UP signatures revealing six commonly modulated genes. **(G)** Schematic representation of single-cell RNA sequencing (scRNAseq) assays done in AML PDXs treated *in vivo* with vehicle or GILT. *In vivo* AML residual cells were collected from three different PDXs (TUH93, TUH84, TUH110) before and after GILT treatment. Human CD33+/CD45+ AML cells were sorted, and single-cell transcriptome was generated using 10X Genomics technology. Single cells were clustered, and gene signature and single gene enrichment analysis were performed. **(H)** Uniform manifold approximation and projection (UMAP) plot of 5395 cells from PDX^TUH93^ using Seurat. Colors indicate clusters derived with the Louvain Algorithm in Seurat.**(I)** UMAP plot colored by treatment condition (vehicle: n=1792 cells; GILT D14: n=3465 cells). **(J)**Proportion of each condition per cluster from PDX^TUH93^ **(K-L)** Visualization on the UMAP plot of FLT3-ITD_UP cell line signature **(K)** and *CEBPA*_UP signature **(L)** enrichment in vehicle-compared to GILT-treated PDX^TUH93^. **(M)** Heatmap showing the expression of each gene per cell in vehicle-compared to GILT-treated PDX^TUH93^. Dendograms represent the hierarchical clustering of cells (*left*) and genes (*bottom*). Cluster identities are shown on the right of the Heatmap. **(N)** Visualization on the UMAP plot of *SCD* co-expressed genes enrichment in vehicle-compared to GILT-treated PDX^TUH93^.

To study C/EBPα function in *FLT3*-mutant AML cells, we performed DEG analysis in MOLM-14 cells depleted in C/EBPα using constitutive shRNA (sh*CEBPA*), or overexpressing (OE) C/EBPα (*CEBPA*-OE) compared to their respective control cells (**Figure S4A-B** and **3E-F**). We identified 198 significantly up- and down-regulated genes after C/EBPα overexpression and depletion, respectively (**Figure 2C** and **Table S7**). This *CEBPA*_UP signature was enriched in genes related to cell cycle, as well as glucose, FA and cholesterol metabolism (**Figure 2D**). Interestingly, we observed an enrichment of this *CEBPA*_UP signature in control compared to FLT3i-treated AML cell lines and primary AML cells, and in publicly accessible gene expression databases of *FLT3*-mutant compared to non-mutant AML cells (**Figure 2E** and **Figure S4C**). Finally, we identified six genes in common between *CEBPA*_UP and *FLT3*-related gene signatures, and among them, four encoded key enzymes involved in FA (*SCD* and *FASN*) or cholesterol (*DHCR7* and *DHCR24*) biosynthesis (**Figure 2F**).

Next, we investigated intra-tumor transcriptional heterogeneity and response to FLT3i *in vivo* using single cell RNA sequencing (scRNA-seq) on sorted human AML cells from vehicle- and GILT-treated mice of three FLT3-ITD PDXs (**Figure 2G**). Using Seurat and unsupervised hierarchical clustering analysis on each cell transcriptomes, we uncovered distinct clusters defined by their transcriptional state (**Figure 2H, S6A** and **S7A**). The presence and proportion of cells in these states fluctuated between vehicle- and GILT-treated mice (**Figure 2I-J, S6B-C** and **S7B-C**). First, we observed an enrichment of the FLT3-ITD_UP cell line signature in GILT-sensitive clusters from vehicle highlighting the cells harboring an activation of FLT3-ITD (cluster #8 for PDX^TUH93^; cluster #0, #2 for PDX^TUH84^; cluster #1, #3 and #4 for PDX^TUH110^; **Figure 2J-K, S5A, S6C-E** and **S7C-E**). This was associated with a concomitant enrichment of *CEBPA*_UP signature in the most sensitive cell cluster to GILT (cluster #8 for PDX^TUH93^; cluster #2 for PDX^TUH84^; cluster #3 for PDX^TUH110^; **Figure 2L, S5B, S6F-G** and **S7F-G**). Interestingly, *SCD, FASN* and *FADS2* were found enriched in cells with an activation of FLT3-ITD in vehicle-groups which were abolished upon GILT treatment (**Figure 2M, S6H** and **S7H**). Based on the single cell gene expression of *SCD*, we generated the consensus cluster of *SCD* co-expressed genes and applied it to all PDXs, independently. This consensus cluster was found enriched in cells harboring an activation of both FLT3-ITD and C/EBPα in vehicle-groups (cluster #8 for PDX^TUH93^; cluster #2 for PDX^TUH84^; cluster #3 for PDX^TUH110^; **Figure 2M-N, S5C, S6H-J** and **S7H-J**). Notably, enrichment in both FLT3-ITD_UP cell line and CEBPA_UP signatures was not restricted to cells harboring a high expression of *MKI67* gene related to cell cycle (**Figure S5D-E, S6K-L** and **S7K-L**).

Together, these transcriptomic analyses showed that activation of *CEBPA* and *FLT3* gene programs is concomitant to control lipid biosynthetic pathways in leukemic cells.

### C/EBPα regulates rate-limiting lipid biosynthetic enzymes downstream of FLT3-ITD

We next investigated the role of C/EBPα in regulating the expression of target lipid biosynthetic enzymes in the response of FLT3-mutant cells to FLT3i. First, we analyzed the transcriptomes of primary blasts from nine AML patients before and during GILT therapy (**Figure 3A; Table S3**) and observed two groups of patients in regard of *CEBPA*_UP signature score before GILT treatment (**Figure 3B**). We also found two transcriptional profiles based on the evolution of *CEBPA*_UP, FLT3-ITD_UP and SCD co-expressed gene signatures in response to this therapy. These signatures were concomitantly depleted or enriched in transcriptomes from eight GILT-treated patients compared to matched transcriptomes before GILT initiation, respectively (**Figure 3C**). This confirmed a correlation between FLT3, C/EBPα and SCD activity in patients with *FLT3*-mutated AML treated with GILT (**Figure 3C**).

**Figure 3:**
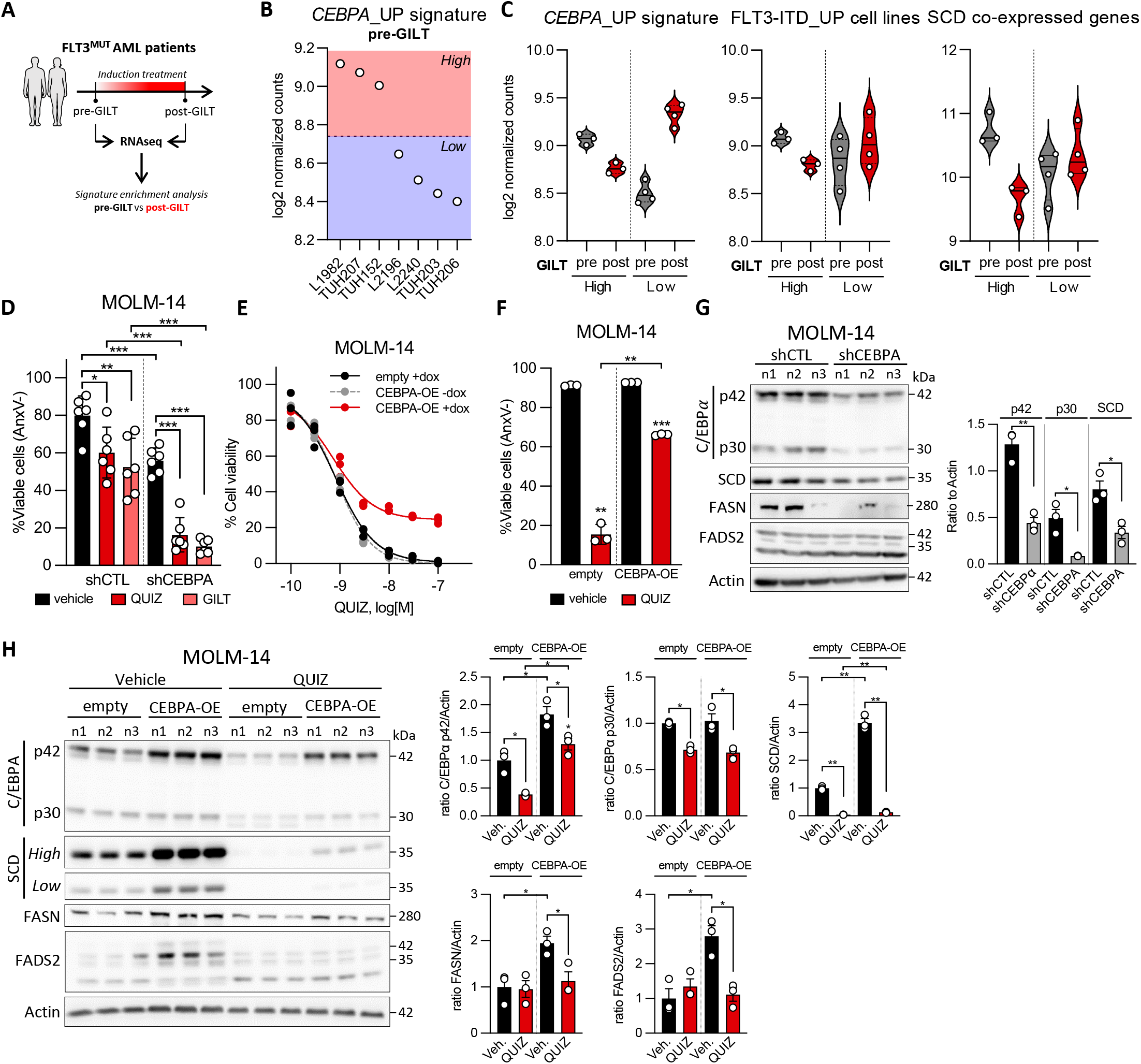
C/EBPα regulates rate-limiting lipid biosynthetic enzymes downstream of FLT3-ITD. **(A)** Bulk RNAseq performed on bone marrow or blood specimens from height patients with *FLT3*-mutant AML collected before or during GILT therapy. **(B)** Enrichment of *CEBPA*_UP signature in FLT3-mutant AML collected before GILT. Two groups of FLT3-mutant AML are highlighted: those with a High compared to those with a Low enrichment of *CEBPA*_UP signature. The dotted line correspond to the mean enrichment of the signature. **(C)** Enrichment of *CEBPA*_UP, FLT3-ITD_UP, and SCD co-expressed gene signatures in FLT3-mutant AML collected before or during GILT therapy. **(D)** MOLM-14 cells transduced with doxycycline (Dox)-inducible control (CTL) or anti-*CEBPA* shRNAs were seeded with 1µg/ml Dox during 3 days, and then incubated with 2nM QUIZ or 25nM GILT for 24h. Cell viability was assessed by annexin V staining. **(E-F)** MOLM-14 cells were transduced with a Dox-inducible vector allowing *CEBPA* overexpression (OE), or with the empty vector, and cultured without or with Dox for 48h. **(E)** CEBPA-OE (without or with Dox) and control cells (with Dox) were incubated with dose-range QUIZ. Viability was assessed by a luminescence-based cell viability assay (Alamar Blue). **(F)** Cell viability by annexin V staining in Dox-treated control or CEBPA-OE MOLM-14 cells incubated with 3nM QUIZ. **(G)** Total protein extracts from shCTL or shCEBPA MOLM-14 cells incubated with Dox for 3 days were immunoblotted with the indicated antibodies (*left panel*). Quantification are shown in *right panel*. **(H)** Total protein extracts from Dox-treated control or CEBPA-OE MOLM-14 cells incubated with vehicle or 3nM QUIZ for 14h were immunoblotted with the indicated antibodies (*left panel*). Quantification were shown in *right panel*. Vertical bars indicate standard deviations. *: p<0.05; **: p<0.01; ***: p<0.001.

In *FLT3*-ITD AML cell lines, C/EBPα depletion by RNA interference significantly reduced the sensitivity threshold to FLT3i (**Figure 3D** and **S8A-B**). Conversely, cytotoxicity of FLT3i was significantly alleviated in FLT3-ITD AML cells overexpressing C/EBPα (**Figure 3E-F**). Interestingly, gene expression of enzymes involved into lipid metabolism including SCD, FASN and FADS2 decreased upon C/EBPα depletion, while C/EBPα overexpression increased their expression (**Figure 3G-H**). Moreover, C/EBPα-induced SCD, FASN and FADS2 expression was abrogated after incubation of AML cells with FLT3i (**Figure 3H** and **S8C-K**). Interestingly, short-term incubation with FLT3i strongly depleted C/EBPα protein expression without modification of its inhibitory phosphorylation at Ser21 (ratio P^S21^-C/EBPα / total C/EBPα p42; **Figure S8A-J**). Then, we used a small compound SCD inhibitor (MF-438^38^), which demonstrated a moderate anti-leukemic activity as single agent, and a synergistic effect when combined with FLT3i against *FLT3*-mutant AML cell lines and patient-derived samples (**Figure S8L**).

Collectively, these results highlight that lipid biosynthetic enzymes (particularly SCD) are important pro-survival effectors downstream of the FLT3-C/EBPα axis in AML cells.

### C/EBPα drives fatty acid biosynthetic fluxes in *FLT3*-mutant leukemic cells

To characterize lipid metabolism in *FLT3*-mutant AML, we performed global lipidomic analyses in MOLM-14 and MV4-11 cells treated with FLT3i. We observed a global decrease in glycerophospholipid (GPs) categories, including phosphatidylethanolamines (PEs) and phosphatidylcholines (PCs), and an increased triglycerides (TGs) content after FLT3i (**Figure S9A** and **Table S8**). To evaluate if these lipid metabolic changes were driven by C/EBPα, we performed global lipidomic analyses in C/EBPα-depleted MOLM-14 and in C/EBPα-OE MOLM-14 cells treated with QUIZ or vehicle. While C/EBPα depletion recapitulated the lipidomic profile observed upon FLT3i, C/EBPα-OE showed the opposite with increased PEs, PCs and their derivates (lyso-PCs (LPCs), lyso-PEs (LPEs)) and decreased TGs content, and strikingly this lipid distribution was abolished upon FLT3i (**Figure 4A** and **S9B-H** and **Table S8**). As lipid droplets represent a reservoir for intracellular TG, we also performed electron microscopy and we observed that FLT3 inhibition led to increased number of lipid droplets in *FLT3*-mutant AML cell lines (**Figure S10A-B**). Similarly, C/EBPα depletion increased cellular neutral lipid content in MOLM-14 cells (**Figure S10C**). However, we observed a decreased expression of TG carriers including FABP5, SLC27A4 and SLC27A2, and an increase in FA transporters such as CD36 or SLC27A3 in leukemic cells after FLT3 or C/EBPα inhibition (**Figure S10D**). Accordingly, FLT3 or C/EBPα inhibition led to decreased lipid import through the plasma membrane in *FLT3*-mutant patient-derived cells and cell lines (**Figure S10E**). Finally, ceramides (CER) and sphingomyelins (SM) were increased in *FLT3*-mutant cell lines after FLT3 or C/EBPα inhibition (**Figure 4A** and **Figure S6A-B)**.

**Figure 4:**
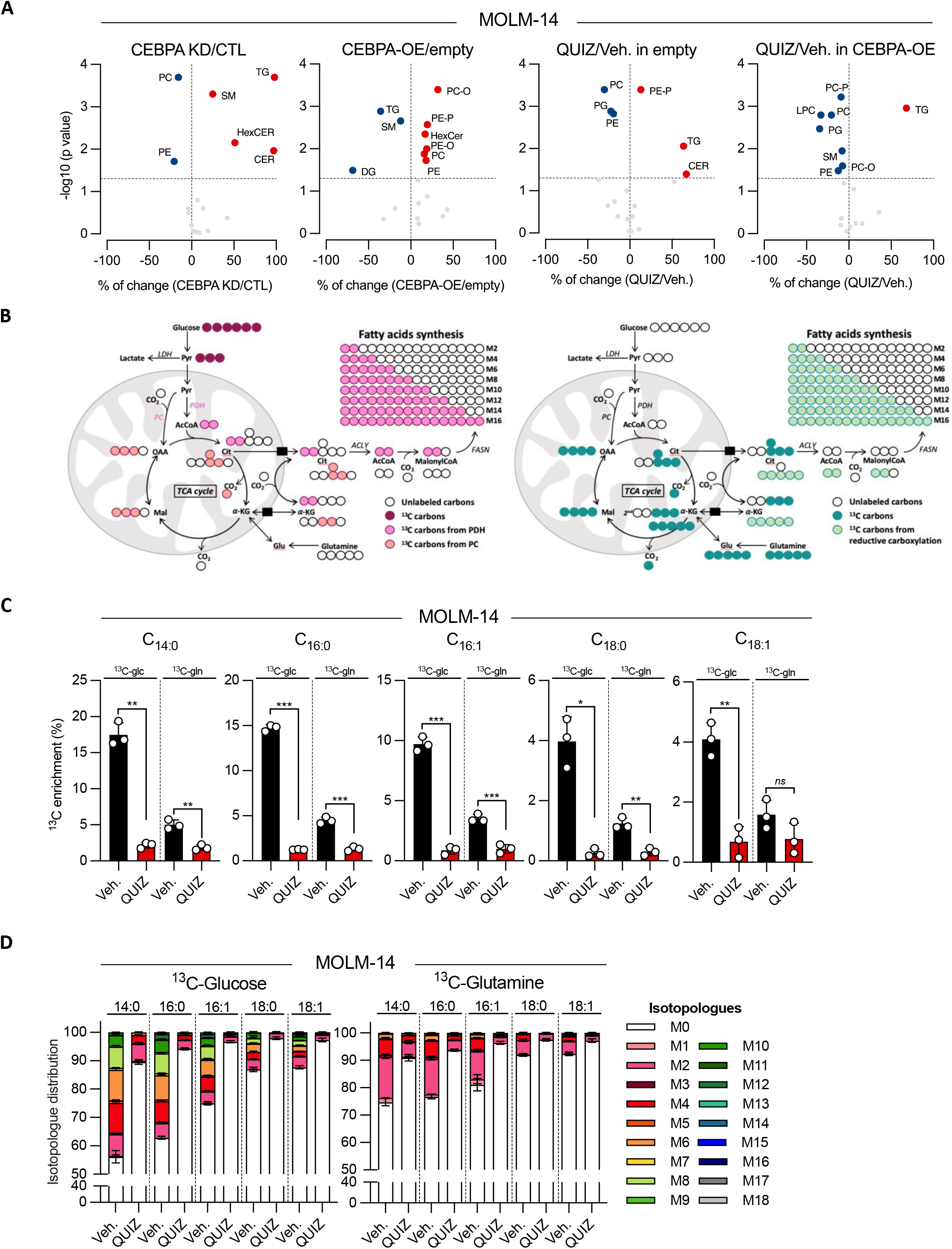
FLT3 drives fatty acid biosynthetic fluxes in leukemic cells. **(A)** Global lipidomics in MOLM-14 cells depleted from C/EBPα (siRNA 24h, dox-inducible shRNA or constitutive shRNA 72h) or overexpressing C/EBPα (dox-inducible 72h) and treated with vehicle or 3nM QUIZ for 14h. PC/PC-P/PC-O: phosphatidylcholine/-plasmogen/-alkyl ether; PE/PE-P/PE-O: phosphatidylethanolamaine/-plasmogen/-alkyl ether; PG: phosphatidylglycerol; LPC: lysophosphatidylcholine; LPE: lysophosphatidylethanolamine; SM: sphingomyelin; CER: ceramide; HexCER: hexosylceramide; TG: triacylglyceride. **(B)** Schematic representation of carbon atom (circles) transitions and tracers used to detect labeled FAs. Isotopic label from [U-^13^C]-glucose (red) or [U-^13^C]-glutamine (blue) to FAs synthesis through Pyruvate Carboxylase (PC; pink) or Pyruvate DeHydrogenase (PDH; orange), tricarboxylic acid (TCA) cycle (blue), or reductive glutamine metabolism (green). Pyr: pyruvate; AcCoA: acetyl-CoA; Cit: citrate; α-KG: α-ketoglutarate; Mal: malate; OOA: oxaloacetate; ACLY: ATP citrate lyase; FASN: Fatty acid synthase; Glu: glutamate. **(C-D)** MOLM-14 seeded with [U-^13^C]-glucose- or [U-^13^C]-glutamine-containing medium and treated with vehicle or 2nM QUIZ for 24h. **(C)** Percentage of ^13^C enrichment in C_14:0_, C_16:0_, C_16:1_, C_18:0_ and C_18:1_ total FA. **(D)** Isotopologue distribution in C_14:0_, C_16:0_, C_16:1_, C_18:0_ and C_18:1_ total FA. Vertical bars indicate standard deviations. *: p<0.05; **: p<0.01; ***: p<0.001.

In addition, we performed untargeted lipidomic analysis using liquid chromatography coupled with high-resolution mass spectrometry (LC-HRMS) of MOLM-14 and MV4-11 incubated with FLT3i and grown with stable uniformly labeled isotopologues of ^13^C_6_-glucose (^13^C_6_-Glc) or ^13^C_5_-glutamine (^13^C_5_-Gln) (**Figure 4B**). The mean enrichment of both ^13^C_6_-Glc and ^13^C_5_-Gln into newly synthetized FA was significantly decreased after FLT3i treatment (**Figure 4C** and **S11A**). Moreover, isotolopogue distribution showed a marked reduction of isotopes incorporation from 14:0 species, suggesting that fluxes of both glucose and glutamine oxidation to support MUFA biosynthesis were reduced upon FLT3 inhibition in *FLT3*-mutant leukemic cells (**Figure 4D** and **Figure S11B**).

Collectively, targeted and isotope tracing lipidomics revealed that FLT3 regulated FA biosynthetic activities in *FLT3*-mutant AML cells in a C/EBPα-dependent manner.

### FLT3 inhibition decreases mono-unsaturated fatty acids dependent on C/EBPα

Fatty acid biosynthesis mostly provides SFAs and MUFAs, while PUFAs are available mostly by food intake and subjected to both desaturation and elongation through FADS and ELOVL, respectively^31,39^ (**Figure 5A**). SFAs and PUFAs are cytotoxic and can be stored into lipid droplets^39^. Indeed, PUFAs are highly susceptible to peroxidation, leading to lipid complexes with reactive oxygen species (ROS) that trigger cell death^40^ (**Figure 5A**). We hypothesized that inhibition of the FLT3-C/EBPα axis could induce changes in lipid metabolism that modify the membrane phospholipid distribution of SFA:MUFA:PUFA in *FLT3*-mutant AML cells (**Figure 5A**). In lipidomic profiling experiments, we first observed a decreased incorporation of both SFAs and MUFAs into phospholipids, especially PE and PC after FLT3i or C/EBPα invalidation, when an increased addition of PUFAs was observed within the same conditions (**Figure 5B-C, S12A-E** and **Table S9**). On the contrary, C/EBPα-OE increased both SFAs and MUFAs into phospholipids which was abolished by FLT3i (**Figure 5B-C, S12A-E** and **Table S9**). Accordingly, the PUFA:MUFA ratio in phospholipids was increased after FLT3 or C/EBPα inhibition and decreased after C/EBPα-OE in MOLM14 cells (**Figure 5D, S13** and **Table S10**).

**Figure 5:**
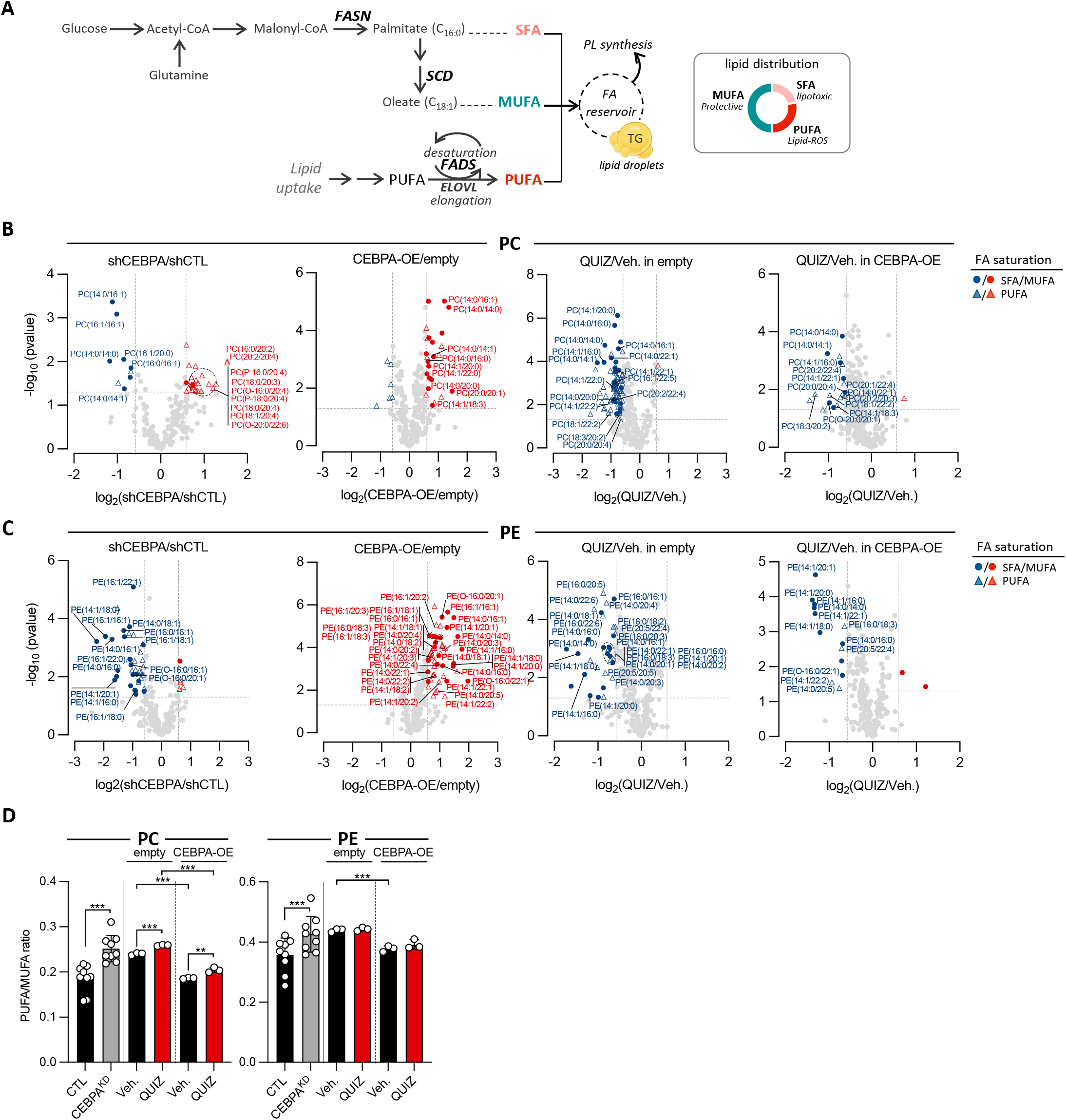
FLT3 inhibition decreases mono-unsaturated fatty acid dependent on C/EBPα. **(A)** Schematic representation of FA synthesis pathways. FASN: fatty Acid Synthase; SCD: Steroyl-CoA Desaturase; FADS: Fatty Acid Desaturase; ELOVL: Elongation of very long chain fatty acids protein; SFA: saturated fatty acid; MUFA: mono-unsaturated fatty acid; PUFA: polyunsaturated fatty acid; Lipid-ROS: lipid peroxides; PL: phospholipids; FA: fatty acid; TG: triglyceride. **(B-D)** Targeted lipidomics showing repartition of SFAs, MUFAs and PUFAs distribution into phosphoethanolamine (PE) **(B)** and phosphatidylcholine (PC) **(C)** in C/EBPα-depleted MOLM-14 compared to control, CEBPA-OE compared to empty MOLM-14, empty MOLM-14 incubated with vehicle or 3nM QUIZ for 14h, and CEBPA-OE MOLM-14 incubated with vehicle or 3nM QUIZ for 14h (from the left to the right, respectively). PUFA:MUFA ratio for PC and PE are shown in **(D)**. Vertical bars indicate standard deviations. ns: not significant; *: p<0.05; **: p<0.01; ***: p<0.001.

These results indicate that C/EBPα is essential to preserve lipid homeostasis through the generation of MUFAs and the control of PUFAs import and storage in *FLT3*-mutant AML cells.

### FLT3i-induced ferroptotic cell death is mediated by inhibition of SCD-dependent mono-unsaturated FA synthesis

Increased lipid peroxidation triggers cell death through a non-apoptotic process referred to as ferroptosis^40^. Conversely, the anti-oxidant enzyme glutathione peroxidase 4 (GPX4) specifically catalyze the reduction of lipid peroxides using reduced glutathione (GSH)^41^. To investigate the vulnerability of *FLT3*-mutant AML cells to lipid redox stress, we used RLS3, a small compound GPX4 inhibitor^41^ (**Figure 6A**). In a dose-dependent manner, RSL3 killed *FLT3*-mutant AML cells with an IC_50_ of 266 nM and 142 nM in MOLM-14 and MV4-11 cell lines, respectively, and this effect was fully rescued by the synthetic antioxidant ferrostatin 1 (Fer1) (**Figure S14A**). Moreover, the cytotoxic effect of RSL3 was abrogated by incubation of leukemic cells with oleate, a mono-unsaturated C_18:0_ FA, which also markedly alleviated RSL3-induced lipid oxidation in MOLM-14 and MV4-11 cells (**Figure 6B-D**).

**Figure 6:**
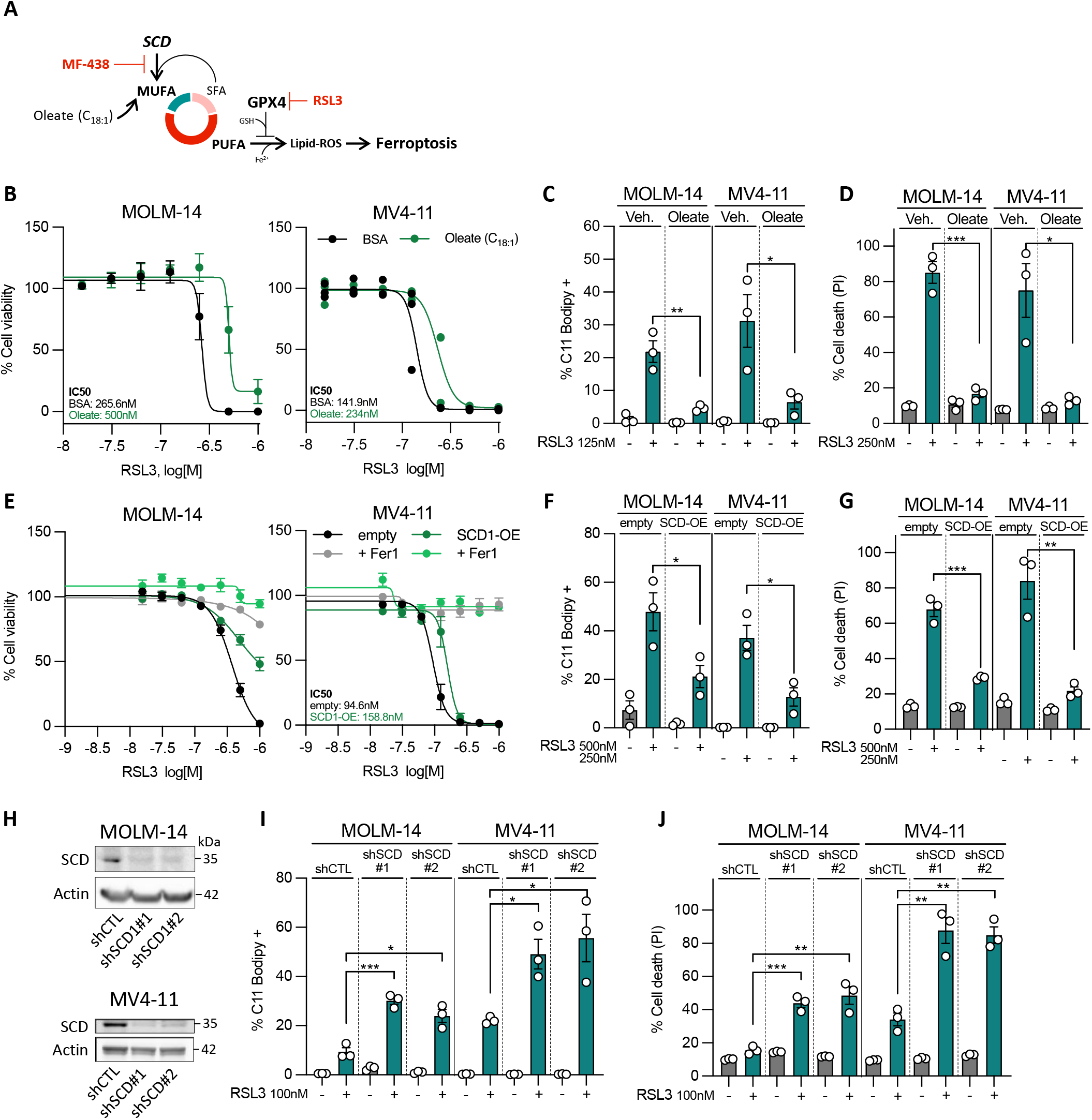
Ferroptotic cell death induced by FLT3i is mediated by inhibition of SCD-dependent mono-unsaturated FA synthesis. **(A)** Decreased amount of MUFA through SCD inhibition or increased amounts of PUFA through GPX4 inhibition lead to lipid peroxides (lipid-ROS) accumulation and cell death by ferroptosis. **(B)** Cell viability assays using Alamar Blue in MOLM-14 (*left panel*) and MV4-11 (*right panel*) supplemented with 90µM oleate (C_18:1_) or control (BSA) and exposed to dose-range RSL3 for 24h. **(C-D)** MOLM-14 cells supplemented with or without oleate were incubated with vehicle or 125nM **(C)** and 250nM RSL3 **(D)**. Lipid peroxidation was measured by C11 Bodipy staining after 12h-treatment **(C)** and cell viability measured by PI staining after 24h-treatment **(D). (E)** Cell viability assays using Alamar Blue in MOLM-14 (*left panel*) and MV4-11 (*right panel*) cells overexpressing SCD (SCD-OE) or the empty vector, and incubated with dose-range RLS3 for 14h, and with vehicle or 10µM Ferrostatin-1 (Fer-1). **(F-G)** Control or SCD-OE MOLM-14 and MV4-11 cells were incubated with vehicle or 250-500nM RLS3. Lipid peroxidation measured by C11 Bodipy staining after 12h-treatment **(G)** and cell viability assays using PI staining after 24h-treatment **(F). (H)** Total protein extracts from MOLM-14 (*top panel*) and MV4-11 (*bottom panel*) cells transduced with constitutive shRNAs against *SCD* (shSCD#1 and shSCD#2) or shCTL for 48h and then immunoblotted with indicated antibodies. **(I-J)** MOLM-14 and MV4-11 cells transduced with CTL or anti-SCD shRNAs were incubated with 100nM RSL3. Lipid peroxidation measured by C11 Bodipy staining after 12h-treatment **(I)** and cell viability assessed by PI staining after 24h-treatment **(J)**. Vertical bars indicate standard deviations. *: p<0.05; **: p<0.01; ***: p<0.001.

To increase endogenous MUFA production, we overexpressed SCD, the rate-limiting enzyme of the last step of MUFA biosynthesis^42^, in MOLM-14 and MV4-11 cell lines (**Figure S14B**). We observed that SCD-OE markedly protected leukemic cells from lipid peroxidation and cell death induced by cytotoxic concentrations of RSL3 (**Figure 6E-G**). Conversely, SCD depletion by shRNA primed *FLT3*-mutant AML cells to lipid peroxidation and cell death induced by non-cytotoxic concentrations of RSL3 (**Figure 6H-J**). Similar results were observed with the small compound SCD inhibitor MF-438 (**Figure S14C-E**).

These results showed that SCD inhibition promotes lipid peroxidation and cytotoxicity, representing a new vulnerability of *FLT3*-ITD leukemic cells.

### C/EBPα-mediated lipid oxidative stress primes *FLT3*-mutant AML cells to ferroptotic cell death upon FLT3 inhibition

In *FLT3*-mutant cell lines, cell death observed after C/EBPα depletion was increased by RSL3, while C/EBPα overexpression protected leukemic cells from GPX4 inhibition (**Figure 7A**). Similarly, combination of RLS3 with FLT3i enhanced lipid peroxidation and cell death in *FLT3*-ITD cell lines (**Figure 7B-C** and **Figure S15A**). This was confirmed in patient-derived AML cells *ex vivo* in which RLS3 was synergistic with FLT3i to induced cell death (**Figure 7D**). We observed similar effects in *FLT3*-depleted AML cell lines incubated with RSL3 (**Figure S15B-C**). As FLT3i and RSL3 combination induced lipid-ROS and cell death both in MOLM-14 and MV4-11 cell lines harboring heterozygous and homozygous *FLT3*-ITD mutation, respectively^43^, we hypothesized that these effects were mostly dependent on *FLT3*-mutant alleles. We used Ba/F3 murine hematopoietic cell line constitutively expressing *FLT3*-ITD (Ba/F3-ITD, growth factor independent), which display exclusive addiction to mutant FLT3 signaling^44^, or non-mutated *FLT3* allele (Ba/F3-WT, dependent on FLT3 ligand) (**Figure S15D**). Ba/F3-ITD cells were highly susceptible to FLT3i, in contrast to Ba/F3-WT cells, as reported^45^ (**Figure S15E**). Moreover, co-incubation with FLT3i and RSL3 had no impact on Ba/F3-WT cells, while near-completely killing Ba/F3-ITD cells *in vitro* (**Figure S15E**). Interestingly, RSL3 enhanced the vulnerability of MOLM-14 and MV4-11 cells to three other FLT3 inhibitors (crenolanib, midostaurin and sorafenib, **Figure S15F**). Finally, the vulnerability to FLT3i was enhanced by two other GPX4 inhibitors, FIN56 and ML210 in MOLM-14 cells (**Figure S15G-H**). These results showed that FLT3 inhibition primed *FLT3*-mutant leukemic cells to lipid oxidation and ferroptotic cell death.

**Figure 7:**
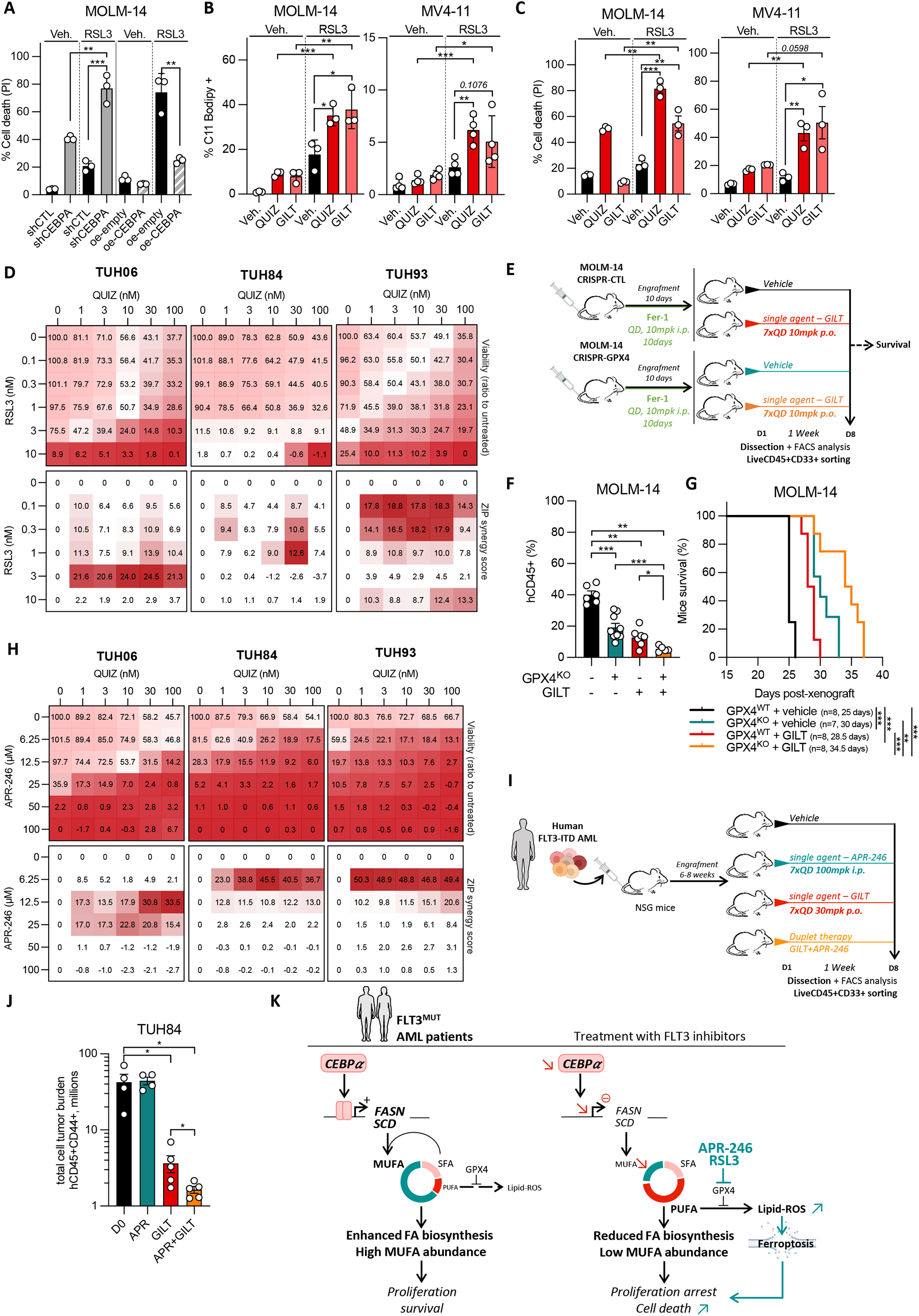
Lipid oxidative stress mediated by C/EBPα inhibition primes *FLT3*-mutant AML cells to ferroptotic cell death during FLT3 inhibition. **(A)** MOLM-14 cells in which C/EBPα was depleted or overexpressed were incubated with vehicle or 100nM or 250nM RSL3 respectively for 24h. **(B)** Lipid peroxidation measured using C11-Bodipy staining in MOLM-14 and MV4-11 cells treated with vehicle or FLT3i (3 and 10 nM QUIZ or 30 and 100nM GILT, respectively), and vehicle or 100nM RSL3 for 12h. **(C)** Cell viability assessed by PI staining in MOLM-14 and MV4-11 cells treated with vehicle or FLT3i (3nM QUIZ or 30nM GILT), and vehicle or 100nM RSL3 for 24h. **(D)** Patient-derived AML samples (n=3) were incubated *ex vivo* with crossed dose-range QUIZ and RSL3 for 48h. Cell viability was measured using a luminescence-based assay (Alamar Blue) and results are expressed as a ratio to the untreated condition for each wheel in a viability matrix (*top panel*) and using synergy maps using zero interaction potency (ZIP) synergy scores and a threshold ZIP>10 considered as significant (*bottom panel*). **(E-G)** MOLM-14 cells were transduced with CTL or GPX4-targeting CRISPR-Cas9 vectors, and cultured with 10µM Fer-1 to avoid cell death induced by GPX4 ablation. These cells were xenografted to immunodeficient NSG mice and treated in vivo with 10mg/kg/day Fer-1 by intraperitoneal (IP) injection during 10 days. Then CTL and GPX4 KO groups were treated with vehicle or 10mg/kg/day GILT by oral gavage for 7 days before evaluation of tumor burden or follow-up for survival. Each group was made of 5-9 mice. **(E)** Schematic overview of the experimental design. **(F)** Quantification of human CD45+ leukemic cell burden at day 7. **(G)** Survival curves. **(H)** Patient-derived AML samples (n=3) were incubated ex vivo with crossed dose-range QUIZ and APR-246 for 48h. Cell viability was measured using a luminescence-based assay (Alamar Blue) and results are expressed as a ratio to the untreated condition for each wheel in a viability matrix (*top panel*) and using synergy maps using zero interaction potency (ZIP) synergy scores and a threshold ZIP>10 considered as significant (*bottom panel*). **(I-J)** Patient-derived xenograft (PDX) assays using TUH84 sample. After engraftment, mice were treated with vehicles, 100mg/kg/day IP APR-246, 30mg/kg/day oral GILT or combination of these two agents for 7 days. **(I)** Schematic representation of the experiment. **(J)** Evaluation of human CD45+ leukemic cell burden at day 7. **(K)** Schematic diagram of lipid metabolism reprograming induced by C/EBPα in FLT3-ITD AML cells through FASN/SCD/FADS2 at the steady state and upon inhibition of C/EBPα or treatment with FLT3i combined with ferroptosis inducers. Vertical bars indicate standard deviations. *: p<0.05; **: p<0.01; ***: p<0.001.

We next investigated the cooperative lethality between redox stress induction and FLT3 inhibition *in vivo*. We depleted *GPX4* by CRISPR/Cas9 in MOLM-14 cells (**Figure S16A**). As *GPX4* knockout (KO) is lethal in our model, AML cells were grown with Fer1 that efficiently prevented lipid oxidation and cell death in GPX4-depleted cells, while Fer1 removal promptly resulted in the opposed effect (**Figure S16B**). Using a different approach by RNA interference, we similarly observed that GPX4 depletion induced leukemic cell death, which was further enhanced by FLT3i treatment (**Figure S16C**). We performed cell line derived xenograft (CLDX) assays using GPX4^KO^ or GPX4^WT^ (transduced with control CRISPR) MOLM-14 cells in immunodeficient NSG mice. Starting ten days from transplant at a stage of established disease, mice were treated by GILT or vehicle for one week before the assessment of leukemia burden, or followed-up for survival without further treatment (**Figure 7E**). Human engraftment and total cell tumor burden were significantly reduced in GPX4^WT^ mice treated with GILT, and in vehicle-treated GPX4^KO^ mice, but residual disease was nearly eradicated in GPX4^KO^ mice treated with GILT (**Figure 7F** and **Figure S16D**). This marked tumor reduction was associated with an increased survival of GPX4^KO^ mice treated with GILT (**Figure 7G**). Together, these results show that this induction of lipid redox stress induction primed *FLT3*-ITD AML cells to FLT3i *in vitro* and *in vivo*.

Finally, we and others previously showed that APR-246, a promising therapeutic agent in AML^46^, induces AML cell killing^47,48^ by decreasing glutathione salvage and increasing lipid peroxidation^49–51^. We therefore investigated the combination between FLT3i and two ferroptosis inductors RLS3 and APR-246 on primary patient AML cells *ex vivo*, and observed that both RLS3 and APR-246 were synergistic with FLT3i (**Figure 7H**). Notably, APR-246 was also synergistic with other FLT3i including quizartinib, midostaurin and crenolanib *in vitro*, and enhanced lipid oxidation and cytotoxicity induced by shRNA-mediated FLT3 depletion in *FLT3*-ITD cell lines (**Figure S17A-C**). Since APR-246 demonstrated activity against high-grade myeloid neoplasms in clinical trials (NCT03072043)^46^, we treated established AML PDX (TUH84) with GILT and/or APR-246 during one week, and observed that combination of FLT3i and APR-246 resulted in a minimal residual disease compared to treatment with single agents (**Figure 7I-J**).

Collectively, these preclinical experiments demonstrate that APR-246 increases lipid oxidation and potentiates FLT3i to kill FLT3-mutant leukemic cells.

## DISCUSSION

*FLT3* is the most frequently mutated gene in AML, leading to an overactivation of signaling pathways downstream this tyrosine kinase receptor. This results in a high proliferation rate of leukemic cells and frequent relapse after intensive chemotherapy. The practice-changing approval of FLT3i at different stage (first line, relapse and post-allogenic stem cell transplant maintenance) of AML therapy provide a significant survival gain for patients with AML^10,11,52^. However, resistance to FLT3i is common and caused by multiple mechanisms including selection of FLT3i-resistant clones harboring *FLT3*-TKD or RAS-activating mutations, or signaling or metabolic adaptations to FLT3i^19,53,54^. The discovery of novel vulnerabilities is therefore an important goal in *FLT3*-mutant AML to uncover alternative therapeutic strategies targeting post-FLT3i residual disease and preventing the emergence of resistance.

In this study, we demonstrated that the biosynthesis of lipids (including FA, glycerophospholipids, sphingolipids and triglycerides) is a critical vulnerability and dependency of *FLT3*-mutant AML mediated by the transcription factor C/EBPα. Amongst metabolic adaptations^5,9^, AML cells with *FLT3*-ITD mutations display activated FA oxidation and increased lysophospholipid amount^55^. Moreover, sphingolipid ceramide are involved in cell cycle arrest and apoptotic cell death in cancer models^56^ and FLT3-ITD represses their biosynthesis in leukemia^57^, suggesting that ceramide accumulation observed in our lipidomic assays could play a role in leukemic cell death mediated by FLT3i. Furthermore, AML cell death induced by ceramide could be regulated by GPX4 activity, as previously described with the isoenzyme GPX1^58^. Interestingly, high-fat diet promotes FLT3-dependent leukemia in MLL-AF9 AML mouse model, highlighting a crucial role of lipid homeostasis with this oncogene^59^. Interestingly, lipid dependency is observed in leukemia with other drivers such as oncogenic isocitrate dehydrogenase isoforms, which can redirect lipid metabolism towards FA oxidation to maintain mitochondrial oxidative phosphorylation by activating C/EBPα-driven transcriptional program^60^. Using a multi-omics approach *in vitro* and *in vivo*, we showed that C/EBPα is a novel essential regulator of FA biosynthesis through the activation of SCD, a rate-limiting enzyme for MUFA biosynthesis^42^, downstream *FLT3*-mutant receptors. This lipid dependency of *FLT3*-mutant AML was further observed at the single cell level *in vivo* in PDX assays, highlighting the link between SCD, lipid peroxidation, ferroptosis priming and post-FLT3i residual disease *in vivo*. In same vein, acyl-CoA dehydrogenase involved into FA metabolism protects glioblastoma cells from toxic lipid oxidation, which favors their adaptive resistance to drug-induced stress^61,62^. Together, C/EBPα may account for the high reliance of leukemic cells on sustained FA catabolic or anabolic pathways.

Isotope tracing lipidomic experiments uncovered that FLT3 inhibition strongly reduced glucose and glutamine utilization to fuel FA biosynthesis, and that C/EBPα inhibition resulted in the same shift in lipid composition with decreased and increased amounts of phospholipids and neutral lipids, respectively. Based on our previous study^63^, this accumulation of neutral lipids might be due to autophagy/lipophagy inhibition. Among lipid species, FLT3 and C/EBPα inhibition particularly reduced the amounts of MUFA through SCD inhibition alongside with increased PUFA import and storage as compensatory adaptation to FADS2 inhibition. Because imported PUFA are more susceptible to peroxidation, lipogenesis inhibitors increase the levels of peroxidation end products, render cancer cells more susceptible to oxidative stress-induced cell death, and act as chemotherapeutic sensitizers^64^. Interestingly, periodic fasting-mimicking and calorie-restriction diets inhibit SCD, decrease MUFA, alter MUFA:SFA ratio and affect tumor growth in mice^65,66^. Our study also showed that an increase in both PUFA:MUFA ratio and propensity to lipid oxidation represents a window of opportunity to trigger ferroptotic cell death^62^. Accordingly, FLT3 and C/EBPα inhibition synergized with GPX4 inhibitors to promote lipid-ROS formation and cell death, which also occurred *in vivo* in GPX4-deficient AML models.

Ferroptosis recently emerged as an iron-dependent, non-apoptotic form of regulated cell death particularly relevant to cancer therapy with implications in response to chemotherapy, radiotherapy and immunotherapy^67–69^. However, no selective GPX4 inhibitor is currently available for patient care. We recently showed that the small compound APR-246 strongly induces ferroptosis through GSH depletion^50^. Interestingly, APR-246 is now known as eprenetapopt and was sucessfully administrated to patients with myeloid neoplasms including AML in phase II clinical trials in association with azacitidine^46,70^. We therefore used APR-246 to trigger lipid oxidation in AML cells, and observed a synergy between APR-246 and diverse FLT3 inhibitors including Gilteritinib, Quizartinib, Midostaurin and Crenolanib, compared to single agents *in vitro* and *in vivo* in PDX models. This further uncovered lipid oxidation as a new target in *FLT3*-mutant AML.

In summary, we describe a novel and protective role of C/EBPα from ferroptotic cell death in cancer through the transcriptional control of SCD-driven MUFA biosynthesis. Similar role of SCD was observed in cancer and ferroptosis^71^. Our study also demonstrates that FLT3 inhibition causes metabolic adaptation in response to C/EBPα-SCD downregulation to induce lipid oxidation, which is therapeutically actionable by ferroptosis inducers in AML cells (**Figure 7K**).

## MATERIALS AND METHODS

### Samples from patients with AML

De-identified primary AML patient specimens are from Toulouse University Hospital (TUH) (Toulouse, France). Frozen samples were obtained from patients diagnosed with AML at TUH after signed written informed consent for research use in accordance with the Declaration of Helsinki, and stored at the HIMIP collection (BB-0033-00060). According to the French law, HIMIP biobank collection has been declared to the Ministry of Higher Education and Research (DC 2008-307, collection 1) and obtained a transfer agreement for research applications (AC 2008-129) after approbation by our institutional review board and ethics committee (Comité de Protection des Personnes Sud-Ouest et Outremer II). Clinical and biological annotations of the samples have been declared to the CNIL (Comité National Informatique et Libertés, i.e. Data processing and Liberties National Committee). Patient characteristics are summarized in Table S1. The genomic landscape of these leukemia samples was obtained using AmpliSeq for Illumina Myeloid Panel (Illumina, San Diego, USA). *FLT3*-ITD mutations were also investigated by quantitative and fluorescent PCR (Eurofins Genomics, Luxembourg), allowing the calculation of a ratio between ITD and wild type *FLT3* alleles.

### AML cell lines

MOLM-14 (DSMZ Cat# ACC-777, RRID:CVCL_7916) and MV4-11 (DSMZ Cat# ACC-102, RRID:CVCL_0064) human AML cell lines were identified by PCR-single-locus-technology (Promega, PowerPlex21 PCR Kit, Eurofins Genomics. MOLM-14 harbors a heterozygous FLT3-ITD mutation while MV4-11 carries a homozygous FLT3-ITD mutation. AML cells were grown in RPMI L-Glutamine (4 mM) supplemented or minimum essential medium (MEM)-α Glutamax supplemented with 10% foetal bovine serum (FBS, Invitrogen), 100 IU/mL penicillin and 100 µg/mL streptomycin (Life Technology) at 37°C with 5% CO2. BaF/3 (DSMZ Cat# ACC-300, RRID:CVCL_0161) murine cell lines were grown in RPMI-1640 supplemented with 10% FBS and 10ng/ml murine IL-3 (Peprotech, Cat# 213-13). HEK 293T/17 (ATCC Cat# CRL-11268, RRID:CVCL_1926) cells were grown in Dulbecco’s modified Eagle medium (DMEM) Glutamax (Life Technology) supplemented with 10% FBS, 100 IU/mL penicillin and 100 µg/mL streptomycin (Life Technology). Doxycycline (Sigma-Aldrich, Cat# D9891) was used at 1µg/mL. The cultured cells were diluted every 2 to 3 days and maintained in an exponential growth phase. These cell lines have been routinely tested for mycoplasma contamination in the laboratory.

### AML mouse xenograft model

Animals were used in accordance with a protocol reviewed and approved by the Institutional Animal Care and Use Committee of Region Midi-Pyrenees (France). NOD/LtSz-SCID/IL-2Rαchain null (NSG) mice were produced at the Genotoul Anexplo platform in Toulouse, France, using breeders obtained from Charles River Laboratories. Mice were housed in sterile conditions using high-efficiency particulate arrestance filtered microisolators and fed with irradiated food and sterile water. Mice (6–9 weeks old) were sublethally treated with busulfan (20 mg/kg for MOLM-14 and 30 mg/kg for PDXs) 24 h before injection of leukemic cells. Leukemia samples were thawed at room temperature, washed twice in PBS1X, and suspended in Hank’s Balanced Salt Solution (HBSS) at a final concentration of 0.2-10×10^6^ cells for PDXs and 1×10^6^ for cell line–derived xenograft (CLDX) per 200 µl of HBSS per mouse for tail vein injection. Daily monitoring of mice for symptoms of disease (ruffled coat, hunched back, weakness, and reduced mobility) determined the time of killing for injected animals with signs of distress. If no signs of distress were seen, mice were initially analyzed for engraftment 8 wk after injection except where otherwise noted. Mice were randomly assigned to the different treatment arms based on the blast counts in blood determined 3–8 days before the beginning of the treatment. The randomization was based on the number of mice with validated engraftment (higher than 20 blasts/µl of blood), sex, and body weight. The duration of treatment was determined before starting each experiment unless the animals died before the end of treatment or were euthanized for ethical reasons. During the treatments, total body weight was monitored every 2 days.

### *In vivo* mice treatment of PDXs

8–18 wk (PDXs) after AML cell transplantation and when mice were engrafted (tested by flow cytometry on peripheral blood or bone marrow aspirates), NSG mice were treated as described below: for GILT treatment, GILT was solubilized in water containing 0.5% methylcellulose 400cp (Sigma-Aldrich, Cat# M0262) before administration to mice. NSG mice were treated daily for 7 days or 14 days by gavage with 30 mpk GILT at 5ml/kg. For control, NSG mice were treated by daily gavage of vehicle. For APR-246 treatment, APR-246 was solubilized in PBS1X before administration to mice. NSG mice were treated each day during 7 days by intraperitoneal injection of 100 mpk APR-246 at 5ml/kg. For control, NSG mice were treated by daily intraperitoneal injection of vehicle. Mice were monitored for toxicity and provided nutritional supplements as needed.

### *In vivo* mice treatment of CLDXs

From the day before the injection of AML cell line, all mice were treated each day for 10 days by intraperitoneal injection of 2 mpk Fer-1 at 10ml/kg. Fer-1 was prepared in PBS1X 1%DMSO. After disease establishment (Day 10 post-xenotransplantation), NSG mice were treated daily for 7 days by gavage with 10 mpk GILT or vehicle at 5ml/kg. GILT was prepared in water containing 0.5% methylcellulose 400cp (Sigma-Aldrich, Cat# M0262) before administration to mice. Mice were monitored for toxicity and provided nutritional supplements as needed.

### Assessment of leukemic engraftment

NSG mice were humanely killed in accordance with European ethics protocols. Bone marrow (mixed from tibias, femurs and hips) and spleen were dissected and flushed in HBSS with 1% FBS. Mononuclear cells from bone marrow and spleen were labeled with mCD45.1-PerCP-Cy5.5 (Cat# 560580, RRID: AB_1727489), hCD45-APC (Cat# 555485, RRID:AB_398600) and hCD44-PECy7 (Cat# 560533, RRID:AB_1727483) or hCD33-PE (Cat# 555450, RRID:AB_395843; all antibodies from BD Biosciences), and Annexin V-V500 (BD Biosciences Cat# 561501, RRID:AB_10694254) to determine the fraction of viable human blasts (AnxV-hCD45+mCD45.1−hCD44+ or AnnxVhCD45+mCD45.1−hCD33+ cells) using flow cytometry. All antibodies used for cytometry were used at concentrations between 1/50 and 1/200 depending on specificity and cell density.

Analyses were performed on a CytoFLEX flow cytometer with CytExpert software (Beckman Coulter, RRID:SCR_017217) and FlowJo 10.2 (Tree Star, RRID:SCR_008520). The number of AML cells/µl peripheral blood and number of AML cells in total cell tumor burden (in bone marrow and spleen) were determined by using CountBright beads (Invitrogen) using described manufacturer protocol.

### Flow cytometry cell sorting

Human AML cells from AML-xenografted mice were stained with Fixable Viability Stain 510 (BD Biosciences Cat# 564406, RRID:AB_2869572) and the following fluorescent conjugated antibodies: hCD45-APCH7 (Cat# 641399, RRID:AB_1645735) and hCD33-PE (Cat# 555450, RRID:AB_395843; all antibodies from BD Biosciences) by BD FACSAria Fusion (Becton Dickinson) and FACSMelody (BD Biosciences).

### Culture of PDX-derived AML cells

Cells were thawed in a 37°C water bath and washed once in Iscove’s modified Dulbecco’s medium (IMDM) supplemented with 10% FBS and 10 µg/mL DNAse I. Next, cells were placed in IMDM Glutamax supplemented with 20% of bovine serum albumin/insulin/transferrin (BIT) medium. BIT composition was: bovine serum albumin (BSA, Sigma-Aldrich, Cat# A9418) 4 g/L, Insulin 5 µg/mL (Sigma-Aldrich, Cat# I2643) and Transferrin 60 µg/mL (Sigma-Aldrich, Cat# T8158), 50 ng/mL FLT3 ligand (Cat# 300-19), 10 ng/mL IL-6 (Cat# 200-06), 50 ng/mL SCF (Cat# 300-07), 25 ng/mL TPO (Cat# 300-18), 10 ng/mL IL-3 (Cat# 200-03) and 10 ng/mL G-CSF (Cat# 300-23; all cytokines from Peprotech), 100 IU/mL penicillin, and 100 µg/mL of streptomycin.

### Reagents

RSL3 (Cat# HY-100218A), APR-246 (Cat# HY-19980), ML-210 (Cat# HY-100003), FIN-56 (Cat# HY-103087), MF-438 (Cat# HY-15822), Ferrostatin-1 (Fer-1, Cat# HY-100579), Sorafenib (Cat# HY-10201), Gilteritinib (GILT, Cat# HY-12432), Crenolanib (Cat# HY-13223), Midostaurin (Cat# HY-10230), Quizartinib (QUIZ, Cat# HY-13001, all from MedChemExpress) were dissolved in DMSO except APR-246 which was dissolved in water. Oleate (Oleic Acid-Albumin from BSA, Cat# O3008) and linoleate (Linoleic Acid-Albumine from BSA,Cat# L9530) were purchased from Sigma-Aldrich.

### Constructs

We obtained human *CEBPA, SCD, FLT3-*WT (NM_004119.3) and *FLT3*-ITD ORFs through GeneArt gene synthesis technology (Thermo Fisher Scientific). *CEBPA* ORF was cloned in pINDUCER21 (ORF-EG) (gift from Stephen Elledge & Thomas Westbrook^72^; Addgene plasmid #46948; http://n2t.net/addgene: 46948 ; RRID:Addgene_46948). *SCD* ORF was cloned in pLenti PGK Puro DEST 5W(w529-2) (gift from Eric Campeau & Paul Kaufman^73^; Addgene plasmid #19068; http://n2t.net/addgene:19068 ; RRID:Addgene_19068). *FLT3-*WT and *FLT3*-ITD ORFs were cloned in pLEX_307 (gift from David Root (Addgene plasmid #41392; http://n2t.net/addgene:41392 ; RRID:Addgene_41392))

We cloned shRNAs against *SCD* and *GPX4* as well as control shRNA in the Tet-pLKO-puro vector (Gift from Dmitri Wiederschain, Addgene plasmid #21915; http://n2t.net/addgene:21915; RRID: Addgene_21915) allowing the conditional expression of hairpins by doxycycline. Sequences of shRNAs are (5’>3’): shCTL, CAACAAGATGAAGAGCACCAA; shSCD#1, CGTCCTTATGACAAGAACATT; shSCD#2, GCACATCAACTTCACCACATT, shGPX4#1, GTGAGGCAAGACCGAAGTAAA, shGPX4#2 CTACAACGTCAAATTCGATAT.

For FLT3 doxycycline-inducible invalidation, SMARTvector inducible human hEF1a-TurboGFP shRNA vectors purchased from Horizon Discovery were used (shFLT3: Clone Id: V3IHSHEG_10441930, CTGGAGAATACCTACTTTT; Non-targeting Control hEF1a-TurboRFP, shCTL: Cat #VSC6573)

For *CEBPA* constitutive invalidation, pLKO.1-puro constitutive vectors containing the following shRNA sequences were used (5′ > 3′): shRNA control, purchased from Sigma-Aldrich (SHC002 MISSION pLKO.1-puro non-mammalian shRNA control; CCGGCAACAAGATGAAGAGCACCAACTC); shRNA against *CEBPA*, purchased from Sigma-Aldrich (SHCLNG-NM_004364, TRCN0000007306 and TRCN0000356198). For *CEBPA* doxycycline-inducible invalidation, pTRIPZ doxycycline-inducible vectors purchased from Horizon Discovery were used (shCEBPA: RHS4696-200706732, clone ID: V2THS_14752, GGAAAGAATCAAGGAGG; shCTL: empty vector shRNA control, RHS4750).

### Lentiviral production

Briefly, we transfected HEK 293T/17 cells with different plasmids together with the packaging plasmids pMD2.G (Gift from Didier Trono (Addgene plasmid #12259; http://n2t.net/addgene:12259 ; RRID:Addgene_12259) and psPAX2 (Gift from Didier Trono (Addgene plasmid # 12260 ; http://n2t.net/addgene:12260; RRID:Addgene_12260) encoding lentiviral proteins using Lipofectamine 2000 Transfection Reagent (Thermo-Fischer Scientific). Twenty-four hours after cell transfection, medium was removed and Opti-MEM culture medium (Life Technology) was added. In order to transduce AML cell lines, HEK 293T/17 culture supernatants containing lentiviral particles were harvested 72h after transduction, filtered, and used to lentiviral infection of AML cells or stored at −80 °C.

### Lentiviral infection

On the day of transduction, 2×10^6^ MOLM-14 or MV4-11 cells were resuspended in a medium containing 8μg/mL polybrene (Sigma-Aldrich, Cat# 107689) and 2 mL of thawed lentivirus-containing HEK 293T/17 supernatant. Three days after infection, transduced cells were selected using 1μg/mL puromycin (LabForce) or sorted on an Astrios cell sorter (Beckman Coulter) dependent on the vector.

### siRNA transfection

The transfection of siRNA into MOLM-14 and MV4-11 cells was performed using the Neon transfection system (Life Technologies) according to the manufacturer’s recommendations (1700V, 20ms, 1 pulse). After transfection, cells were seeded at 1×10^6^ cells/ml for 24 h in culture media. 20 µM of siRNA were used per condition for 10 million cells. RNA interference-mediated gene knockdown was achieved using pre-validated Sigma-Aldrich siRNAs (59>39) for CEBPA#1 (CGGACUUGGUGCGUCUAAG), and CEBPA#2 (GCAACUCUAGUAUUUAGGA), and Control (UAGCAAUGACGAAUGCGUA).

### Flow cytometry-based assay

For lipid peroxide production measurements using C11-BODIPY (581/591) (2 μM) (Invitrogen, Cat# D3861), 2×10^5^ cells were labeled with C11-BODIPY in 1 mL of warm complete medium for 10 min in a tissue culture incubator (37°C, 5% CO_2_) in the dark. Cells were then washed twice and resuspended in 200 μl of fresh PBS.

For neutral lipids uptake measurements using BODIPY FL-C_16_ (505/512) (10 µM) (Invitrogen, Cat# D3821), 2×10^5^ cells were labeled with BODIPY FL-C_16_ in 1 mL of warm complete medium for 2 hours in a tissue culture incubator (37°C, 5% CO_2_) in the dark. Cells were then washed thrice and resuspended in 200 μl of fresh PBS. Data were collected using CytoFLEX flow cytometer and analyzed with FlowJo software (Becton Dickinson).

For neutral lipids measurements using BODIPY (493/503) (0.5µg/ml) (Invitrogen, Cat# D3922), 2×10^5^ were labeled with BODIPY (493/503) in 1 mL of warm complete medium for 20 minutes in a tissue culture incubatoir (37°C, 5% CO_2_) in the dark. Cells were then washed twice and resuspended in 200µL of fresh PBS.

For viability measurement using Annexin-V, 2×10^5^ cells were washed with PBS1X and resuspended in 100 µL of Annexin-V binding buffer (BD Biosciences). 1 µL of Annexin-V-APC (Cat# 550474), FITC (Cat# 556420), BV421 (Cat# 563973) or V500 (Cat# 561501; all from BD Biosciences) were added for 15 min at room temperature in the dark.

For cell death measurements using Propidium iodide (25 μg/mL) (Invitrogen, Cat# P1304MP), 2×10^5^ cells were labeled with Propidium iodide in 1 mL of warm complete medium for 15 minutes in a tissue culture incubator (37°C, 5% CO2) in the dark.

Data were collected using a C6 Accuri flow cytometer (Becton Dickinson) with CFlow Plus software or using CytoFLEX flow cytometer (Beckman Coulter). 10,000 events were captured for subsequent analysis with CFlow Plus software (Becton Dickinson) or with FlowJo software (Becton Dickinson, RRID:SCR_008520).

### Viability assay

AML cells were plated at 20 × 10^4^/ml in 100μl of 10% FBS-supplemented RPMI prior to the addition of compounds. Cells were cultured in the presence of the test compounds for 14 to 96 h at 37°C. Viability was quantified using the fluorescence based Alamar Blue assay (Invitrogen, Cat# DAL1025). Alamar blue was added to each well in 10 μl aliquots. Fluorescence was then measured with a spectramax paradigm from Molecular Devices. Fluorescence values were normalized to DMSO-treated controls for each AML cell line.

### Measurement of synergistic effects

Cell viability was calculated for every dose combination of APR-246 and ferroptosis inducer using the Synergy Finder 2.0 (https://synergyfinder.fimm.fi/)74 and compared to each agent alone. Calculations were based on the ZIP model18. ZIP score interpretation: ZIP<-10, antagonist effect; -10<ZIP<10, additive effect; ZIP>10, synergistic effect.

### Western blots

Cells were lysed in NuPAGE™ LDS Sample Buffer supplemented with NuPAGE™ Sample Reducing Agent (Life Technologies) and heated for 5 min at 90°C. Then proteins were separated using 4–12% gradient polyacrylamide SDS–PAGE gels (Life Technologies) and electrotransferred to 0.2 µm nitrocellulose membranes (GE Healthcare). After blocking in Tris-buffered saline with 0.2% Tween and 5% bovine serum albumin, membranes were blotted overnight at 4 °C with the appropriate primary antibodies. Primary antibodies were detected using the appropriate horseradish peroxidase-conjugated secondary antibodies. Immunoreactive bands were visualized by enhanced chemiluminescence (Thermo-Fisher Scientific, Cat# PI32209) with a Pxi camera (Syngene) using GeneSys software (Syngene, RRID:SCR_015770) or a fusion FX7 edge imaging System (Vilber) using Fusion FX software (Vilber). Protein levels were quantified using GeneTool software (Syngene) or Fusion FX software (Vilber, France) and normalized to nonvariable proteins such as Actin, GAPDH or HSP90. Primary antibodies used were: Actin (Cat# MAB1501, RRID:AB_2223041) was purchased from Millipore; AKT (Cat# 9272, RRID:AB_329827), Phospho-AKT (Ser493) (Cat# 9271, RRID:AB_329825), C/EBPα (Cat# 8178, RRID:AB_11178517), FASN (Cat# 3189, RRID:AB_2100798), FLT3 (Cat# 3462, RRID:AB_2107052), Phospho-FLT3 (Tyr591) (Cat# 3461, RRID:AB_331060), GAPDH (Cat# 5174, RRID:AB_10622025), HSP90 (Cat# 4874, RRID:AB_2121214), p44/p42 MAPK (ERK1/2) (Cat# 9102, RRID:AB_330744), Phospho-p44/42 MAPK (ERK1/2) (Thr202/Tyr204) (Cat# 9101, RRID:AB_331646), STAT5 (Cat# 94205, RRID:AB_2737403) and Phospho-STAT5 (Tyr694) (Cat# 9351, RRID:AB_2315225) were purchased from Cell Signaling Technology; FADS2 (Cat# ab72189, RRID:AB_2041175), GPX4 (Cat# ab41787, RRID:AB_941790), SREBP1 (Cat# ab28481, RRID:AB_778069), SREBP2 (Cat# ab30682, RRID:AB_779079) and SCD (Cat# ab19862, RRID:AB_445179) were purchased from Abcam.

### RNA extraction and Quantitative RT-qPCR

Total RNA was prepared with GenElute Mammalian Total RNA miniprep (Sigma-Aldrich). cDNA was synthesized by Superscript II Reverse Transcriptase (Invitrogen). Genes of interest were quantified using SYBR Green Power UP (Life Technologies). All reactions were carried out in a StepOne instrument (Applied Biosystems) using StepOne software (Applied Biosystems, RRID:SCR_014281). Results were quantified by the ΔCt method using HYWAZ as standard nonvariable genes. Primer sequences used were (5’>3’): *CEBPA*, forward: TATAGGCTGGGCTTCCCCTT, reverse: AGCTTTCTGGTGTGACTCGG; *FASN*, forward: GCAGGAGCTCAAGAAGGTGA, reverse: CCAGGTTGTCCCCTGTGATCC; *SCD*, forward: GGAGCCACCGCTCTTACAAA, reverse: GCCAGGTTTGTAGTACCTCC; *FADS2*, forward: TCCATAAGAACTGGGTGGACCT, reverse: CTCCCAGGATGCCGTAGAAA; *HYWAZ*, forward: ACTTTTGGTACATTGTGGCTTCAA, reverse: CCGCCAGGACAAACCAGTAT.

### Electron microscopy

2×106 cells were fixed in 2% glutaraldehyde in 0.1M sodium phosphate buffer (pH 7.4) for 45 min, then post-fixed for 1.5 hours with 1% osmium tetraoxide and incubated for 12h in 2% uranyl acetate. Cells were then dehydrated by successive washes in a buffer with increasing ethanol concentration (70%, 90%, 100% and 100%), and successive incubation in 100% propylene oxyde, 50-50% propylene oxide/epon and epon. After polymerization, 80-90 nm sections were prepared using an ultracut E microtome (Reichert, Buffalo), stained with 2% uranyl acetate plus Reynold’s lead citrate, and visualized under a Morgagni transmission electron microscope (FEI Compan). All reagents were from Sigma-Aldrich except uranyl acetate (Polysciences, Philadelphia).

### Multi-omic approaches

For lipidomic, C^13^-fluxomic, proteomic, bulk RNAseq, single cell RNAseq analyses of parental MOLM-14 and MV4-11 treated with FLT3i or vehicle, MOLM-14 cells depleted from or overexpressing *CEBPA*, primary patients and PDXs, see **Supplementary Materials and Methods**.

### Statistics

Statistical analyses were conducted using GraphPad Prism software v9.2.0 (RRID:SCR_002798). For *in vitro* studies, statistical significance was determined by the two-tailed unpaired Student’s t test with Welch’s correction, or paired t-test or ratio paired t-test when appropriate. For *in vivo* studies, statistical significance was determined by the non-parametric Mann-Whitney test. Unless otherwise indicated, all data represent the mean ± SD from at least three independent experiments. *p<0.05, **p<0.01, ***p<0.001.

## ACKNOWLEDGMENTS

We thank all members of mice core facilities (UMS006, ANEXPLO, Inserm, Toulouse) in particular Marie Lulka, Christine Campi and Cédric Baudelin for their support and technical assistance. We thank Anne-Marie Benot, Muriel Serthelon and Stéphanie Nevouet for their daily help about the administrative and financial management of our Team. We thank the Geneva University Medical School electron microscopy (PFMU), flow cytometry, bioimaging, genomics (iGE3), proteomics, Reader Assay Development and Screening (READS) and zootechnie core facilities for technical support.

This work was supported by grants from the Laboratoire d’Excellence Toulouse Cancer (TOUCAN and TOUCAN2.0; contract ANR11-LABEX), INCA (PLBIO 2020-010, DIALAML), the Fondation Toulouse Cancer Santé, the Fondation ARC, the Ligue National de Lutte Contre le Cancer and the association GAEL. A.S. is a fellow from the European Regional Development Fund (ERDF) through the Interreg V-A Spain-France-Andorra (POCTEFA) program, project PROTEOblood (EFA360/19). M.S. and R.B. were fellows from the Fondation ARC contre le Cancer. Team JE Sarry is a member of OPALE Carnot Institute, The Organization for Partnerships in Leukemia. Untargeted lipidomics analyses were supported within the MetaboHUB French infrastructure (ANR-INBS-0010). This work was supported by grants from the Dr. Henri Dubois-Ferrière, Dinu Lipatti Fondation and Geneva university hospital private Foundation. This work was also supported by funding from Ligue Genevoise contre le Cancer, Fondation Copley May, Fondation Medic and Fondation Pastré through the Translational Research Center for Hemato-Oncology (University of Geneva, Faculty of Medicine, Geneva, Switzerland).

## REFERENCES

1. Papaemmanuil E, Gerstung M, Bullinger L, et al. Genomic Classification and Prognosis in Acute Myeloid Leukemia. N Engl J Med. 2016;374(23):2209–2221. doi:10.1056/nejmoa1516192

2. Thiede C, Steudel C, Mohr B, et al. Analysis of FLT3-activating mutations in 979 patients with acute myelogenous leukemia: Association with FAB subtypes and identification of subgroups with poor prognosis. Blood. 2002;99(12):4326–4335. doi:10.1182/blood.V99.12.4326

3. Kottaridis PD, Gale RE, Frew ME, et al. The presence of a FLT3 internal tandem duplication in patients with acute myeloid leukemia (AML) adds important prognostic information to cytogenetic risk group and response to the first cycle of chemotherapy: Analysis of 854 patients from the United King. Blood. 2001;98(6):1752–1759. doi:10.1182/blood.V98.6.1752

4. Tiesmeier J, Müller-Tidow C, Westermann A, et al. Evolution of FLT3-ITD and D835 activating point mutations in relapsing acute myeloid leukemia and response to salvage therapy. Leuk Res. 2004;28(10):1069–1074. doi:10.1016/j.leukres.2004.02.009

5. Bjelosevic S, Gruber E, Newbold A, et al. Serine biosynthesis is a metabolic vulnerability in flt3-itd–driven acute myeloid leukemia. Cancer Discov. 2021;11(6):1582–1599. doi:10.1158/2159-8290.CD-20-0738

6. Gregory MA, D’Alessandro A, Alvarez-Calderon F, et al. ATM/G6PD-driven redox metabolism promotes FLT3 inhibitor resistance in acute myeloid leukemia. Proc Natl Acad Sci U S A. 2016;113(43):E6669–E6678. doi:10.1073/pnas.1603876113

7. Ju HQ, Zhan G, Huang A, et al. ITD mutation in FLT3 tyrosine kinase promotes Warburg effect and renders therapeutic sensitivity to glycolytic inhibition. Leukemia. 2017;31(10):2143–2150. doi:10.1038/leu.2017.45

8. Zavorka Thomas ME, Lu X, Talebi Z, et al. Gilteritinib Inhibits Glutamine Uptake and Utilization in FLT3-ITD-positive AML. Mol Cancer Ther. September 2021:molcanther.0071.2021. doi:10.1158/1535-7163.mct-21-0071

9. Gallipoli P, Giotopoulos G, Tzelepis K, et al. Glutaminolysis is a metabolic dependency in FLT3 ITD acute myeloid leukemia unmasked by FLT3 tyrosine kinase inhibition. Blood. 2018;131(15):1639–1653. doi:10.1182/blood-2017-12-820035

10. Stone RM, Mandrekar SJ, Sanford BL, et al. Midostaurin plus Chemotherapy for Acute Myeloid Leukemia with a FLT3 Mutation. N Engl J Med. 2017;377(5):454–464. doi:10.1056/nejmoa1614359

11. Perl AE, Martinelli G, Cortes JE, et al. Gilteritinib or Chemotherapy for Relapsed or Refractory FLT3 -Mutated AML. N Engl J Med. 2019;381(18):1728–1740. doi:10.1056/nejmoa1902688

12. Sato T, Yang X, Knapper S, et al. FLT3 ligand impedes the efficacy of FLT3 inhibitors in vitro and in vivo. Blood. 2011;117(12):3286–3293. doi:10.1182/blood-2010-01-266742

13. Kojima K, McQueen T, Chen Y, et al. p53 activation of mesenchymal stromal cells partially abrogates microenvironment-mediated resistance to FLT3 inhibition in AML through HIF-1α-mediated down-regulation of CXCL12. Blood. 2011;118(16):4431–4439. doi:10.1182/blood-2011-02-334136

14. Traer E, Martinez J, Javidi-Sharifi N, et al. FGF2 from marrow microenvironment promotes resistance to FLT3 inhibitors in acute myeloid leukemia. Cancer Res. 2016;76(22):6471–6482. doi:10.1158/0008-5472.CAN-15-3569

15. Dumas PY, Naudin C, Martin-Lannerée S, et al. Hematopoietic niche drives FLT3-ITD acute myeloid leukemia resistance to quizartinib via STAT5- And hypoxia-dependent upregulation of AXL. Haematologica. 2019;104(10):2017–2027. doi:10.3324/haematol.2018.205385

16. Javidi-Sharifi N, Martinez J, English I, et al. Fgf2-fgfr1 signaling regulates release of leukemia-protective exosomes from bone marrow stromal cells. Elife. 2019;8. doi:10.7554/eLife.40033

17. Joshi SK, Nechiporuk T, Bottomly D, et al. The AML microenvironment catalyzes a stepwise evolution to gilteritinib resistance. Cancer Cell. 2021;39(7):999-1014.e8. doi:10.1016/j.ccell.2021.06.003

18. Smith CC, Levis MJ, Perl AE, et al. Emerging Mutations at Relapse in Patients with FLT3-Mutated Relapsed/Refractory Acute Myeloid Leukemia Who Received Gilteritinib Therapy in the Phase 3 Admiral Trial. Blood. 2019;134(Supplement_1):14–14. doi:10.1182/blood-2019-122620

19. McMahon CM, Ferng T, Canaani J, et al. Clonal selection with RAS pathway activation mediates secondary clinical resistance to selective FLT3 inhibition in acute myeloid leukemia. Cancer Discov. 2019;9(8):1050–1063. doi:10.1158/2159-8290.CD-18-1453

20. Alotaibi AS, Yilmaz M, Kanagal-Shamanna R, et al. Patterns of Resistance Differ in Patients with Acute Myeloid Leukemia Treated with Type I versus Type II FLT3 Inhibitors. Blood Cancer Discov. 2021;2(2):125–134. doi:10.1158/2643-3230.bcd-20-0143

21. Avellino R, Delwel R. Expression and regulation of C/EBPα in normal myelopoiesis and in malignant transformation. Blood. 2017;129(15):2083–2091. doi:10.1182/blood-2016-09-687822

22. Taube F, Georgi JA, Kramer M, et al. CEBPA mutations in 4708 patients with acute myeloid leukemia: differential impact of bZIP and TAD mutations on outcome. Blood. 2022;139(1):87–103. doi:10.1182/blood.2020009680

23. Pabst T, Mueller BU, Harakawa N, et al. AML1-ETO downregulates the granulocytic differentiation factor C/EBPα in t(8;21) myeloid leukemia. Nat Med. 2001;7(4):444–451. doi:10.1038/86515

24. Christy RJ, Yang VW, Ntambi JM, et al. Differentiation-induced gene expression in 3T3-L1 preadipocytes: CCAAT/enhancer binding protein interacts with and activates the promoters of two adipocyte-specific genes. Genes Dev. 1989;3(9):1323–1335. doi:10.1101/gad.3.9.1323

25. Tae HJ, Luo X, Kim KH. Roles of CCAAT/enhancer-binding protein and its binding site on repression and derepression of acetyl-CoA carboxylase gene. J Biol Chem. 1994;269(14):10475–10484. doi:10.1016/s0021-9258(17)34084-x

26. Pedersen TÅ, Bereshchenko O, Garcia-Silva S, et al. Distinct C/EBPα motifs regulate lipogenic and gluconeogenic gene expression in vivo. EMBO J. 2007;26(4):1081–1093. doi:10.1038/sj.emboj.7601563

27. Spiekermann K, Bagrintseva K, Schwab R, Schmieja K, Hiddemann W. Overexpression and constitutive activation of FLT3 induces STAT5 activation in primary acute myeloid leukemia blast cells. Clin Cancer Res. 2003;9(6):2140–2150.

28. Brandts CH, Sargin B, Rode M, et al. Constitutive activation of Akt by Flt3 internal tandem duplications is necessary for increased survival, proliferation, and myeloid transformation. Cancer Res. 2005;65(21):9643–9650. doi:10.1158/0008-5472.CAN-05-0422

29. Chen W, Drakos E, Grammatikakis I, et al. MTOR signaling is activated by FLT3 kinase and promotes survival of FLT3-mutated acute myeloid leukemia cells. Mol Cancer. 2010;9(1):1–7. doi:10.1186/1476-4598-9-292

30. Gregory MA, Nemkov T, Park HJ, et al. Targeting glutamine metabolism and redox state for leukemia therapy. Clin Cancer Res. 2019;25(13):4079–4090. doi:10.1158/1078-0432.CCR-18-3223

31. Guillou H, Zadravec D, Martin PGP, Jacobsson A. The key roles of elongases and desaturases in mammalian fatty acid metabolism: Insights from transgenic mice. Prog Lipid Res. 2010;49(2):186–199. doi:10.1016/j.plipres.2009.12.002

32. Shimano H, Sato R. SREBP-regulated lipid metabolism: Convergent physiology-divergent pathophysiology. Nat Rev Endocrinol. 2017;13(12):710–730. doi:10.1038/nrendo.2017.91

33. Radomska HS, Bassères DS, Zheng R, et al. Block of C/EBPα function by phosphorylation in acute myeloid leukemia with FLT3 activating mutations. J Exp Med. 2006;203(2):371–381. doi:10.1084/jem.20052242

34. Radomska HS, Alberich-Jordà M, Will B, Gonzalez D, Delwel R, Tenen DG. Targeting CDK1 promotes FLT3-activated acute myeloid leukemia differentiation through C/EBPα. J Clin Invest. 2012;122(8):2955–2966. doi:10.1172/JCI43354

35. Tyner JW, Tognon CE, Bottomly D, et al. Functional genomic landscape of acute myeloid leukaemia. Nature. 2018;562(7728):526–531. doi:10.1038/s41586-018-0623-z

36. Network TCGAR. Genomic and Epigenomic Landscapes of Adult De Novo Acute Myeloid Leukemia. N Engl J Med. 2013;368(22):2059–2074. doi:10.1056/nejmoa1301689

37. Verhaak RGW, Wouters BJ, Erpelinck CAJ, et al. Prediction of molecular subtypes in acute myeloid leukemia based on gene expression profiling. Haematologica. 2009;94(1):131–134. doi:10.3324/haematol.13299

38. Léger S, Black WC, Deschenes D, et al. Synthesis and biological activity of a potent and orally bioavailable SCD inhibitor (MF-438). Bioorganic Med Chem Lett. 2010;20(2):499–502. doi:10.1016/j.bmcl.2009.11.111

39. Butler LM, Perone Y, Dehairs J, et al. Lipids and cancer: Emerging roles in pathogenesis, diagnosis and therapeutic intervention. Adv Drug Deliv Rev. 2020;159:245–293. doi:10.1016/j.addr.2020.07.013

40. Jiang X, Stockwell BR, Conrad M. Ferroptosis: mechanisms, biology and role in disease. Nat Rev Mol Cell Biol. 2021;22(4):266–282. doi:10.1038/s41580-020-00324-8

41. Yang WS, Sriramaratnam R, Welsch ME, et al. Regulation of ferroptotic cancer cell death by GPX4. Cell. 2014;156(1-2):317–331. doi:10.1016/j.cell.2013.12.010

42. Flowers MT, Ntambi JM. Role of stearoyl-coenzyme A desaturase in regulating lipid metabolism. Curr Opin Lipidol. 2008;19(3):248–256. doi:10.1097/MOL.0b013e3282f9b54d

43. Hospital MA, Jacquel A, Mazed F, et al. RSK2 is a new Pim2 target with pro-survival functions in FLT3-ITD-positive acute myeloid leukemia. Leukemia. 2018;32(3):597–605. doi:10.1038/leu.2017.284

44. Hayakawa F, Towatari M, Kiyoi H, et al. Tandem-duplicated Flt3 constitutively activates STAT5 and MAP kinase and introduces autonomous cell growth in IL-3-dependent cell lines. Oncogene. 2000;19(5):624–631. doi:10.1038/sj.onc.1203354

45. Tse KF, Novelli E, Civin CI, Bohmer FD, Small D. Inhibition of FLT3-mediated transformation by use of a tyrosine kinase inhibitor. Leukemia. 2001;15(7):1001–1010. doi:10.1038/sj.leu.2402199

46. Sallman DA, DeZern AE, Garcia-Manero G, et al. Eprenetapopt (APR-246) and azacitidine in TP53-mutant myelodysplastic syndromes. In: Journal of Clinical Oncology. Vol 39. J Clin Oncol; 2021:1584–1594. doi:10.1200/JCO.20.02341

47. Tessoulin B, Descamps G, Moreau P, et al. PRIMA-1Met induces myeloma cell death independent of p53 by impairing the GSH/ROS balance. Blood. 2014;124(10):1626–1636. doi:10.1182/blood-2014-01-548800

48. Bykov VJN, Zhang Q, Zhang M, Ceder S, Abrahmsen L, Wiman KG. Targeting of mutant P53 and the cellular redox balance by APR-246 as a strategy for efficient cancer therapy. Front Oncol. 2016;6(FEB):21. doi:10.3389/fonc.2016.00021

49. Liu DS, Duong CP, Haupt S, et al. Inhibiting the system xC-/glutathione axis selectively targets cancers with mutant-p53 accumulation. Nat Commun. 2017;8. doi:10.1038/ncomms14844

50. Birsen R, Larrue C, Decroocq J, et al. APR-246 induces early cell death by ferroptosis in acute myeloid leukemia. Haematologica. 2022;107(2):403–416. doi:10.3324/haematol.2020.259531

51. Fujihara KM, Benitez MC, Cabalag CS, et al. SLC7A11 Is a Superior Determinant of APR-246 (Eprenetapopt) Response than TP53 Mutation Status. Mol Cancer Ther. 2021;20(10):1858–1867. doi:10.1158/1535-7163.MCT-21-0067

52. Burchert A, Bug G, Fritz L V., et al. Sorafenib maintenance after allogeneic hematopoietic stem cell transplantation for acute myeloid leukemia with FLT3-internal tandem duplication mutation (SORMAIN). J Clin Oncol. 2020;38(26):2993–3002. doi:10.1200/JCO.19.03345

53. Smith CC, Wang Q, Chin C-S, et al. Validation of ITD mutations in FLT3 as a therapeutic target in human acute myeloid leukaemia. Nat 2012 4857397. 2012;485(7397):260–263. doi:10.1038/nature11016

54. Green AS, Maciel TT, Hospital MA, et al. Pim kinases modulate resistance to FLT3 tyrosine kinase inhibitors in FLT3-ITD acute myeloid leukemia. Sci Adv. 2015;1(8). doi:10.1126/sciadv.1500221

55. Stockard B, Garrett T, Guingab-Cagmat J, Meshinchi S, Lamba J. Distinct Metabolic features differentiating FLT3-ITD AML from FLT3-WT childhood acute myeloid leukemia. Sci Rep. 2018;8(1). doi:10.1038/s41598-018-23863-9

56. Ogretmen B. Sphingolipid metabolism in cancer signalling and therapy. Nat Rev Cancer. 2017;18(1):33–50. doi:10.1038/nrc.2017.96

57. Dany M, Gencer S, Nganga R, et al. Targeting FLT3-ITD signaling mediates ceramide-dependent mitophagy and attenuates drug resistance in AML. Blood. 2016;128(15):1944–1958. doi:10.1182/blood-2016-04-708750

58. Gouazé V, Mirault ME, Carpentier S, Salvayre R, Levade T, Andrieu-Abadie N. Glutathione peroxidase-1 overexpression prevents ceramide production and partially inhibits apoptosis in doxorubicin-treated human breast carcinoma cells. Mol Pharmacol. 2001;60(3):488–496. https://pubmed.ncbi.nlm.nih.gov/11502879/. Accessed April 15, 2022.

59. Hermetet F, Mshaik R, Simonet J, Callier P, Delva L, Quéré R. High-fat diet intensifies MLL-AF9-induced acute myeloid leukemia through activation of the FLT3 signaling in mouse primitive hematopoietic cells. Sci Rep. 2020;10(1). doi:10.1038/s41598-020-73020-4

60. Stuani L, Sabatier M, Saland E, et al. Mitochondrial metabolism supports resistance to IDH mutant inhibitors in acute myeloid leukemia. J Exp Med. 2021;218(5). doi:10.1084/jem.20200924

61. Puca F, Yu F, Bartolacci C, et al. Medium-chain acyl-coa dehydrogenase protects mitochondria from lipid peroxidation in glioblastoma. Cancer Discov. 2021;11(11):2904–2923. doi:10.1158/2159-8290.CD-20-1437

62. Mbah NE, Lyssiotis CA. Metabolic regulation of ferroptosis in the tumor microenvironment. J Biol Chem. 2022;298(3):101617. doi:10.1016/j.jbc.2022.101617

63. Heydt Q, Larrue C, Saland E, et al. Oncogenic FLT3-ITD supports autophagy via ATF4 in acute myeloid leukemia. Oncogene. 2018;37(6):787–797. doi:10.1038/onc.2017.376

64. Rysman E, Brusselmans K, Scheys K, et al. De novo lipogenesis protects cancer cells from free radicals and chemotherapeutics by promoting membrane lipid saturation. Cancer Res. 2010;70(20):8117–8126. doi:10.1158/0008-5472.CAN-09-3871

65. de Groot S, Lugtenberg RT, Cohen D, et al. Fasting mimicking diet as an adjunct to neoadjuvant chemotherapy for breast cancer in the multicentre randomized phase 2 DIRECT trial. Nat Commun. 2020;11(1). doi:10.1038/s41467-020-16138-3

66. Lien EC, Westermark AM, Zhang Y, et al. Low glycaemic diets alter lipid metabolism to influence tumour growth. Nature. 2021;599(7884):302–307. doi:10.1038/s41586-021-04049-2

67. Chen X, Kang R, Kroemer G, Tang D. Broadening horizons: the role of ferroptosis in cancer. Nat Rev Clin Oncol. 2021;18(5):280–296. doi:10.1038/s41571-020-00462-0

68. Lei G, Zhuang L, Gan B. Targeting ferroptosis as a vulnerability in cancer. Nat Rev Cancer. 2022. doi:10.1038/s41568-022-00459-0

69. Stockwell BR, Friedmann Angeli JP, Bayir H, et al. Ferroptosis: A Regulated Cell Death Nexus Linking Metabolism, Redox Biology, and Disease. Cell. 2017;171(2):273–285. doi:10.1016/j.cell.2017.09.021

70. Cluzeau T, Sebert M, Rahmé R, et al. Eprenetapopt plus azacitidine in TP53-mutated myelodysplastic syndromes and acute myeloid Leukemia: A phase II study by the groupe francophone des Myélodysplasies (GFM). In: Journal of Clinical Oncology. Vol 39. J Clin Oncol; 2021:1575–1583. doi:10.1200/JCO.20.02342

71. Tesfay L, Paul BT, Konstorum A, et al. Stearoyl-CoA desaturase 1 protects ovarian cancer cells from ferroptotic cell death. Cancer Res. 2019;79(20):5355–5366. doi:10.1158/0008-5472.CAN-19-0369

72. Meerbrey KL, Hu G, Kessler JD, et al. The pINDUCER lentiviral toolkit for inducible RNA interference in vitro and in vivo. Proc Natl Acad Sci U S A. 2011;108(9):3665–3670. doi:10.1073/pnas.1019736108

73. Campeau E, Ruhl VE, Rodier F, et al. A versatile viral system for expression and depletion of proteins in mammalian cells. PLoS One. 2009;4(8). doi:10.1371/journal.pone.0006529

74. Ianevski A, Giri AK, Aittokallio T. SynergyFinder 2.0: visual analytics of multi-drug combination synergies. Nucleic Acids Res. 2020;48(W1):W488–W493. doi:10.1093/NAR/GKAA216

