## Supplementary data for "C/EBPα confers dependence to fatty acid anabolic pathways and vulnerability to lipid oxidative stress in FLT3-mutant leukemia"

*Sabatier, Birsen et al.*

Table of Content:

- Supplementary Material and Methods
- Supplementary Figure 1-17
- Supplementary Table 1-10 (see attached)

**Supplementary Material and Methods**

**Lipidomic analysis of parental MOLM-14 and MV4-11**

*Sample preparation for untargeted lipidomics:*

Cell lipid extracts were obtained from a frozen pellet of $4.10^{+6}$ harvested cells. Samples were resuspended in 70µL of ultrapure water, and then sonicated 5 times for 10 s using a sonication probe (vibra cell). At this step, 20µL of each sample were withdrawn for further determining the total protein concentration (colorimetric quantification / Pierce BCA Protein Assay Kit, Thermo Fisher Scientific). Samples were extracted using an adapted method previously described[1]. Briefly, volumes of CHCL3/MeOH 2:1 (v/v) were adjusted depending total protein concentration at 3.5µg/µl in addition to internal standards. Samples were vortexed for 60 s, sonicated for 30 s using an ultrasonic probe (Bioblock Scientific Vibra Cell VC 75,185, Thermo Fisher Scientific) and incubated for 2 h at 4 °C with mixing. According to volumes of CHCL3/MeOH 2:1 (v/v), H2O was then added and samples were vortexed for 60 s before centrifugation at 15,000 g for 15 min at 4 °C. The upper phase (aqueous phase), containing gangliosides, lysoglycerophospholipids, and short chain glycerophospholipids, was transferred into a glass tube and dried under a stream of nitrogen. The protein disk interphase was discarded and the lower lipid-rich phase (organic phase) was pooled with the dried upper phase and the mixture dried under nitrogen. All samples were resuspended in the same initial volume of CHCL3/MeOH 2:1 (v/v) and 40 µl of each extract was 5-fold diluted in a solution of MeOH/isopropanol/H2O 65:35:5 (v/v/v) before injection. A quality control (QC) sample was obtained by pooling 20 µL of each sample preparation. QC samples was injected every 10 samples in order to evaluate the signal variations of lipid species.

*Untargeted lipidomics and data processing*

Lipidomic profiles were determined using an Ultimate 3000 liquid chromatography system (Thermo Fisher Scientific) coupled to a high resolution Thermo Orbitrap Fusion (Thermo Fisher Scientific) equipped with an electrospray source (ESI). Chromatographic separation was performed on a Phenomenex Kinetex C8 column (150 x 2.1 mm, 2.6 µm) at 0.4 mL/min, 60 °C and using an injection volume of 10 µL. Mobile phases A and B were H2O/ MeOH 60:40 (v/v), 0.1% formic acid and isopropanol/MeOH 90:10 (v/ v), 0.1% formic acid in negative ionization mode, respectively. Ammonium formate (10 mM) was added to both mobile phases in the positive ionization mode in order to detect glycerolipids and cholesteryl-esters under [M+NH4] + form. The gradient elution was solvent B was maintained for 2.5 min at 32%, from 2.5 to 3.5 min it was increased to 45% B, from 3.5 to 5 min to 52% B, from 5 to 7 min to 58% B, from 7 to 10 min to 66% B, from 10 to 12 min to 70% B, from 12 to 15 min to 75% B, from 15 to 19 min to 80% B, from 19 to 22 min to 85% B, and from 22 to 23 min to 95% B; from 23 to 25 min, 95% B was maintained; from 25 to 26 min solvent B was decreased to 32% and then maintained for 4 min for column re-equilibration. The mass resolving power of the mass spectrometer was 240,000 (FWHM) for MS experiments. Samples were analyzed in both positive and negative ionization modes. The ESI source parameters were as follows: the spray voltage was set to 3.7 kV and -3.2 kV in positive and negative ionization mode, respectively. The heated capillary was kept at 360 °C and the sheath and auxiliary gas flow were set to 50 and 15 (arbitrary units), respectively. Mass spectra were recorded in full-scan MS mode from m/z 50 to m/z 2000. After LC-HRMS analysis of samples and annotation of features, QC samples were re-injected for higher energy collisional dissociation (HCD) MS/MS experiments in positive and negative ion modes on the same instrument set in targeted mode using inclusion lists. The isolation width was set at m/z 0.8, the stepped normalized collision energy was set at 20 % ± 10 % and the mass resolution was set at 17 500 FWHM at m/z 200. HCD mass spectra were inspected manually in order to confirm annotations.

Obtained Raw files were converted to the mzML or mzXML format with the ProteoWizard software[2]. Mass spectra were processed using the xcms R package deployed in the Workflow4Metabolomics Galaxy platform[3] (https://workflow4metabolomics.org). Features were annotated by using an in-house database based on accurate measured masses and several filtration steps such as retention time windows depending lipid classes and relative isotopic abundance (RIA) of lipid species in order to limit false annotations for lipidomics.

**Lipidomic analysis of MOLM-14 cells depleted from or overexpressing *CEBPA***

*Lipid extraction*

An amount of cells containing 10 µg of DNA was homogenized in 700 µL of water with a handheld sonicator and was mixed with 800 µl HCl(1M):CH3OH 1:8 (v/v), 900 µl CHCl3, 200 µg/ml of the antioxidant 2,6-di-tert-butyl-4-methylphenol (BHT, Sigma-Aldrich, Cat# B1378) and 3 µl of SPLASH® LIPIDOMIX® Mass Spec Standard (Avanti Polar Lipids, Cat# 330707). After vortexing and centrifugation, the lower organic fraction was collected and evaporated using a Savant Speedvac spd111v (Thermo Fisher Scientific) at room temperature and the remaining lipid pellet was stored at -20°C under argon.

*Mass spectrometry*

Just before mass spectrometry analysis, lipid pellets were reconstituted in 100% ethanol. Lipid species were analyzed by liquid chromatography electrospray ionization tandem mass spectrometry (LC-ESI/MS/MS) on a Nexera X2 UHPLC system (Shimadzu) coupled with hybrid triple quadrupole/linear ion trap mass spectrometer (6500+ QTRAP system; AB SCIEX).

Chromatographic separation was performed on a XBridge amide column (150 mm × 4.6 mm, 3.5 µm; Waters) maintained at 35°C using mobile phase A [1 mM ammonium acetate in water-acetonitrile 5:95 (v/v)] and mobile phase B [1 mM ammonium acetate in water-acetonitrile 50:50 (v/v)] in the following gradient: (0-6 min: 0% B → 6% B; 6-10 min: 6% B → 25% B; 10-11 min: 25% B → 98% B; 11-13 min: 98% B → 100% B; 13-19 min: 100% B; 19-24 min: 0% B) at a flow rate of 0.7 mL/min which was increased to 1.5 mL/min from 13 minutes onwards. SM, CE, CER, DCER, HCER, LCER were measured in positive ion mode with a precursor scan of 184.1, 369.4, 264.4, 266.4, 264.4 and 264.4 respectively. TAG, DAG and MAG were measured in positive ion mode with a neutral loss scan for one of the fatty acyl moieties. PC, LPC, PE, LPE, PG, PI and PS were measured in negative ion mode by fatty acyl fragment ions. Lipid quantification was performed by scheduled multiple reactions monitoring (MRM), the transitions being based on the neutral losses or the typical product ions as described above. The instrument parameters were as follows: Curtain Gas = 35 psi; Collision Gas = 8 a.u. (medium); IonSpray Voltage = 5500 V and −4,500 V; Temperature = 550°C; Ion Source Gas 1 = 50 psi; Ion Source Gas 2 = 60 psi; Declustering Potential = 60 V and −80 V; Entrance Potential = 10 V and −10 V; Collision Cell Exit Potential = 15 V and −15 V.

The following fatty acyl moieties were taken into account for the lipidomic analysis: 14:0, 14:1, 16:0, 16:1, 16:2, 18:0, 18:1, 18:2, 18:3, 20:0, 20:1, 20:2, 20:3, 20:4, 20:5, 22:0, 22:1, 22:2, 22:4, 22:5 and 22:6 except for TGs which considered: 16:0, 16:1, 18:0, 18:1, 18:2, 18:3, 20:3, 20:4, 20:5, 22:2, 22:3, 22:4, 22:5, 22:6.

*Data Analysis*

Peak integration was performed with the MultiQuantTM software version 3.0.3. Lipid species signals were corrected for isotopic contributions (calculated with Python Molmass 2019.1.1) and were quantified based on internal standard signals and adheres to the guidelines of the Lipidomics Standards Initiative (LSI) (level 2 type quantification as defined by the LSI). Unpaired t-test p-values was calculated in Python StatsModels version 0.10.1. Lipid species were considered to have significantly changed in abundance with an FDR ≤ 0.05 and an absolute fold change FC≥ |1.5| (log2FC ≥ |0.58|). Lipid species were excluded when more than one replicate was impaired per condition. If only one replicate was impaired, it was excluded and the lipid species conserved. PUFA:MUFA ratios for each phospholipid subclasses were calculated using this formula:

$$\frac{PUFA}{MUFA} = \frac{\Sigma\ (relative\ quantity\ of\ PUFA\ from\ 2\ to\ 6\ unsaturations)}{relative\ quantity\ of\ MUFA}$$

**Fatty acids isotopic labeling profiling**

*Sample preparation*

MV4-11 and MOLM-14 cells were plated at $0.4x10^6$ cells per mL and treated with QUIZ in media supplemented with U-$^{13}C_6$-glucose (5.6 mM) or U-$^{13}C_5$-glutamine (2 mM) for 24 h before sampling. Lipids corresponding total FA from cell pellets of three million cells of three independent experiments were extracted according to Bligh and Dyer[4] in dichloromethane/water/methanol 2% acetic acid (2.5:2.5:2, v/v/v), in the presence of the internal standards glyceryl trinonadecanoate (4µg). After centrifugation during 6min at 2500 rpm, the lipid extract was hydrolysed in KOH (0.5 M in methanol, 1mL) at 55°C for 30 min. A second extraction was performed with dichloromethane/water/methanol (2.5/2/1.5, v/v/v). The lipid extract was evaporated to dryness and dissolved in ethyl acetate (10 µL). Derivatization was performed with PentafluoroBenzylBrommide in ACN (1%) and Diisopropylethylamine in ACN (1%) (1/1, v/v).

*GC-MS Analysis*

Fatty acid methyl ester (1 µL) were analyzed on a gas chromatography Thermo Trace 1310 (Thermo Fisher Scientific) with mass spectrometer Thermo TSQ 8000 EVO (Thermo Fisher Scientific). Chemical ionization is used as ionization, helium as carrier gas, and a HP-5MS column (30 m x 0.25 mm, 0.25 µm) is used to separation. Oven temperature was programmed from 130°C to 232°C at a rate of 4°C/min (8 min), from 232°C to 230°C at a rate of 10°C/min (2 min). A full scan method was used to detected compounds.

*Data Processing*

GC-MS analysis produced a mass spectrum for each FA, which contains the abundance of each isotopologue. For each FA, the lightest (unlabeled) isotopologue is denoted M + 0 whereas the isotopologue with 1 atom [$^{13}$C] PFB-palmitate (M+1) has a mass of 256.3, etc. Isotopic clusters were obtained by integrating gas chromatographic signals for each isotopologue. Isotopologue distributions were obtained from the corresponding isotopic clusters after correction for natural abundance of carbon and non-tracer elements (oxygen and hydrogen) using IsoCor software[5] (https://isocor.readthedocs.io/en/latest/index.html), and purity of the tracer was corrected assuming 99% $^{13}$C-purity. Finally, the $^{13}$C-enrichment, which represents the mean content in tracer atoms ($^{13}$C) within the molecule, was calculated from the corresponding IDs.

**Proteomics**

*Sample preparation*

Cell pellets were resuspended in 100 µl of 0.1 % RapiGest Surfactant (Waters) in 0.1M triethylammonium bicarbonate (TEAB). Samples were heated for 5 min at 95°C. Lysis was performed by sonication (6 x 30 sec.) at 70% amplitude and 0.5 pulse. Samples were kept 30 sec. on ice between each cycle of sonication. Samples were centrifuged for 5 min. at 17’000 x g. Remaining pellet was solubilized in 100 µl of 0.1 % RapiGest, with 2 µl of 0.1M MgCl2 and 0.2 µl of benzonase for DNA digestion. Protein lysates were pooled together and protein concentration was measured by Bradford assay and 25 µg of each sample was subjected to protein digestion as follow: sample volume was adjusted to 100 µl with 0.1M TEAB to obtain a final concentration of RapiGest 0.1%. 2 µl of Dithioerythritol (DTE) 50 mM in distilled water were added and the reduction was carried out at 60°C for 1h. Alkylation was performed by adding 2 µl of iodoacetamide (400 mM in distilled water) during 1 hour at room temperature in the dark. Overnight digestion was performed at 37 °C with 5 µL of freshly prepared trypsin (Promega; 0.1 µg/µl in TEAB 0.1M). To remove RapiGest, samples were acidified with TFA, heated at 37°C for 45 min. and centrifuged 10 min. at 17’000 x g. Supernatants were then desalted with a C18 microspin column (Harvard Apparatus) according to manufacturer’s instructions, completely dried under speed-vacuum and stored at -20°C.

*ESI-LC-MSMS*

Samples were diluted in 25 µl of loading buffer (5% CH3CN, 0.1% FA). Biognosys iRT peptides (1.5 µl) were added to each sample and 2 µl were injected on column. LC-ESI-MS/MS was performed on an Orbitrap Fusion Lumos Tribrid mass spectrometer (Thermo Fisher Scientific) equipped with an Easy nLC1200 liquid chromatography system (Thermo Fisher Scientific). Peptides were trapped on a Acclaim pepmap100, C18, 3µm, 75µm x 20mm nano trap-column (Thermo Fisher Scientific) and separated on a 75 µm x 500 mm, C18 ReproSil-Pur (Dr. Maisch GmBH), 1.9 µm, 100 Å, home-made column. The analytical separation was run for 135 min using a gradient of H2O/FA 99.9%/0.1% (solvent A) and CH3CN/H2O/FA 80.0%/19.9%/0.1% (solvent B). Data-Independant Acquisition (DIA) was performed with MS1 full scan at a resolution of 60,000 (FWHM) followed by 30 DIA MS2 scan with variable windows. MS1 was performed in the Orbitrap with an AGC target of 1 x $10^6$, a maximum injection time of 50 ms and a scan range from 400 to 1240 m/z. DIA MS2 was performed in the Orbitrap using higher-energy collisional dissociation (HCD) at 30%. Isolation windows was set to 28 m/z with an AGC target of 1 x 106 and a maximum injection time of 54 ms.

*Data analysis*

DIA raw files were loaded into Spectronaut v.14 (Biognosys) and analysed by directDIA using default settings. Briefly, data were searched against Human reference proteome fasta database (Uniprot, release 2020_09, 55471 entries). Trypsin was selected as the enzyme, with one potential missed cleavage. Variable amino acid modifications were oxidized methionine and deaminated (NQ). Fixed amino acid modification was carbamidomethyl cysteine. Both peptide precursor and protein FDR were controlled at 1% (Q value < 0.01). Single Hit Proteins were excluded. For quantitation, Top 3 precursor area per peptides were used, "only protein group specific" was selected as proteotypicity filter and normalization was set to "global normalization". The quantitative analysis was performed with MapDIA tool, using the precursor quantities extracted from Spectronaut output. No further normalization was applied. The following parameters were used: min peptides = 2, max peptides = 10, min correl = -1, Min_DE = 0.01, max_DE = 0.99, and experimental_design = replicate design. Proteins were considered to have significantly changed in abundance with an FDR ≤ 0.05 and an absolute fold change FC≥ |1.5| (log2FC ≥ |0.58|). FDR q-value was calculated using the two-stage step-up method of Benjamini, Krieger and Yekutieli (Graphpad Prism, RRID:SCR_002798). Proteins were excluded when more than one replicate was impaired per condition. If only one replicate was impaired, it was excluded and the protein conserved.

**Gene expression profiling**

*Gene chip hybridization*

RNA was extracted using a RNeasy Mini Kit (Qiagen, Redwood City, CA, USA) and quality was evaluated with a Bioanalyzer 2100 (using Agilent RNA6000 nano chip kit), and 100 ng of total RNA was reverse transcribed using the GeneChip® WT Plus Reagent Kit according to the manufacturer's instructions (Affymetrix, Thermo Fischer Scientific). Briefly, Double strand cDNA was used for in vitro transcription with T7 RNA polymerase and 5.5mg of Sens Target DNA was fragmented and labelled with biotin. cDNA were then hybridized to GeneChip® Clariom S Human (Affymetrix) at 45°C for 17 hours, then washed on the fluidic station FS450 (Affymetrix), and scanned using the GCS3000 7G (Thermo Fischer Scientific). Scanned images were then analyzed with Expression Console software (Affymetrix, Thermo Fischer Scientific) to obtain raw data (.cel files) and metrics for quality controls.

*Data processing and statistics*

Raw fluorescence intensity values were normalized using Robust Multiarray Average (RMA) algorithm in R to generate the normalized data matrix by performing background correction, quantile normalization and log2 transformation of raw fluorescence intensity values of each gene. All quality controls and statistics were performed using Partek® Genomics Suite software (Partek, St. Louis, MO, USA). Data were normalized using custom brainarray CDF files (v20 ENTREZG). To identify differentially expressed genes, we applied a classical analysis of variance (ANOVA) with a FDR permutation-base for each gene. We created a matrix with only the significant ANOVA site and performed Z-scoring of rows. Hierarchical clustering by Pearson's dissimilarity and average linkage and principal components analysis (PCA) were conducted in an unsupervised fashion to control for experimental bias or outlier samples. We set a filter for those genes that displayed at least a ≥1,5 or ≤-1,5 fold difference in expression between groups and achieved an FDR of <0.05. Data were then interrogated for evidence of biologic pathway dysregulation using EnrichR userfriendly website (https://maayanlab.cloud/Enrichr/)[6–8] and Gene set enrichment analysis (GSEA, Broad Institute). Enrichment rates were considered as significant for *p*-value<0.05

### GSEA analysis

GSEA analysis was performed using GSEA version 4.1.0 (https://www.gsea-msigdb.org/gsea/index.jsp, BroadInstitute, RRID:SCR_003199)[9,10]. Gene signatures used in this study were from Broad Institute database, literature, or in-house built. Following parameters were used: Number of permutations = 1000, permutation type = gene_set. Other parameters were left at default values.

### RNA-sequencing

*Library preparation*

RNA sequencing library preparation was performed using Illumina TruSeq stranded mRNA protocol. The average number of per sample reads was 47 million. We mapped the reads to the human genome reference "hg38_chr_only_and_herpes.fa" using STAR aligner (STAR v2.6.1a). Base-calling accuracy scores (Phred quality score, Q30) were obtained from demultiplexer bcl2fastq v2.20. Detailed methods for RNA-seq quantification and differential gene expression analysis are provided in the Supplemental Material.

*RNA-seq quantification*

Paired-end reads were aligned to the human reference transcriptome "Homo_sapiens.GRCh38.cdna.all.fa.gz" using Salmon 1.3.0. For mapping, we used salmon quant command with parameters like –gcBias, --useVBOpt, --seqBias, and –validateMappings, correcting specific biases in the sequenced reads and increasing the sensitivity and specificity of the mapping. After mapping, R (R version R version 4.0.2) packages tximport (version 1.14.0) and DESeq2 (version 1.26.0) were used to summarize transcript-level estimate to the gene-level estimates. Further, we used the raw gene-level abundance estimates from salmon to perform to identify differentially expressed genes.

*Differential gene expression analysis*

Differential gene expression analysis was performed using an R Bioconductor software package edgeR 4. First, we filtered the genes with a meager count using an in-built function in edgeR filterByExpr (), and the library size was recomputed. Following this, the normalization factors were estimated using the calcNormFactors () function, and the per gene dispersion was assessed using the estimateDisp () function in edgeR. Here, a design formula (linear model, ~Rasopathy) was provided for dispersion calculation. Finally, the negative binomial generalized linear model was fit using glmFi t() function, and likelihood ratio test glmLRT () was used to test for differential expression. We assessed the differentially expressed genes between the Rasopathy-Yes vs. Rasopathy-No conditions. The topTags function was used to extract the top-ranked genes, and the results were presented as a volcano plot using an R Bioconductor package EnhancedVolcano. We also performed gene set enrichment analysis 5 using the oncogenic signature gene set in which we manually added several gene sets from recent literature.

Bulk RNAseq data from patients was collected in two batches, therefore a batch effect correction procedure was required. We applied the function removeBatchEffect from the R package limma[11]. The patient id was used as the batch parameter in calling the function, therefore removing at the same time the batch effect and the individual patient variance from the data.

### Single cell RNA-sequencing

*Purification of hCD45+CD33+ from PDX*

After red blood cells lysis with EL buffer (Qiagen, 79217), $10x10^6$ cells from mice bone marrow of PDXs treated with vehicle or GILT were stained with Fixable Viability Stain 510 (BD Biosciences Cat# 564406, RRID:AB_2869572) and the fluorescent conjugated antibodies hCD45-APC (BD Biosciences Cat# 555485, RRID:AB_398600) and hCD33-PE (BD Biosciences Cat# 555450, RRID:AB_395843) in cell staining buffer (Biolegend, BLE420201) following to the manufacturer's instructions. Viable hCD45+CD33+ were sorted using BD FACSAria Fusion flow cytometer (Becton Dickinson) and gently re-suspended at 1500cells/µL in PBS1X supplemented with 0.04% Ultrapure BSA (Invitrogen, Cat# AM2616).

*Single cell library preparation and sequencing*

Up to 10,000 cells per sample were encapsulated using the 10X Chromium controller (10X Genomics) following the Chromium Next GEM Single Cell 3' User Guide v3.1 (10X Genomics). Briefly, cells were encapsulated into oil droplets with barcoded Gel Beads and reagents to convert mRNA into complementary DNA (cDNA). cDNAs were then amplified, fragmented and Illumina adapters were added during ligation. After performing a last index PCR, libraries quality was assessed using Tapestation 2200 (Agilent) and frozen before quantitation and sequencing. Barcoding was done on three levels: cell barcodes allow attribution of each sequence read to its cell of origin; unique molecular identifiers (UMIs) upstream to poly(d)T primers allow tagging of each original molecule to avoid amplification bias; and index allows pooling of different samples. scRNA-seq libraries were quantified with HSD1000 reagents on TapeStation 2200 system (Agilent) and pooled at equimolar concentrations within 200 to 700bp. Pool libraries were sequenced at 1.8pM 1% phiX using a NextSeq 500 sequencer (Illumina), at a sequencing depth of 40,000-50,000 reads per cell. Sequencing were configured Paired-End at 28 cycles for the first read, 91 cycles for the second one and 8 cycles for a single index.

*Data analysis*

Single cell RNA sequencing output bcl2 files were converted to FASTQ format by using Cell Ranger v.6.1.1.1[12]. Count matrices from Cell Ranger were imported with the Seurat package v4.1.0[13] into the R programming environment v4.1.2. Cells that were deemed of lower quality were removed, based on their number of unique genes (<200 or >7500 unique genes) and the proportion of mitochondrial RNA (>20%). Gene counts were normalized with a regularized negative binomial regression, using the SCT normalization workflow from the Seurat package[14]. In order to compare gene expression and signatures values after GILT treatment with those before treatment, the datasets from pre and post GILT treatment from each of the PDX sample were integrated following the Seurat dataset integration protocol[15]. For dataset integration we used 30 principal components and 3000 genes. We then applied dimensional reduction where the 30 principal components were further summarized using UMAP dimensionality reduction. Cell clusters were identified using Louvain clustering within the Seurat workflow, the resolution parameter was set to 0.25. We identified clustering of co-expressing genes by hierarchical cluster with the package nclust (https://gitlab.com/pwirapati/nclust). We computed the values of the signatures on SCT transformed data with the AddModuleScore method from Seurat[16]. The signature values were compared with UMAP and violin plots.

### References


1. Seyer A, Boudah S, Broudin S, Junot C, Colsch B. Annotation of the human cerebrospinal fluid lipidome using high resolution mass spectrometry and a dedicated data processing workflow. *Metabolomics*. 2016;12(5). doi:10.1007/s11306-016-1023-8
2. Kessner D, Chambers M, Burke R, Agus D, Mallick P. ProteoWizard: Open source software for rapid proteomics tools development. *Bioinformatics*. 2008;24(21):2534-2536. doi:10.1093/bioinformatics/btn323
3. Giacomoni F, Le Corguillé G, Monsoor M, et al. Workflow4Metabolomics: A collaborative research infrastructure for computational metabolomics. *Bioinformatics*. 2015;31(9):1493-1495. doi:10.1093/bioinformatics/btu813
4. Bligh EG, Dyer WJ. A rapid method of total lipid extraction and purification. *Can J Biochem Physiol*. 1959;37(8):911-917. doi:10.1139/o59-099
5. Millard P, Delépine B, Guionnet M, Heuillet M, Bellvert F, Létisse F. IsoCor: Isotope

correction for high-resolution MS labeling experiments. *Bioinformatics*. 2019;35(21):4484-4487. doi:10.1093/bioinformatics/btz209

6. Chen EY, Tan CM, Kou Y, et al. Enrichr: Interactive and collaborative HTML5 gene list enrichment analysis tool. *BMC Bioinformatics*. 2013;14. doi:10.1186/1471-2105-14-128

7. Kuleshov M V., Jones MR, Rouillard AD, et al. Enrichr: a comprehensive gene set enrichment analysis web server 2016 update. *Nucleic Acids Res*. 2016;44(W1):W90-W97. doi:10.1093/nar/gkw377

8. Xie Z, Bailey A, Kuleshov M V., et al. Gene Set Knowledge Discovery with Enrichr. *Curr Protoc*. 2021;1(3):e90. doi:10.1002/cpz1.90

9. Subramanian A, Tamayo P, Mootha VK, et al. Gene set enrichment analysis: A knowledge-based approach for interpreting genome-wide expression profiles. *Proc Natl Acad Sci U S A*. 2005;102(43):15545-15550. doi:10.1073/pnas.0506580102

10. Mootha VK, Lindgren CM, Eriksson KF, et al. PGC-1α-responsive genes involved in oxidative phosphorylation are coordinately downregulated in human diabetes. *Nat Genet*. 2003;34(3):267-273. doi:10.1038/ng1180

11. Ritchie ME, Phipson B, Wu D, et al. Limma powers differential expression analyses for RNA-sequencing and microarray studies. *Nucleic Acids Res*. 2015;43(7):e47. doi:10.1093/nar/gkv007

12. Zheng GXY, Terry JM, Belgrader P, et al. Massively parallel digital transcriptional profiling of single cells. *Nat Commun*. 2017;8. doi:10.1038/ncomms14049

13. Hao Y, Hao S, Andersen-Nissen E, et al. Integrated analysis of multimodal single-cell data. *Cell*. 2021;184(13):3573-3587.e29. doi:10.1016/j.cell.2021.04.048

14. Hafemeister C, Satija R. Normalization and variance stabilization of single-cell RNA-seq data using regularized negative binomial regression. *Genome Biol*. 2019;20(1). doi:10.1186/s13059-019-1874-1

15. Stuart T, Butler A, Hoffman P, et al. Comprehensive Integration of Single-Cell Data. *Cell*. 2019;177(7):1888-1902.e21. doi:10.1016/j.cell.2019.05.031

16. Tirosh I, Izar B, Prakadan SM, et al. Dissecting the multicellular ecosystem of metastatic melanoma by single-cell RNA-seq. *Science (80- )*. 2016;352(6282):189-196. doi:10.1126/science.aad0501

### SUPPLEMENTAL FIGURES

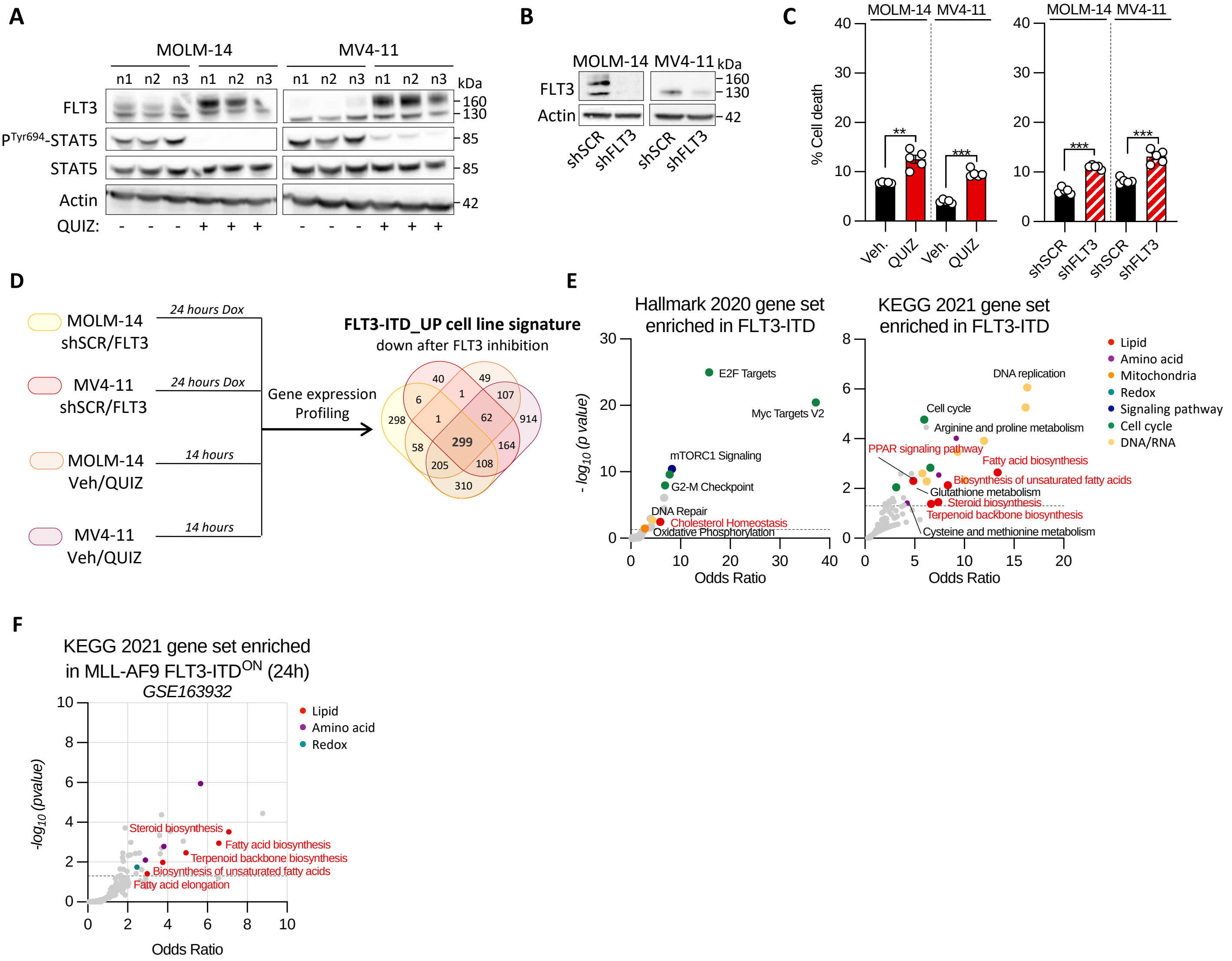
A
MOLM-14
MV4-11
n1 n2 n3 n1 n2 n3
n1 n2 n3 n1 n2 n3
kDa
FLT3
PTyr694-STAT5
STAT5
Actin
QUIZ:
- - - + + +
- - - + + +
160
130
85
85
42
B
MOLM-14 MV4-11
kDa
FLT3
Actin
160
130
42
shSCR
shFLT3
shSCR
shFLT3
C
MOLM-14 MV4-11
% Cell death
**
***
***
***
Veh.
QUIZ
Veh.
QUIZ
shSCR
shFLT3
shSCR
shFLT3
D
MOLM-14 shSCR/FLT3
24 hours Dox
MV4-11 shSCR/FLT3
24 hours Dox
MOLM-14 Veh/QUIZ
14 hours
MV4-11 Veh/QUIZ
14 hours
Gene expression Profiling
FLT3-ITD_UP cell line signature
down after FLT3 inhibition
298
40
6
1
49
107
1
62
914
58
299
164
205
108
310
E
Hallmark 2020 gene set enriched in FLT3-ITD
- log10 (p value)
E2F Targets
Myc Targets V2
mTORC1 Signaling
G2-M Checkpoint
DNA Repair
Cholesterol Homeostasis
Oxidative Phosphorylation
Odds Ratio
KEGG 2021 gene set enriched in FLT3-ITD
DNA replication
Cell cycle
Arginine and proline metabolism
PPAR signaling pathway
Fatty acid biosynthesis
Biosynthesis of unsaturated fatty acids
Glutathione metabolism
Steroid biosynthesis
Terpenoid backbone biosynthesis
Cysteine and methionine metabolism
Odds Ratio
Lipid
Amino acid
Mitochondria
Redox
Signaling pathway
Cell cycle
DNA/RNA
F
KEGG 2021 gene set enriched in MLL-AF9 FLT3-ITDON (24h)
GSE163932
-log10 (pvalue)
Lipid
Amino acid
Redox
Steroid biosynthesis
Fatty acid biosynthesis
Terpenoid backbone biosynthesis
Biosynthesis of unsaturated fatty acids
Fatty acid elongation
Odds Ratio

**Figure S1**

**Supplementary Figure 1: Lipid metabolism dependency uncovered by transcriptomics in FLT3-mutant AML. (A)** Total protein extracts from MOLM-14 and MV4-11 cells incubated with vehicle or 3nM QUIZ for 14h were immunoblotted with indicated antibodies (n=3). **(B)** MOLM-14 and MV4-11 cells were transduced with Dox-inducible CTL or anti-FLT3 shRNAs. Total protein exctracts were immunoblotted with indicated antibodies. **(C)** Cell viability was measured by PI staining in MOLM-14 and MV4-11 cells after FLT3 inhibition using 3nM QUIZ for 14h (*left panel*) or shFLT3 induction (*right panel*). **(D)** Differential gene expression (DGE) was analyzed in MOLM-14 and MV4-11 cells after FLT3 inhibition using QUIZ or shFLT3, compared to vehicle or shCTL conditions, respectively. The FLT3-ITD_UP cell line signature was generated by 299 commonly downregulated genes (FC<0.67; FDR q-value <0.05) after FLT3 inhibition. **(E)** Enrichr analyses of Hallmark 2020 (*left panel*) and KEGG 2021 (*right panel*) gene signatures enriched in the FLT3-ITD_UP cell line signature (p-value<0.05) **(F)** Enrichr analyses of KEGG 2021 gene signatures enriched in upregulated genes in FLT3-ITD$^{ON}$ compared to FLT3-ITD$^{OFF}$ MLL-AF9 murine AML cells from the GSE163932 dataset (p-value<0.05).

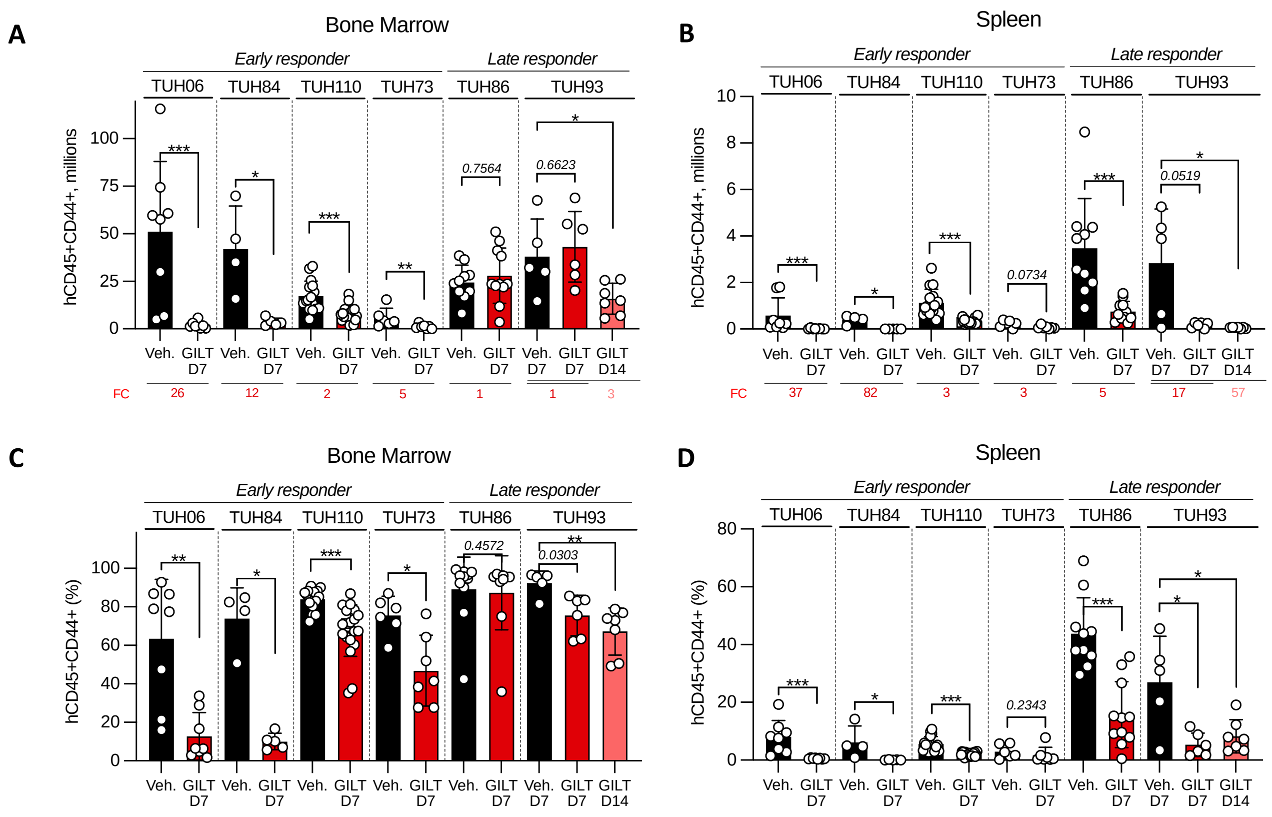

A
Bone Marrow
Early responder
Late responder
TUH06
TUH84
TUH110
TUH73
TUH86
TUH93
hCD45+CD44+, millions
Veh.
GILT D7
GILT D14
FC
26
12
2
5
1
1
3
0.7564
0.6623
B
Spleen
hCD45+CD44+, millions
FC
37
82
3
3
5
17
57
0.0734
0.0519
C
Bone Marrow
hCD45+CD44+ (%)
0.4572
0.0303
D
Spleen
hCD45+CD44+ (%)
0.2343


**Figure S2**

**Supplementary Figure 2: Anti-leukemic activity of GILT across a panel of AML PDX *ex vivo* and *in vivo*.**

**(A-B)** Number of viable CD33+/CD45+ human AML cells in bone marrow **(A)** and spleen **(B)** of six AML PDXs in vehicle and GILT groups. Fold change (FC) between the mean number of leukemic cells in vehicle and GILT groups are provided. **(C-D)** Percentage of viable CD33+/CD45+ human AML cells in bone marrow **(C)** and spleen **(D)** of six AML PDXs in vehicle and GILT groups. Vertical bars indicate standard deviations. *: $p<0.05$; **: $p<0.01$; ***: $p<0.001$.

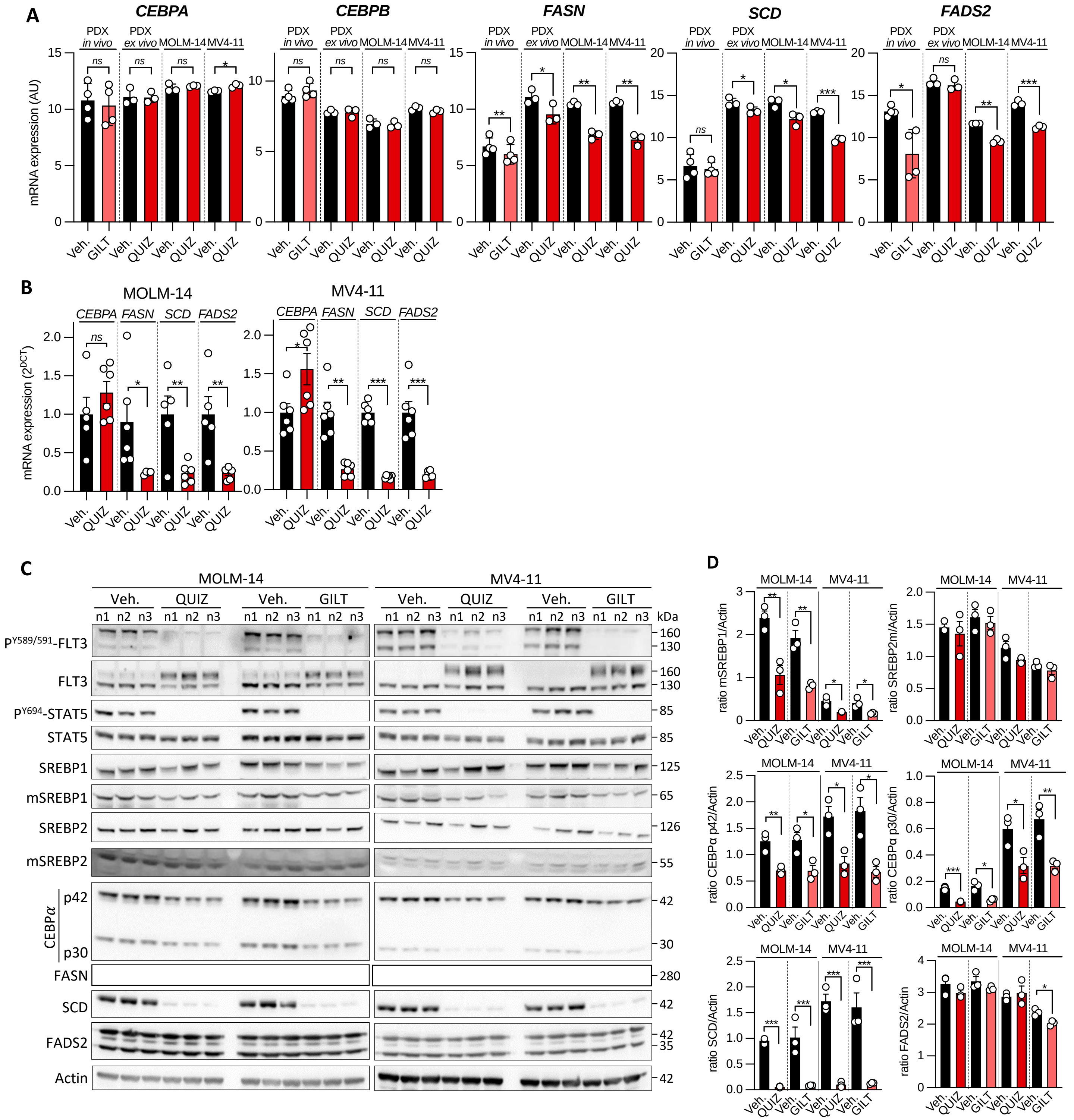
A
CEBPA
CEBPB
FASN
SCD
FADS2
PDX in vivo
PDX ex vivo
MOLM-14
MV4-11
mRNA expression (AU)
Veh.
GILT
QUIZ
B
MOLM-14
MV4-11
CEBPA
FASN
SCD
FADS2
mRNA expression (2^DCT)
C
MOLM-14
MV4-11
Veh.
QUIZ
GILT
n1
n2
n3
kDa
PY589/591-FLT3
FLT3
PY694-STAT5
STAT5
SREBP1
mSREBP1
SREBP2
mSREBP2
CEBPα
p42
p30
FASN
SCD
FADS2
Actin
D
ratio mSREBP1/Actin
ratio SREBP2m/Actin
ratio CEBPα p42/Actin
ratio CEBPα p30/Actin
ratio SCD/Actin
ratio FADS2/Actin

**Figure S3**

**Supplementary Figure 3: FLT3-ITD regulates the expression of C/EBPα and of proteins related to lipid biosynthesis in AML cell lines.**

**(A)** Gene expression of *CEBPA, CEBPB, FASN, SCD* and *FADS2* upon FLT3i in *in vivo*-treated PDX, *ex* vivo-treated PDX and AML cell lines (MOLM-14 and MV4-11) from transcriptomic analysis. **(B-D)** MOLM-14 and MV4-11 cells were incubated with 3nM QUIZ or 30nM GILT for 14h. **(B)** Gene expression of *CEBPA*, *FASN*, *SCD* and *FADS2* upon QUIZ was investigated by quantitative RT-PCR. **(C-D)** Total protein extracts were submitted to Western blots with indicated antibodies **(C)** and quantified **(D)** (n=3).

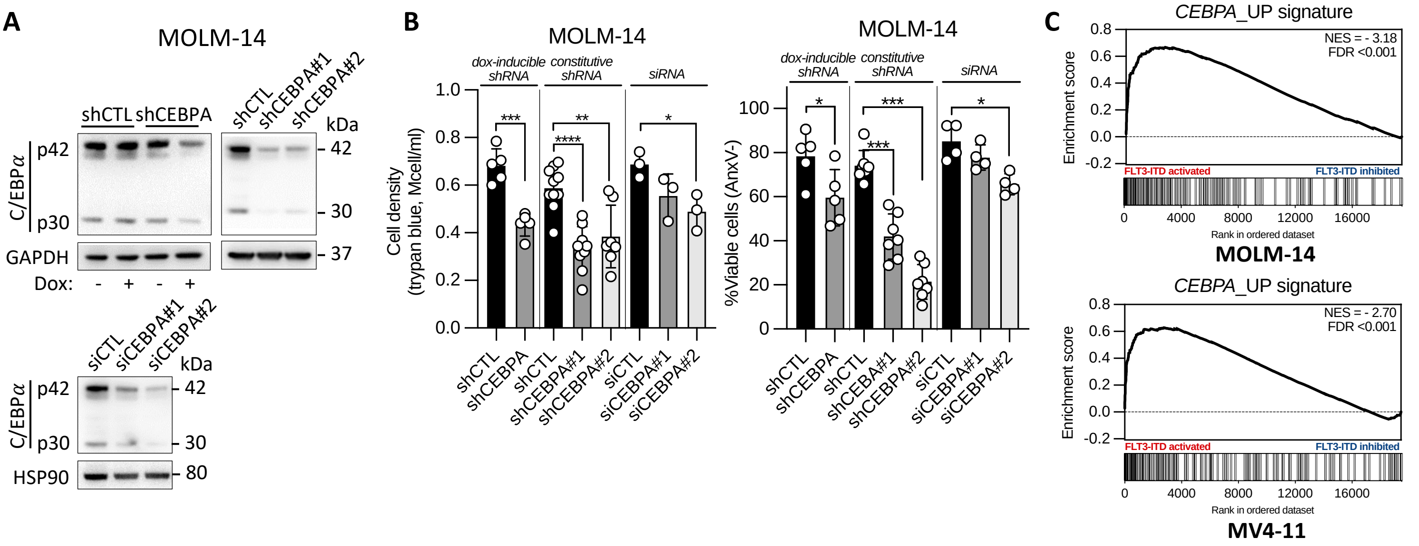

A
MOLM-14
shCTL
shCEBPA
shCTL
shCEBPA#1
shCEBPA#2
kDa
C/EBPα
p42
p30
GAPDH
Dox:
- + - +
42
30
37
siCTL
siCEBPA#1
siCEBPA#2
kDa
42
30
HSP90
80
B
MOLM-14
dox-inducible shRNA
constitutive shRNA
siRNA
Cell density (trypan blue, Mcell/ml)
%Viable cells (AnxV-)
shCTL
shCEBPA
shCEBPA#1
shCEBPA#2
siCTL
siCEBPA#1
siCEBPA#2
shCEBA#1
C
CEBPA_UP signature
NES = - 3.18
FDR <0.001
NES = - 2.70
Enrichment score
FLT3-ITD activated
FLT3-ITD inhibited
Rank in ordered dataset
MOLM-14
MV4-11


**Figure S4**

**Supplementary Figure 4: FLT3 and C/EBPα co-dependency in FLT3-ITD AML. (A-B)** MOLM-14 cells were transduced with Dox-inducible or constitutive (72h) shRNAs using lentivirus, or transfected with siRNA (24h) to deplete *CEBPA* and compared to their cognate control interfering RNA. **(A)** Total protein extracts were immunoblotted with indicated antibodies. **(B)** Cell viability assessed by trypan blue exclusion assay (*left panel*) and by annexin V binding assays (*right panel*). **(C)** GSEA of *CEBPA*_UP signature in MOLM-14 (*top panel*) and MV4-11 (*bottom panel*) cells incubated with vehicle (FLT3-ITD activated) or 3nM QUIZ (FLT3-ITD inhibited) for 14h.

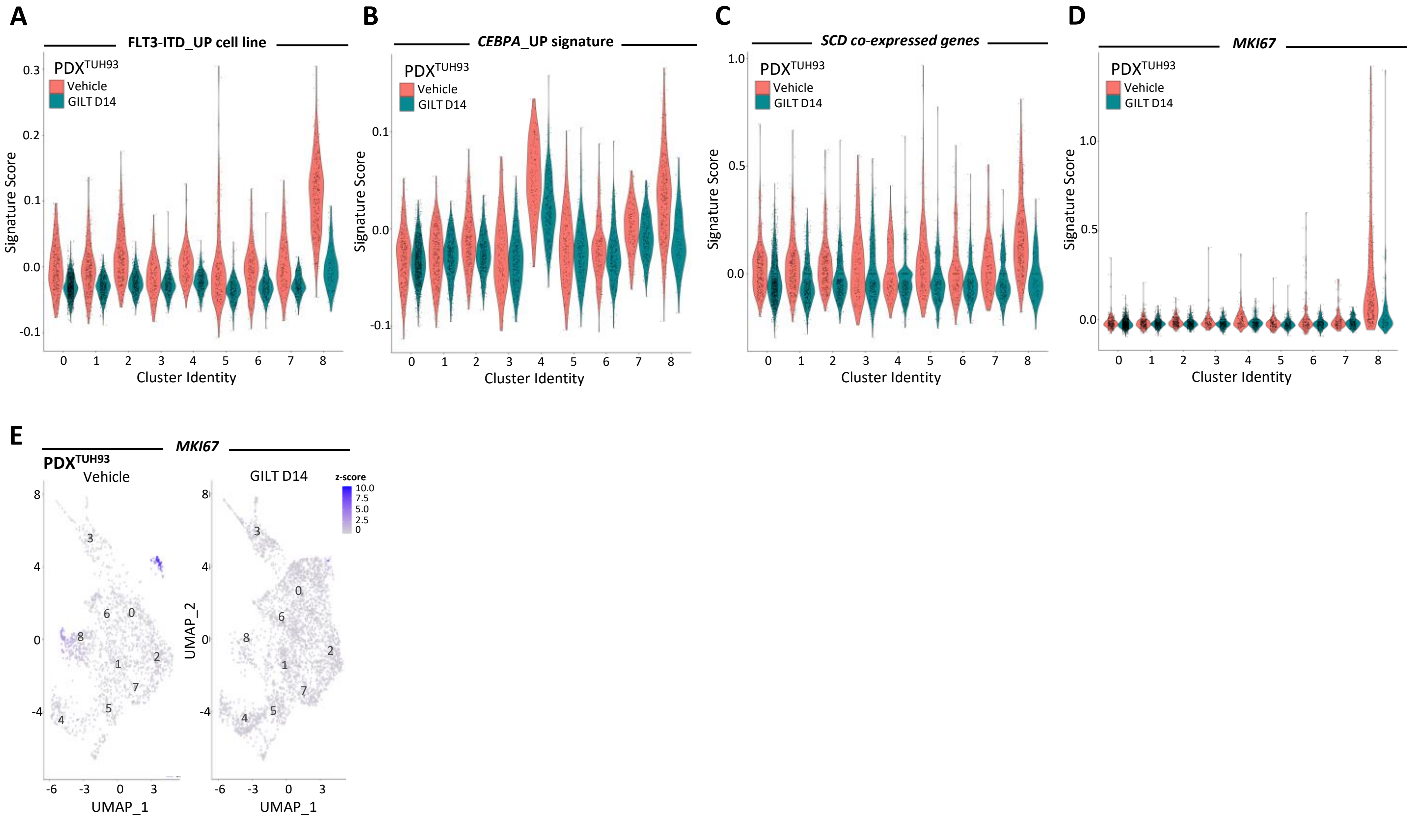
A
FLT3-ITD_UP cell line
B
CEBPA_UP signature
C
SCD co-expressed genes
D
MKI67
PDX^TUH93
Vehicle
GILT D14
Signature Score
Cluster Identity
E
MKI67
PDX^TUH93
Vehicle
GILT D14
z-score
UMAP_1
UMAP_2

Figure S5

**Supplementary Figure 5: Enrichment analysis in scRNA-seq from PDX$^{TUH93}$**. **(A-D)** Violin plot showing the enrichment of FLT3-ITD_UP cell line signature **(A)**, CEBPA_UP signature **(B)**, *SCD* co-expressed genes **(C)** and *MKI67* gene **(D)** by cluster in vehicle- compared to GILT-treated PDX$^{TUH93}$. **(E)** Visualization on the UMAP plot of *MKI67* gene enrichment in vehicle- compared to GILT-treated PDX$^{TUH93}$

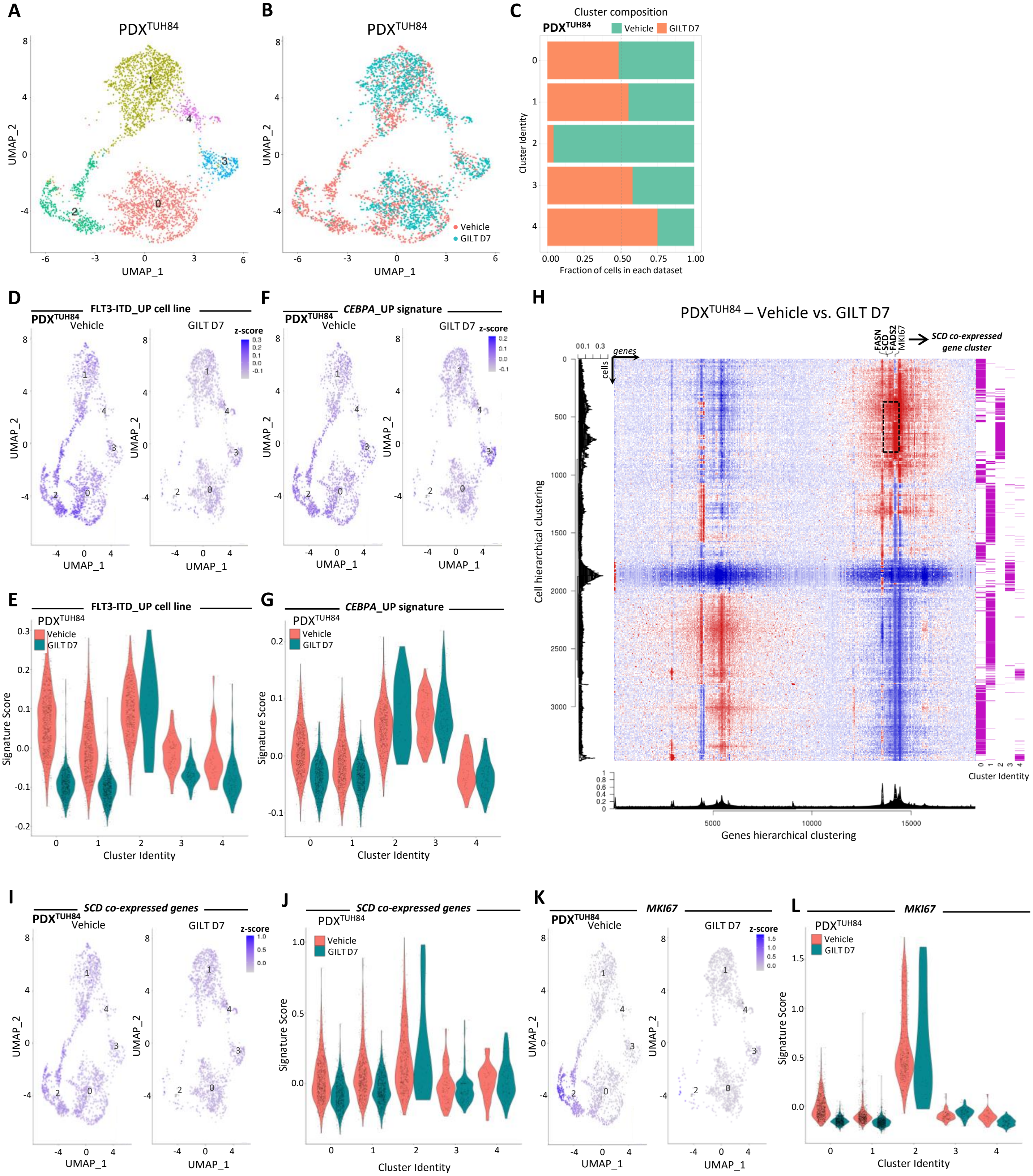

A
PDX TUH84
UMAP_1
UMAP_2
B
Vehicle
GILT D7
C
Cluster composition
Cluster Identity
Fraction of cells in each dataset
D
FLT3-ITD_UP cell line
z-score
E
Signature Score
Cluster Identity
F
CEBPA_UP signature
G
H
PDX TUH84 – Vehicle vs. GILT D7
FASN
SCD
FADS2
MKI67
SCD co-expressed gene cluster
genes
cells
Cell hierarchical clustering
Genes hierarchical clustering
I
SCD co-expressed genes
J
K
MKI67
L


Figure S6

**Supplementary Figure 6: scRNA-seq analysis from PDX$^{TUH84}$. (A)** Uniform manifold approximation and projection (UMAP) plot of 3623 cells from PDX$^{TUH84}$ using Seurat. Colors indicate k-means clusters (k=8). **(B)** UMAP plot colored by treatment condition (vehicle: n=1972 cells; GILT D7: n=1651 cells). **(C)** Proportion of each condition per cluster from PDX$^{TUH84}$ **(D)** Visualization on the UMAP plot of FLT3-ITD_UP cell line signature enrichment in vehicle- compared to GILT-treated PDX$^{TUH84}$. **(E)** Violin plot showing the enrichment of FLT3-ITD_UP cell line signature by cluster in vehicle- compared to GILT-treated PDX$^{TUH84}$. **(F)** Visualization on the UMAP plot of *CEBPA*_UP signature enrichment in vehicle- compared to GILT-treated PDX$^{TUH84}$. **(G)** Violin plot showing the enrichment of *CEBPA*_UP signature by cluster in vehicle- compared to GILT-treated PDX$^{TUH84}$. **(H)** Heatmap showing the expression of each gene per cell in vehicle- compared to GILT-treated PDX$^{TUH84}$. Dendograms reprensents the hierarchical clustering of cells (*left*) and genes (*bottom*). Cluster identities are shown on the right of the Heatmap. **(I)** Visualization on the UMAP plot of *SCD* co-expressed genes enrichment in vehicle- compared to GILT-treated PDX$^{TUH84}$. **(J)** Violin plot showing the enrichment of *SCD* co-expressed genes by cluster in vehicle- compared to GILT-treated PDX$^{TUH84}$. **(K)** Visualization on the UMAP plot of *MKI67* gene enrichment in vehicle- compared to GILT-treated PDX$^{TUH84}$. **(L)** Violin plot showing the enrichment of *MKI67* gene by cluster in vehicle- compared to GILT-treated PDX$^{TUH84}$.

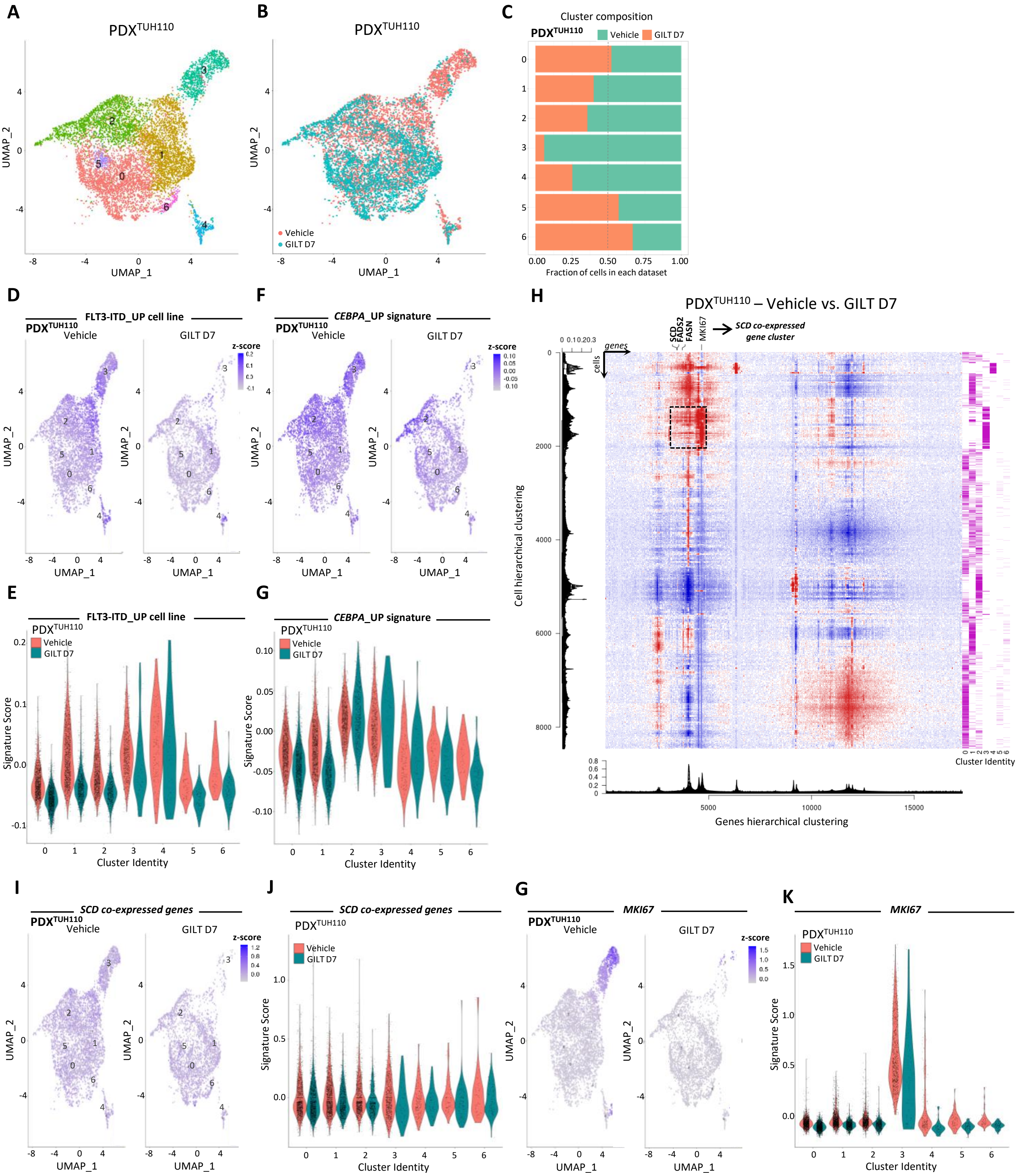

A
PDX^TUH110
UMAP_1
UMAP_2
B
Vehicle
GILT D7
C
Cluster composition
Fraction of cells in each dataset
D
FLT3-ITD_UP cell line
z-score
E
Signature Score
Cluster Identity
F
CEBPA_UP signature
G
H
PDX^TUH110 – Vehicle vs. GILT D7
SCD
FADS2
FASN
MKI67
SCD co-expressed gene cluster
genes
cells
Cell hierarchical clustering
Genes hierarchical clustering
I
SCD co-expressed genes
J
K
MKI67


Figure S7

**Supplementary Figure 7: scRNA-seq analysis from PDX$^{TUH110}$. (A)** Uniform manifold approximation and projection (UMAP) plot of 8739 cells from PDX$^{TUH110}$ using Seurat. Colors indicate k-means clusters (k=8). **(B)** UMAP plot colored by treatment condition (vehicle: n=5229 cells; GILT D7: n=3510 cells). **(C)** Proportion of each condition per cluster from PDX$^{TUH110}$. **(D)** Visualization on the UMAP plot of FLT3-ITD_UP cell line signature enrichment in vehicle- compared to GILT-treated PDX$^{TUH110}$. **(E)** Violin plot showing the enrichment of FLT3-ITD_UP cell line signature by cluster in vehicle- compared to GILT-treated PDX$^{TUH110}$. **(F)** Visualization on the UMAP plot of *CEBPA*_UP signature enrichment in vehicle- compared to GILT-treated PDX$^{TUH110}$. **(G)** Violin plot showing the enrichment of *CEBPA*_UP signature by cluster in vehicle- compared to GILT-treated PDX$^{TUH110}$. **(H)** Heatmap showing the expression of each gene per cell in vehicle- compared to GILT-treated PDX$^{TUH110}$. Dendograms reprensents the hierarchical clustering of cells (*left*) and genes (*bottom*). Cluster identities are shown on the right of the Heatmap. **(I)** Visualization on the UMAP plot of *SCD* co-expressed genes enrichment in vehicle- compared to GILT-treated PDX$^{TUH110}$. **(J)** Violin plot showing the enrichment of *SCD* co-expressed genes by cluster in vehicle- compared to GILT-treated PDX$^{TUH110}$. **(K)** Visualization on the UMAP plot of *MKI67* gene enrichment in vehicle- compared to GILT-treated PDX$^{TUH110}$. **(L)** Violin plot showing the enrichment of *MKI67* gene by cluster in vehicle- compared to GILT-treated PDX$^{TUH110}$.

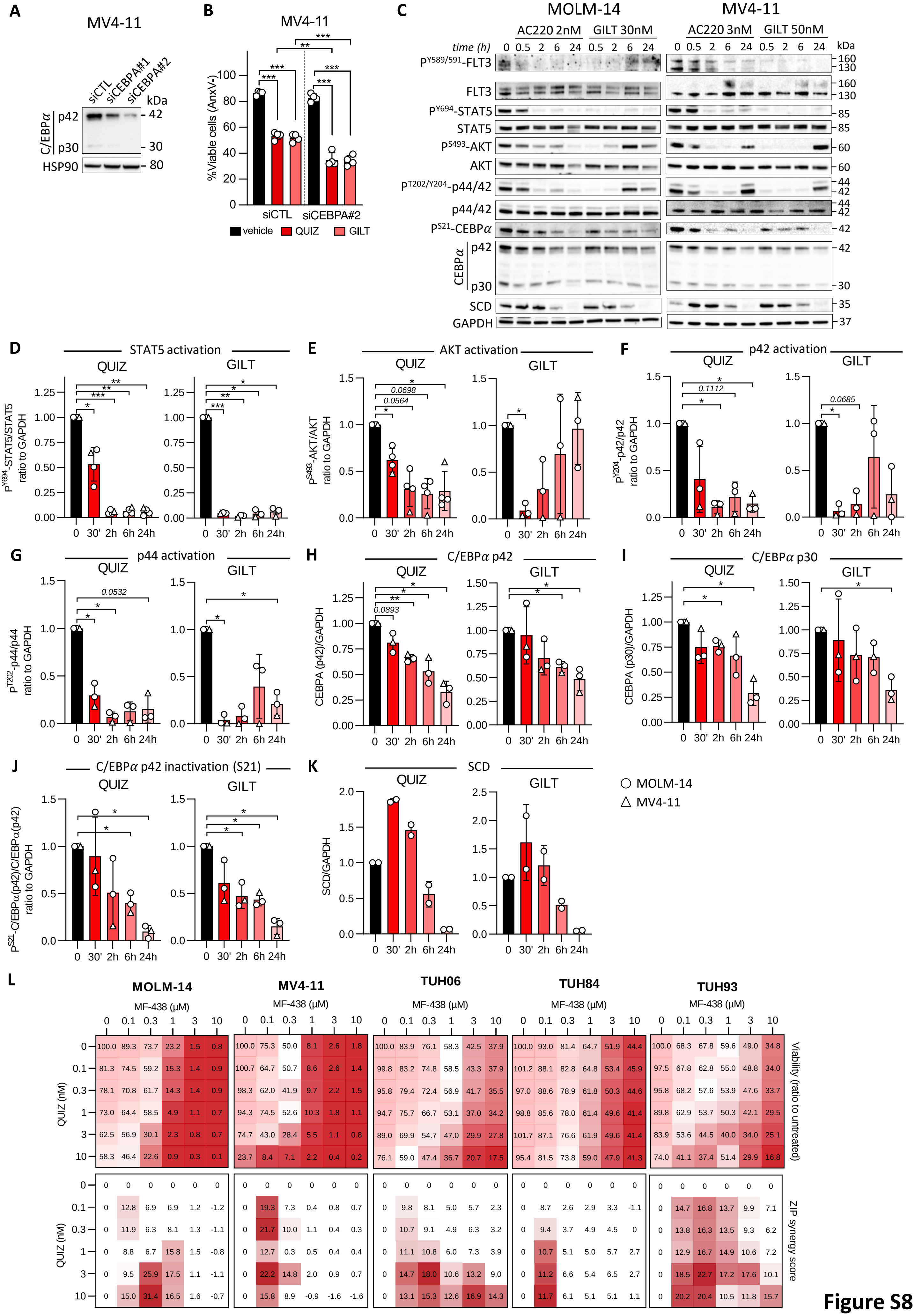
A
MV4-11
siCTL
siCEBPA#1
siCEBPA#2
kDa
C/EBPα
p42
p30
HSP90
B
MV4-11
%Viable cells (AnxV-)
siCTL
siCEBPA#2
vehicle
QUIZ
GILT
C
MOLM-14
MV4-11
AC220 2nM
GILT 30nM
AC220 3nM
GILT 50nM
time (h)
kDa
FLT3
STAT5
AKT
p44/42
CEBPα
SCD
GAPDH
D
STAT5 activation
E
AKT activation
F
p42 activation
G
p44 activation
H
C/EBPα p42
I
C/EBPα p30
J
C/EBPα p42 inactivation (S21)
K
SCD
QUIZ
GILT
SCD/GAPDH
MOLM-14
MV4-11
L
MOLM-14
MV4-11
TUH06
TUH84
TUH93
MF-438 (μM)
QUIZ (nM)
Viability (ratio to untreated)
ZIP synergy score

Figure S8

**Supplementary Figure 8: C/EBPα regulates rate-limiting lipid biosynthetic enzymes downstream of FLT3-ITD. (A-B)** MV4-11 cells were transfected with anti-*CEBPA* (siCEBPA#1 and siCEBPA#2) and control (siCTL) siRNAs. **(A)** Total protein extracts were analyzed 24h after transfection by Western blot with indicated antibodies. **(B)** Cell viability assessed by Annexin V staining after 14h incubation with 20nM QUIZ or 100nM GILT starting 6h after siRNAs transfection. **(C-K)** MOLM-14 and MV4-11 cells were incubated with 2-3nM QUIZ or 30-50nM GILT, respectively, for 30'to 24h and immunoblotted with indicated antibodies **(C)** and quantified **(D-K)**. **(L)** Patient-derived AML samples (n=3) and FLT3-ITD AML cell lines (MOLM-14 and MV4-11) were incubated with crossed dose-range QUIZ and MK-436 for 24h. Cell viability was measured using a luminescence-based assay (Alamar Blue) and results are expressed as a ratio to the untreated condition for each well in a viability matrix (*top panel*) and using synergy maps using zero interaction potency (ZIP) synergy scores and a threshold ZIP>10 considered as significant (*bottom panel*). Vertical bars indicate standard deviations. ns: not significant; *: p<0.05; **: p<0.01; ***: p<0.001.

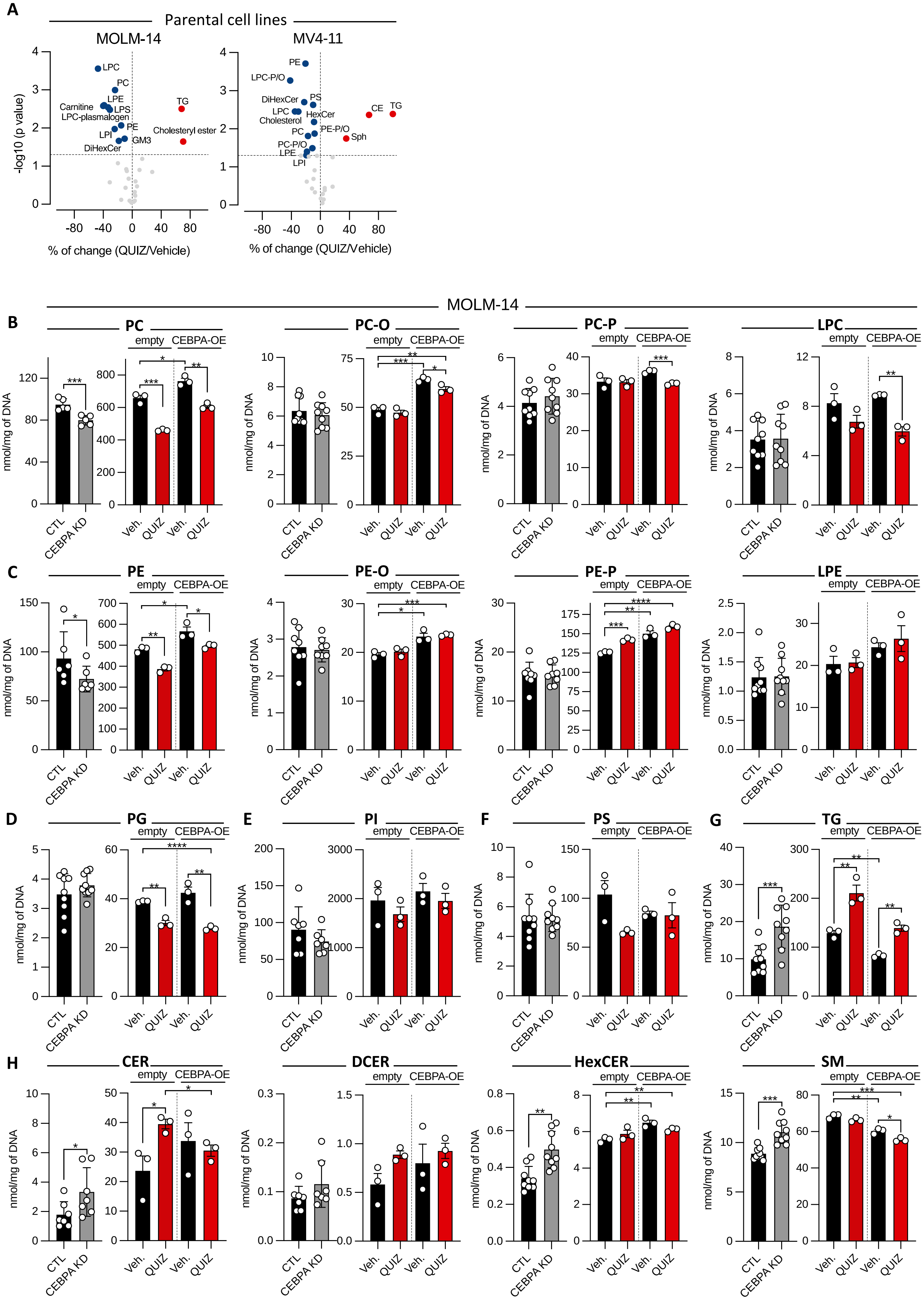
A
Parental cell lines
MOLM-14
MV4-11
-log10 (p value)
% of change (QUIZ/Vehicle)
LPC
PC
LPE
Carnitine
LPS
LPC-plasmalogen
PE
LPI
GM3
DiHexCer
TG
Cholesteryl ester
PE
LPC-P/O
DiHexCer
PS
LPC
HexCer
Cholesterol
PC
PE-P/O
PC-P/O
LPE
LPI
CE
TG
Sph
MOLM-14
B
PC
PC-O
PC-P
LPC
C
PE
PE-O
PE-P
LPE
D
PG
E
PI
F
PS
G
TG
H
CER
DCER
HexCER
SM
empty
CEBPA-OE
nmol/mg of DNA
CTL
CEBPA KD
Veh.
QUIZ

Figure S9

**Supplementary Figure 9: C/EBPα controls lipid amount in FLT3-ITD cell lines. (A)** Global lipidomics in MOLM-14 (*left panel*) and MV4-11 (*right panel*) parental AML cell lines incubated with vehicle or 3nM QUIZ for 14h. LPC/PC: lyso-/phosphatidylcholine; LPE/PE: lyso-/phosphatidylethanolamine; LPI/PI: lyso-/phosphatidylinositol; LPS/PS: lyso-/phosphatidylserine; TG: triglceride; CE/Cer: ceramide; DiHexCer: dihexosylceramide; HexCER: Hexosylceramide; SM/Sp: sphingomyelin; GM3: monosiolodihexosylganglioside; -P: plasmalogen; -O: ether. **(B-H)** Targeted lipidomics was performed in MOLM-14 cells were depleted from C/EBPα (siRNA 24h, dox-inducible shRNA or constitutive shRNA 72h) or overexpressing C/EBPα (dox-inducible 48h) and treated with vehicle or 3nM QUIZ for 24h. Each graph represents the quantity of each lipid species relative to DNA amount per sample. Vertical bars indicate standard deviations. *: $p<0.05$; **: $p<0.01$; ***: $p<0.001$

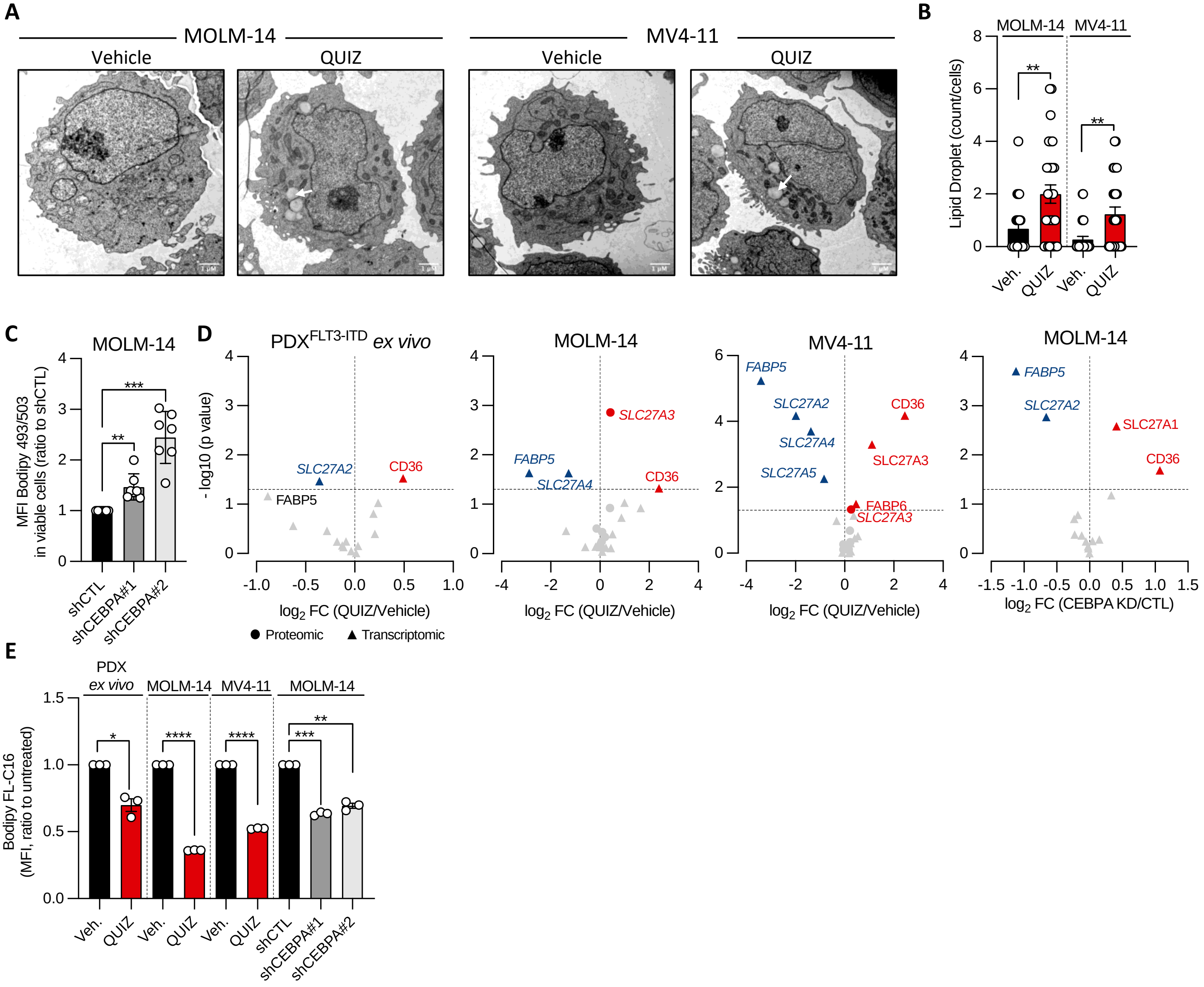
A
MOLM-14
MV4-11
Vehicle
QUIZ
Vehicle
QUIZ
B
MOLM-14
MV4-11
Lipid Droplet (count/cells)
**
**
Veh.
QUIZ
Veh.
QUIZ
C
MOLM-14
MFI Bodipy 493/503 in viable cells (ratio to shCTL)
***
**
shCTL
shCEBPA#1
shCEBPA#2
D
PDX^FLT3-ITD ex vivo
MOLM-14
MV4-11
MOLM-14
- log10 (p value)
SLC27A2
CD36
FABP5
SLC27A3
FABP5
SLC27A4
CD36
FABP5
SLC27A2
CD36
SLC27A4
SLC27A3
SLC27A5
FABP6
SLC27A3
FABP5
SLC27A2
SLC27A1
CD36
log2 FC (QUIZ/Vehicle)
log2 FC (QUIZ/Vehicle)
log2 FC (QUIZ/Vehicle)
log2 FC (CEBPA KD/CTL)
Proteomic
Transcriptomic
E
PDX ex vivo
MOLM-14
MV4-11
MOLM-14
Bodipy FL-C16 (MFI, ratio to untreated)
*
****
****
***
**
Veh.
QUIZ
Veh.
QUIZ
Veh.
QUIZ
shCTL
shCEBPA#1
shCEBPA#2

Figure S10

**Supplementary Figure 10: FLT3 inhibitors induce a lipid switch increasing neutral lipids dependent on C/EBPα. (A-B)** MOLM-14 and MV4-11 cells were incubated with vehicle or 3nM QUIZ for 14h. **(A)** Electron microscopy (EM) morphology (scale: 5μM). **(B)** EM quantification of lipid droplet content per cell. Each dot corresponds to a single cell. **(C)** Neutral lipid content in MOLM14 silenced for C/EBPα compared to control using constitutive shRNA measured by Bodiby 493/503 staining. **(D)** Gene and protein expression of FA binding protein and lipid transporter in FLT3-ITD patient-derived AML cells and cell lines incubated with vehicle or QUIZ, and in MOLM-14 cells transduced with CTL or anti-CEBPA shRNAs (decrease and increase FC<0.67 and >1.5, respectively). **(E)** Patient-derived FLT3-ITD AML cells and cell lines were incubated with 3nM QUIZ for 14h, and MOLM-14 cells were transduced with CTL or two different anti-CEBPA shRNAs. Lipid transport was assessed using Bodipy FL-C16 staining. Vertical bars indicate standard deviations. *: p<0.05; **: p<0.01; ***: p<0.001.

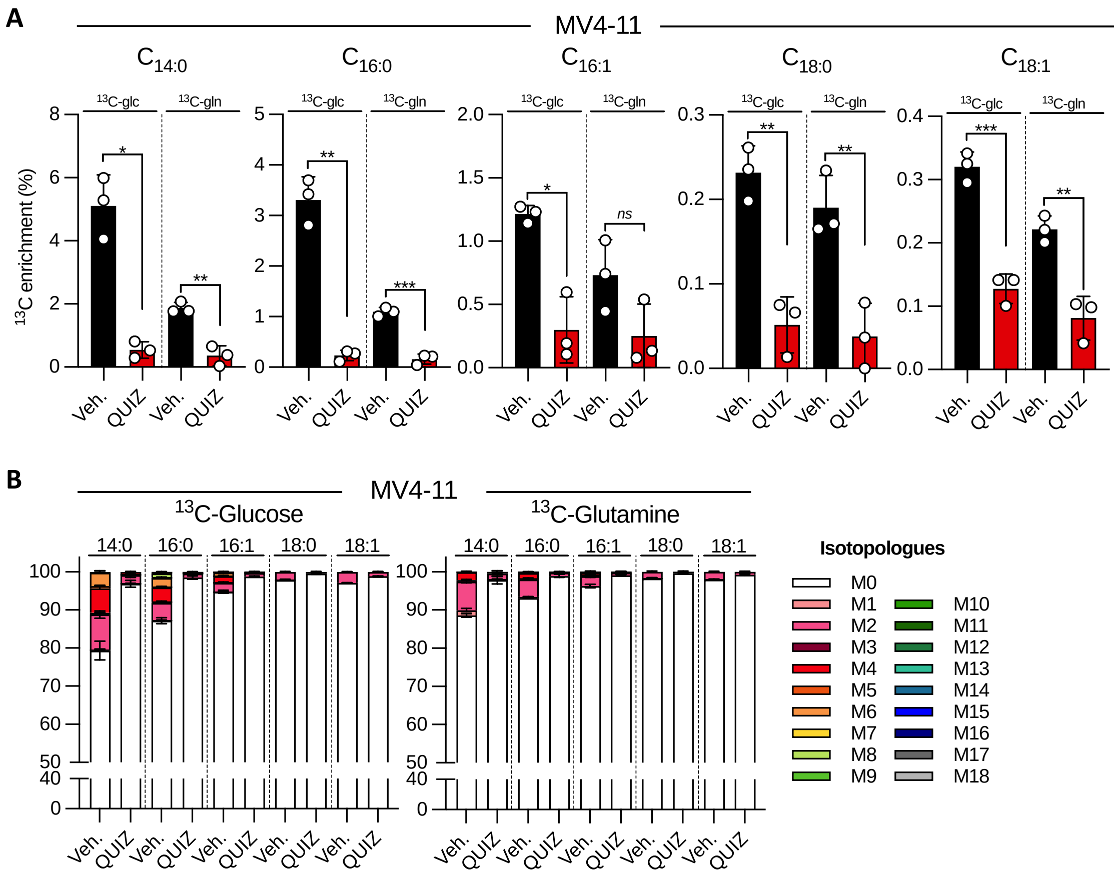
A
MV4-11
C14:0
C16:0
C16:1
C18:0
C18:1
13C-glc
13C-gln
13C enrichment (%)
Veh.
QUIZ
ns
B
MV4-11
13C-Glucose
13C-Glutamine
14:0
16:0
16:1
18:0
18:1
Isotopologues
M0
M1
M2
M3
M4
M5
M6
M7
M8
M9
M10
M11
M12
M13
M14
M15
M16
M17
M18

**Figure S11**

**Supplementary Figure 11: FLT3 inhibitor inhibits fatty acid synthesis fueled by glucose and glutamine. (A)** Percentage of $^{13}C$ enrichment in $C_{14:0}$, $C_{16:0}$, $C_{16:1}$, $C_{18:0}$ and $C_{18:1}$ total FA in MV4-11 cultured on [U-$^{13}C$]-glucose or [U-$^{13}C$]-glutamine and treated with vehicle or 3nM QUIZ for 24h. **(B)** Isotopologues distribution in $C_{14:0}$, $C_{16:0}$, $C_{16:1}$, $C_{18:0}$ and $C_{18:1}$ total FA in MV4-11 24-cultured on [U-$^{13}C$]-glucose (*left* panel) or [U-$^{13}C$]-glutamine (*right* panel) and incubated with vehicle or 3nM QUIZ. Vertical bars indicate standard deviations. *: $p<0.05$; **: $p<0.01$; ***: $p<0.001$.

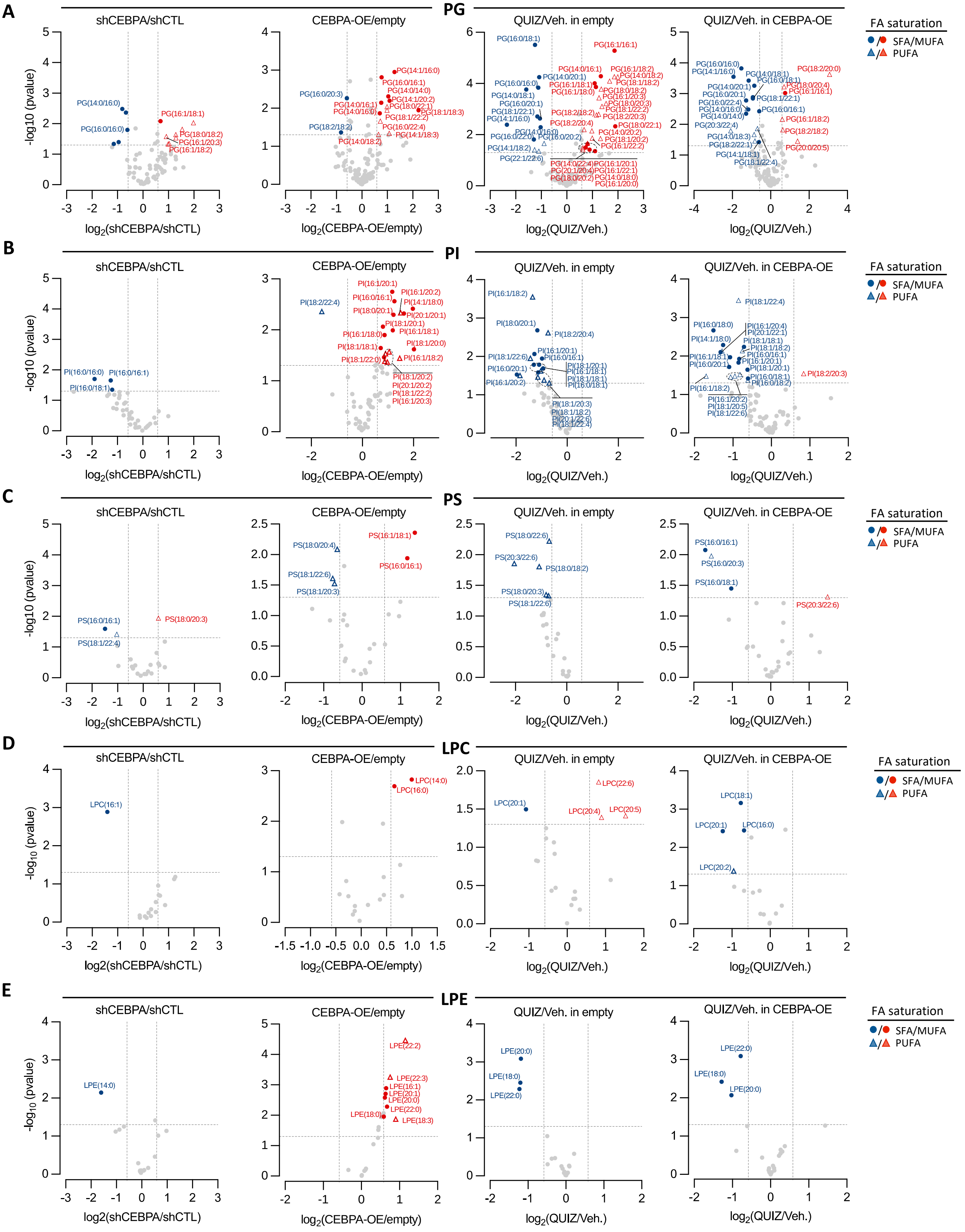

A
PG
shCEBPA/shCTL
CEBPA-OE/empty
QUIZ/Veh. in empty
QUIZ/Veh. in CEBPA-OE
-log10 (pvalue)
log2(shCEBPA/shCTL)
log2(CEBPA-OE/empty)
log2(QUIZ/Veh.)
FA saturation
SFA/MUFA
PUFA
B
PI
C
PS
D
LPC
E
LPE


**Figure S12**

**Supplementary Figure 12: FLT3 inhibition decreases monounsaturated fatty acid dependent on C/EBPα.** Targeted lipidomics showing repartition of SFAs, MUFAs and PUFAs distribution into phosphatidylglycerol (PG) **(A)**, phosphatidylinositol (PI) **(B)**, phosphatidylserine (PS), lysophosphatidylcholine (LPC) and lysophosphatidylethanolamine (LPE) in C/EBPα-depleted MOLM-14 compared to control, CEBPA-OE compared to empty MOLM-14, empty MOLM-14 incubated with vehicle or 3nM QUIZ for 14h, and CEBPA-OE MOLM-14 incubated with vehicle or 3nM QUIZ for 14h (from the left to the right, respectively).

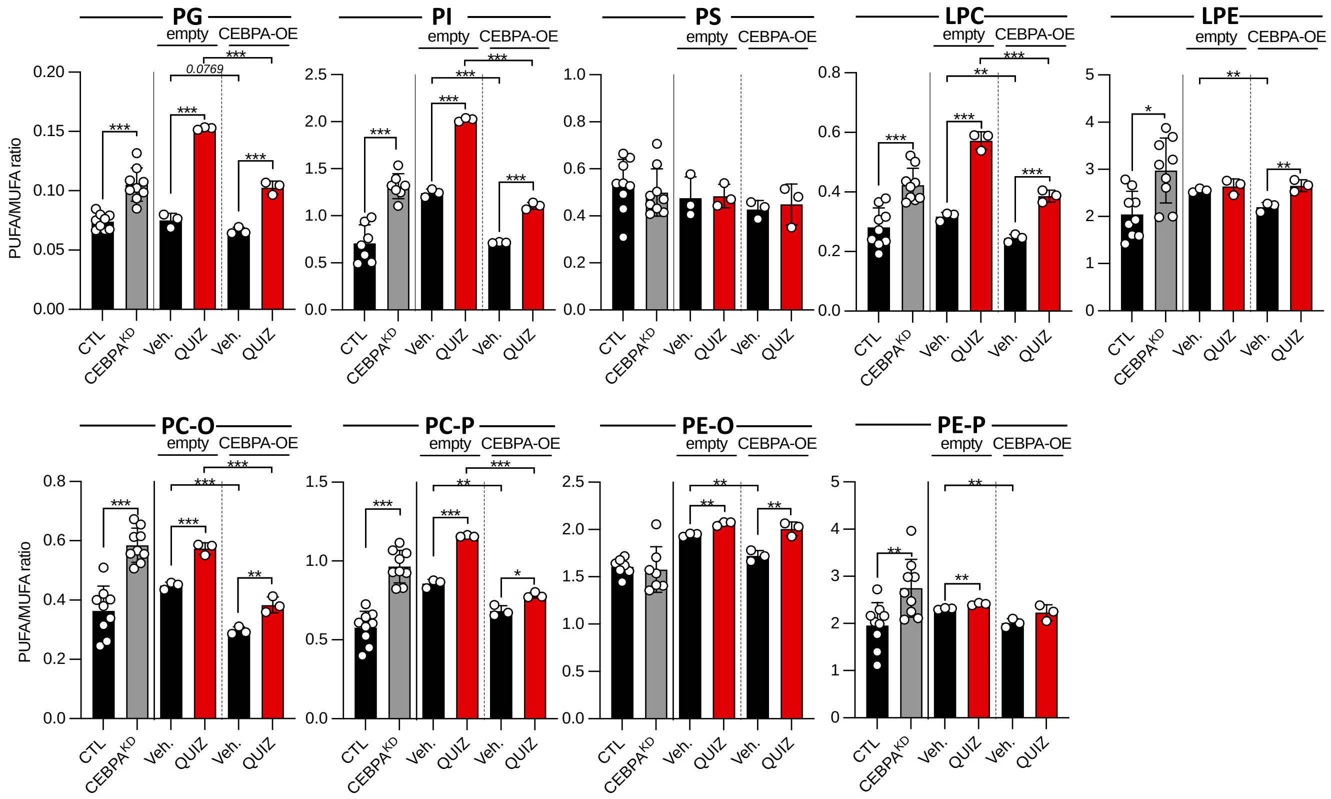
PG
PI
PS
LPC
LPE
PC-O
PC-P
PE-O
PE-P
empty
CEBPA-OE
PUFA/MUFA ratio
0.0769
CTL
CEBPA$^{KD}$
Veh.
QUIZ

**Figure S13**

**Supplementary Figure 13: FLT3 inhibition increases PUFA/MUFA ratio dependent on C/EBPα.** PUFA/MUFA ratio into phosphatidylglycerol (PG), phosphatidylinositol (PI), phosphatidylserine (PS), lysophosphatidylcholine (LPC), lysophosphatidylethanolamine (LPE), phosphatidylcholine-ether/-plasmogen (PC-O/PC-P) and phosphatidylethanolamine-ether/-plasmogen (PE-O/PE-P) in C/EBPα-depleted MOLM-14 compared to control, CEBPA-OE compared to empty MOLM-14, empty MOLM-14 incubated with vehicle or 3nM QUIZ for 14h, and CEBPA-OE MOLM-14 incubated with vehicle or 3nM QUIZ for 14h (from the left to the right, respectively). Vertical bars indicate standard deviations. *: $p<0.05$; **: $p<0.01$; ***: $p<0.001$.

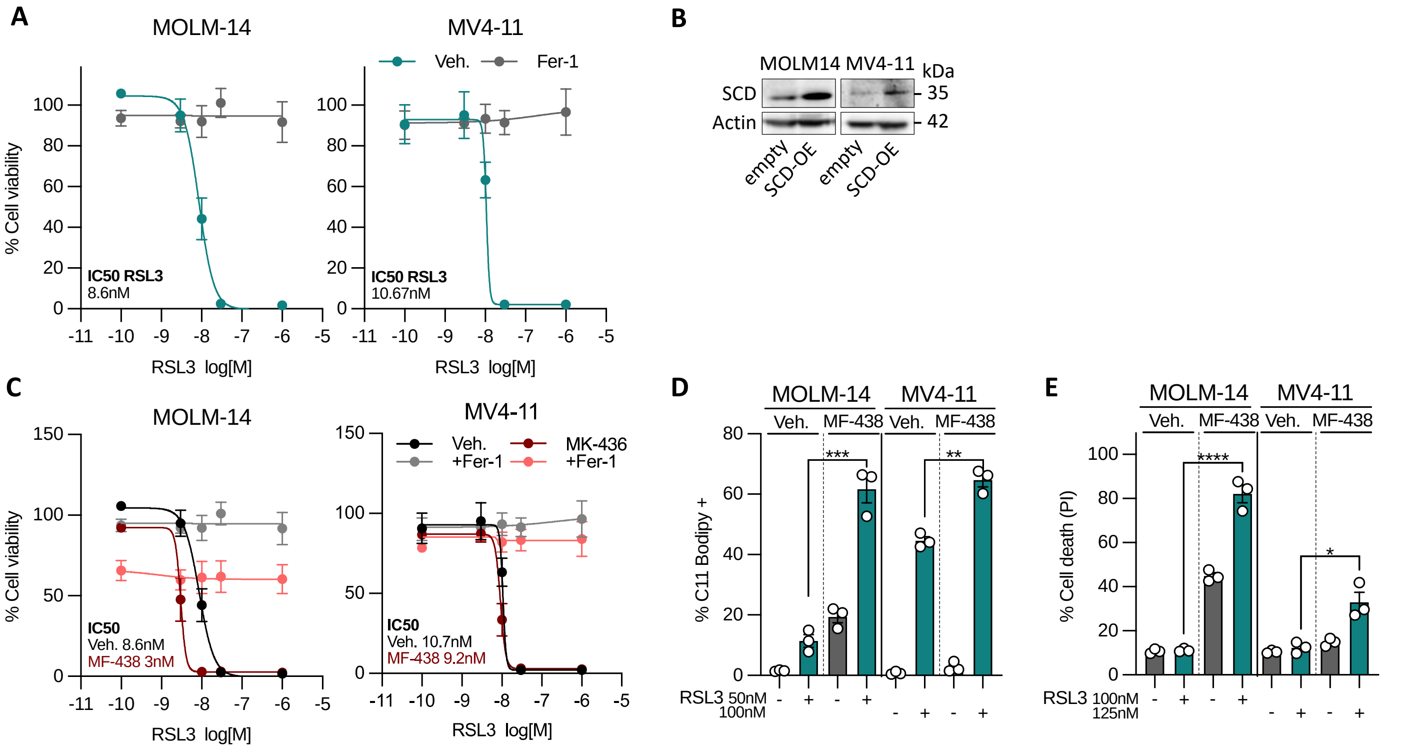
A
MOLM-14
MV4-11
Veh.
Fer-1
% Cell viability
IC50 RSL3
8.6nM
IC50 RSL3
10.67nM
RSL3 log[M]
B
MOLM14 MV4-11
kDa
SCD
Actin
35
42
empty
SCD-OE
C
MOLM-14
MV4-11
Veh.
+Fer-1
MK-436
+Fer-1
IC50
Veh. 8.6nM
MF-438 3nM
Veh. 10.7nM
MF-438 9.2nM
D
MF-438
% C11 Bodipy +
RSL3 50nM
100nM
E
% Cell death (PI)
RSL3 100nM
125nM

Figure S14

**Supplementary Figure 14: Ferroptotic cell death induced by FLT3i is mediated by inhibition of SCD-dependent mono-unsaturated FA synthesis. (A)** MOLM-14 (*left panel*) and MV4-11 (*right panel*) cells were exposed to dose-range RLS3 with or without 10µM Fer-1 for 48h. Cell viability assays using Alamar Blue. **(B)** MOLM-14 and MV4-11 cells were transduced with a vector allowing a constitutive expression of SCD, or with the empty vector. Total protein exctracts were immunoblotted with indicated antibodies. **(C)** MOLM-14 (*left panel*) and MV4-11 (*right panel*) cells were incubated with vehicle or 3µM MF-438, and exposed to dose-range RLS3 with or without 10µM Fer-1 for 48h. Cell viability assays using Alamar Blue. **(D-E)** MOLM-14 and MV4-11 cells were incubated with vehicle or 1µM MF-438, combined with vehicle or 50-125nM RSL3. Lipid peroxidation was measured by C11 Bodipy after 12h-treatment **(D)** and cell viability was assessed by PI staining after 24h-treatment **(E).** Vertical bars indicate standard deviations. *: $p<0.05$; **: $p<0.01$; ***: $p<0.001$.

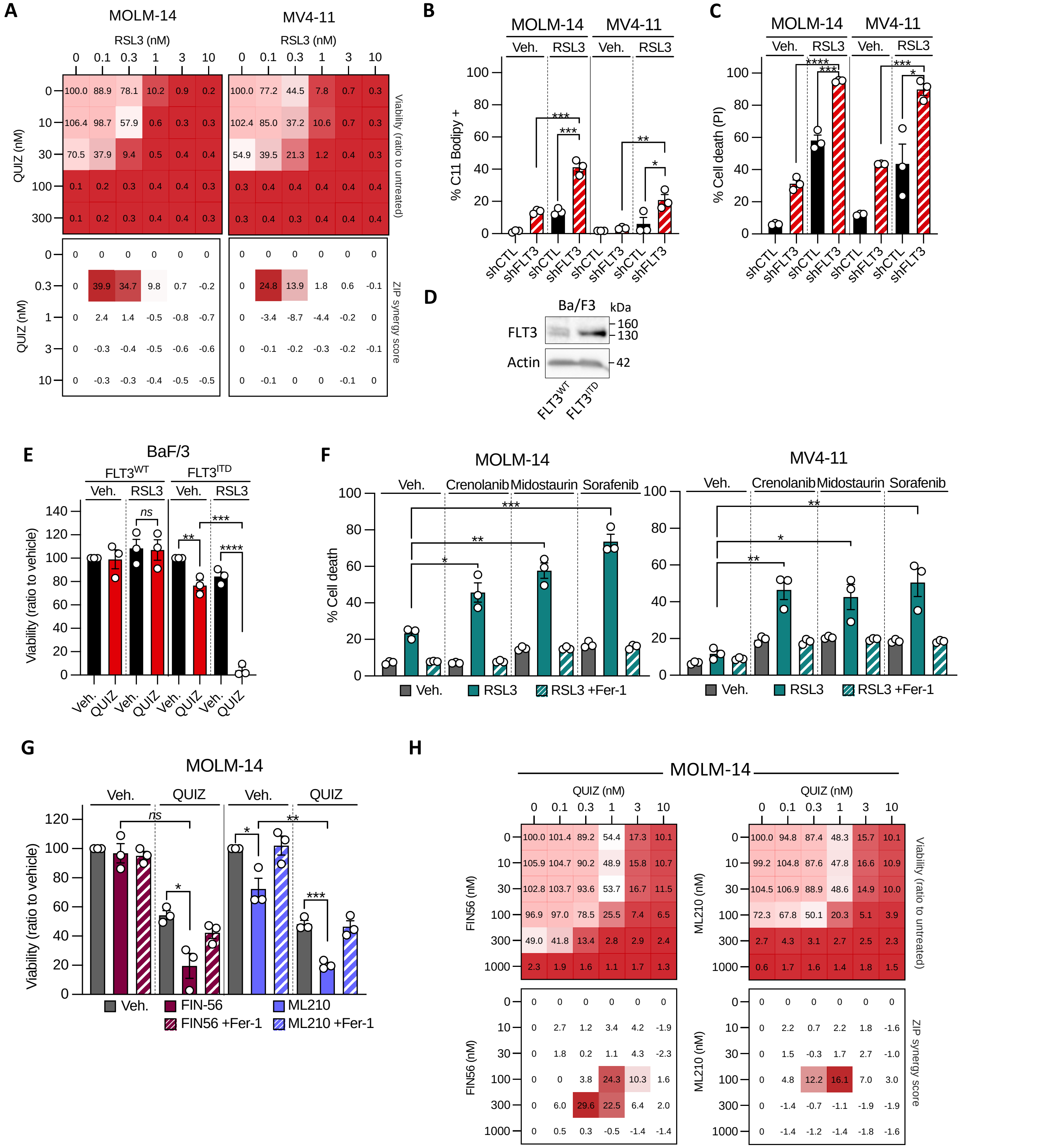
A
MOLM-14
MV4-11
RSL3 (nM)
QUIZ (nM)
Viability (ratio to untreated)
ZIP synergy score
B
% C11 Bodipy +
Veh.
RSL3
shCTL
shFLT3
C
% Cell death (PI)
D
Ba/F3
kDa
FLT3
Actin
160
130
42
FLT3WT
FLT3ITD
E
BaF/3
FLT3WT
FLT3ITD
Viability (ratio to vehicle)
QUIZ
ns
F
% Cell death
Crenolanib
Midostaurin
Sorafenib
RSL3 +Fer-1
G
FIN-56
FIN56 +Fer-1
ML210
ML210 +Fer-1
H
FIN56 (nM)
ML210 (nM)

Figure S15

**Supplementary Figure 15: FLT3 inhibitors unmask a vulnerability of FLT3-mutant leukemic cells to ferroptosis. (A)** FLT3-ITD AML cell lines (MOLM-14 and MV4-11) were incubated with crossed dose-range QUIZ and RSL3 for 48h. Cell viability was measured using a luminescence-based assay (Alamar Blue) and results are expressed as a ratio to the untreated condition for each wheel in a viability matrix (top panel) and using synergy maps using zero interaction potency (ZIP) synergy scores and a threshold ZIP>10 considered as significant (bottom panel). **(B-C)** MOLM-14 and MV4-11 cells were transduced with CTL or FLT3-targeting shRNAs (dox-inducible, 24h), and incubated with vehicle or 100nM **(B)** or 50nM **(C)** RSL3. Lipid peroxidation was measured by C11 Bodipy staining after 14h-treatment **(B)**, and cell viability was assessed by PI staining after 24h-treatment **(C). (D-E)** IL3-dependent Ba/F3 murine hematopoietic cells were transduced with expression vectors for *FLT3* ($FLT3^{WT}$) or *FLT3*-ITD-mutated ($FLT3^{ITD}$) genes. $FLT3^{WT}$ and $FLT3^{ITD}$ Ba/F3 cells were grown with and without IL3, respectively. **(D)** Total protein extracts were immunoblotted with indicated antibodies. **(E)** Cells were cultured 48h with vehicle or 3nM QUIZ, and vehicle or 1μM RSL3, and cell viability was assessed using a luminescence-based assay (Alamar Blue). Results are expressed as a ratio to the vehicle conditions. **(F)** MOLM-14 (*left panel*) and MV4-11(*right panel*) cells were incubated with vehicle, or 10nM and 20nM Crenolanib respectively, or 60nM and 70nM Midostaurin respectively, or 30nM Sorafenib, with vehicle or 100nM RSL3 and with or without 10μM Fer-1 for 24h as indicated. **(G)** MOLM-14 cells were incubated with vehicle or 1nM QUIZ, combined without or with 100nM FIN-56 or 100nM ML-210 and with or without 10μM Fer-1 for 48h as indicated, and cell viability was assessed using a luminescence-based assay (Alamar Blue). **(H)** MOLM-14 cells were incubated with crossed dose-range QUIZ and FIN56 for 48h. Cell viability was measured using a luminescence-based assay (Alamar Blue) and results are expressed as a ratio to the untreated condition for each wheel in a viability matrix (top panel) and using synergy maps using zero interaction potency (ZIP) synergy scores and a threshold ZIP>10 considered as significant (bottom panel).

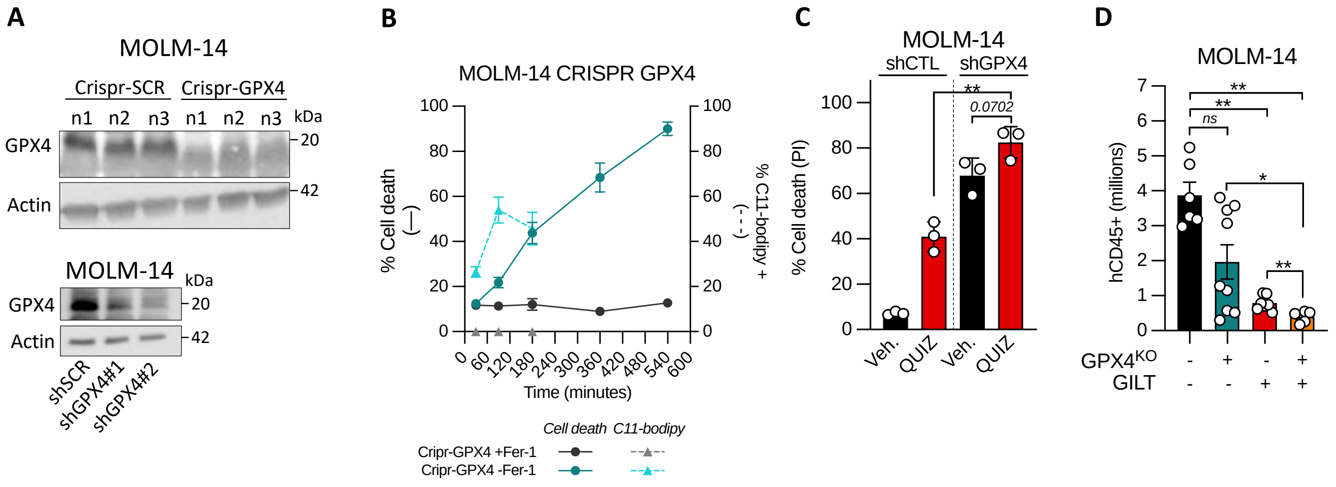
A
MOLM-14
Crispr-SCR
Crispr-GPX4
n1 n2 n3 n1 n2 n3
kDa
GPX4
Actin
20
42
MOLM-14
shSCR
shGPX4#1
shGPX4#2
B
MOLM-14 CRISPR GPX4
% Cell death (—)
% C11-bodipy + (- - -)
Time (minutes)
Cell death
C11-bodipy
Cripr-GPX4 +Fer-1
Cripr-GPX4 -Fer-1
C
MOLM-14
shCTL
shGPX4
% Cell death (PI)
0.0702
Veh.
QUIZ
Veh.
QUIZ
D
MOLM-14
hCD45+ (millions)
ns
GPX4KO
GILT

Figure S16

**Supplementary Figure 16: lipid redox stress induction by GPX4 inhibition primed FLT3i activity in FLT3-ITD AML cells. (A)** Total lysates of MOLM14 CRISPR-GPX4 versus CRISPR-CTL and shGPX4 versus shCTL were immunoblotted with indicated antibodies. **(B)** MOLM-14 cells were invalidated for GPX4 using CRISPR-Cas9, and cultured with 10μM Fer-1. Cell viability (using PI staining) and lipid peroxidation (using C11 Bodipy staining) were measured without or with Fer-1 dependent on time. **(C)** MOLM-14 cells were transduced with CTL or GPX4-targeting shRNAs (dox-inducible, 72h), and incubated with vehicle or 3nM QUIZ for 24h. Cell viability was measured by PI staining. **(D)** Number of viable CD45+ human AML cells in bone marrow of NSG mice xenografted with MOLM-14 cells invalidated or not for GPX4 using CRISPR-Cas9 and then treated with vehicle or GILT for 7 days.

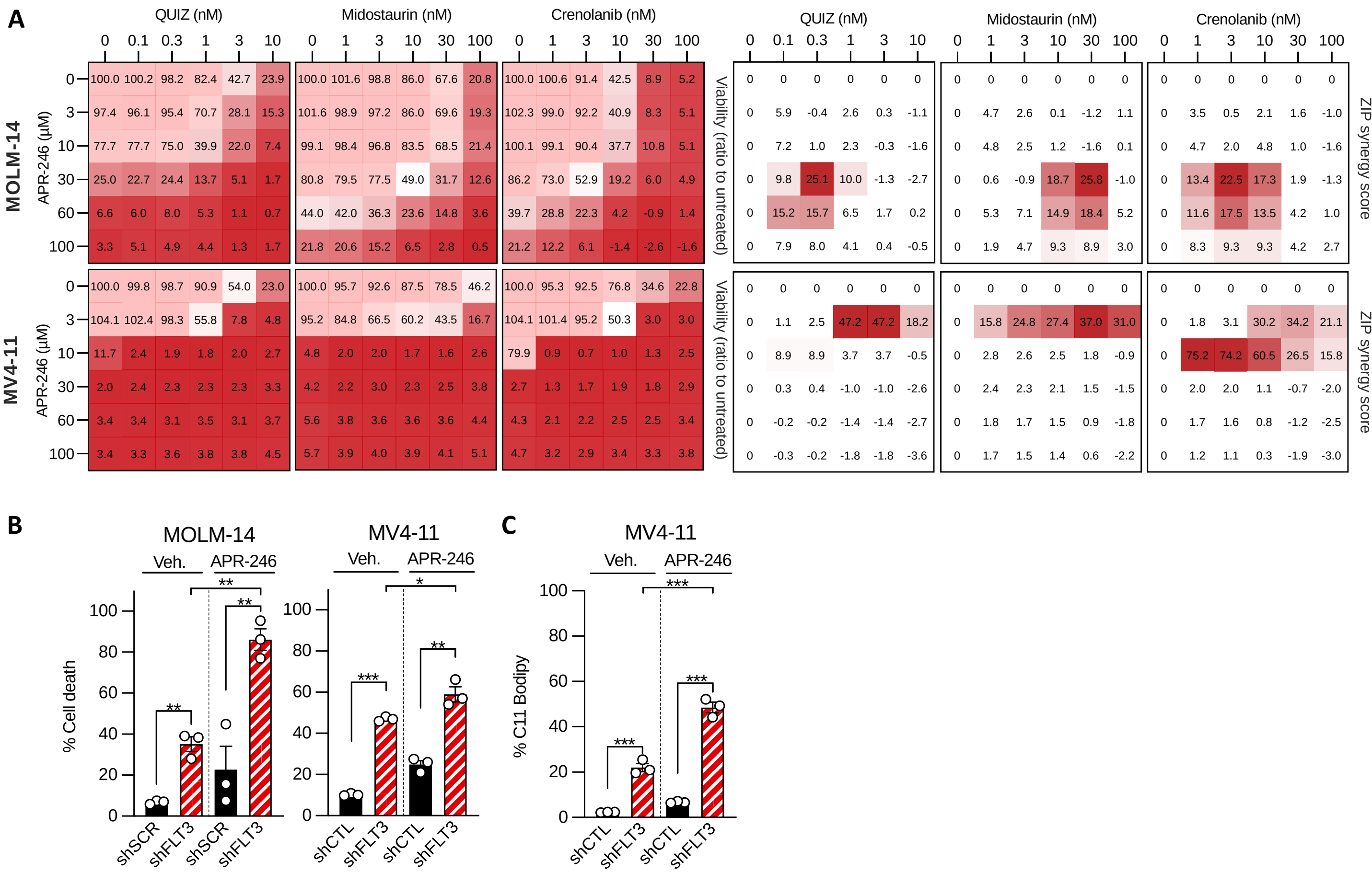
A
B
C
MOLM-14
MV4-11
QUIZ (nM)
Midostaurin (nM)
Crenolanib (nM)
APR-246 (µM)
Viability (ratio to untreated)
ZIP synergy score
% Cell death
% C11 Bodipy
Veh.
APR-246
shSCR
shFLT3
shCTL

**Figure S17**

**Supplementary Figure 17: Synergy between FLT3 inhibition and APR-246 in vitro. (A)** FLT3-ITD AML cell lines (MOLM-14 and MV4-11) were incubated with crossed dose-range QUIZ, Midostaurin or Crenolanib and APR-246 for 48h. Cell viability was measured using a luminescence-based assay (Alamar Blue) and results are expressed as a ratio to the untreated condition for each wheel in a viability matrix (*left panels*) and using synergy maps using zero interaction potency (ZIP) synergy scores and a threshold ZIP>10 considered as significant (*right panels*). **(B-C)** MOLM-14 and MV4-11 cells transduced with CTL or FLT3-targeting shRNAs were cultured with vehicle or 30-5µM APR-246, respectively, for 24h. **(B)** Cell viability measured using PI staining. **(C)** Lipid peroxidation measured using C11 Bodipy staining.
